# Characterizing grapevine 3D inflorescence architecture using X-ray imaging and advanced morphometrics: implications for understanding cluster density

**DOI:** 10.1101/557819

**Authors:** Mao Li, Laura L. Klein, Keith E. Duncan, Ni Jiang, Daniel H. Chitwood, Jason Londo, Allison J. Miller, Christopher N. Topp

## Abstract

Inflorescence architecture provides the scaffold on which flowers and fruits develop, and consequently is a primary trait under investigation in many crop systems. Yet the challenge remains to analyze these complex 3D branching structures with appropriate tools. High information content data sets are required to represent the actual structure and facilitate full analysis of both the geometric and topological features relevant to phenotypic variation in order to clarify evolutionary and developmental inflorescence patterns. We combined advanced imaging (X-ray tomography) and computational approaches (topological and geometric data analysis and structural simulations) to comprehensively characterize grapevine inflorescence architecture (the rachis and all branches without berries) among 10 wild *Vitis* species. Clustering and correlation analyses revealed unexpected relationships, for example pedicel branch angles were largely independent of other traits. We identified multivariate traits that typified species, which allowed us to classify species with 78.3% accuracy, versus 10% by chance. Twelve traits had strong signals across phylogenetic clades, providing insight into the evolution of inflorescence architecture. We provide an advanced framework to quantify 3D inflorescence and other branched plant structures that can be used to tease apart subtle, heritable features for a better understanding of genetic and environmental effects on plant phenotypes.

## Introduction

Inflorescences are major adaptations of the angiosperm lineage whose architectural variation affects fertilization, fruit development, dispersal, and crop yield (Wyatt, 1982; Hake, 2008; de Ribou *et al.*, 2013; Kirchoff & Claßen-Bockhoff, 2013; Périlleux *et al.*, 2014; Chanderbali *et al.*, 2016). These branched reproductive structures with multiple flowers reflect the extraordinary diversity across angiosperm species, from an ear of corn to palms with inflorescences measuring five meters long (Hodel *et al*., 2015). Yet seemingly simple processes give rise to these vastly different shapes - during development reproductive meristems may either switch to floral identity or proliferate additional inflorescence meristems and branches (Prusinkiewicz *et al.*, 2007). Complex topologies reflect the evolution of this functional diversity, but have proven difficult to quantify with conventional tools.

Detailed descriptions of inflorescences by trained experts are often unique to specific research communities or groups of taxa, and are not always readily transferable, hindering meaningful comparative analysis (Endress, 2010). Inflorescences are sometimes described typologically: indeterminate or determinate, simple or compound, as a raceme, cyme, panicle or spike, etc. (Wyatt, 1982; Weberling, 1992). Other approaches describe qualitative attributes of inflorescences such as the presence or absence of certain structures (Weberling, 1992; Doebley *et al.*, 1997; Feng *et al.*, 2011; Hertweck & Pires, 2014). A third method for characterizing inflorescences is through quantification of component structures (e.g., branch length, inflorescence length and width, angular traits; Kuijt, 1981; Marguerit *et al.*, 2009; Landrein *et al.*, 2012; Le *et al.*, 2018). Although these classical quantitative approaches facilitate comparative statistical analyses, the three-dimensional (3D) complexity of inflorescences is largely undescribed. Furthermore, descriptions may be confounded by developmental stage at the time of measurement, and distinguishing between vegetative and reproductive branching structures can be difficult (Wyatt, 1982; Weberling, 1992; Guédon *et al.*, 2001). Thus, new technological and analytical approaches that can represent comprehensive, multi-dimensional information about inflorescence diversity are needed to normalize and enrich analysis of these structures.

One promising approach for capturing 3D shapes of inflorescences and other plant structures is X-ray tomography (XRT). XRT generates high quality reconstructions of the internal and external shapes of plants, preserving nearly complete geometric and topological information in 3D. These 3D digital models then can be used to extract quantitative data (features) from plant structures. X-rays have been used to quantify wheat and rice seed and inflorescence traits from intact samples for non-destructive yield calculations (Hughes *et al.*, 2017; Jhala & Thaker, 2015), internal anatomy of willow trees (Brereton *et al.*, 2015), stem morphology and anatomy in sorghum (Gomez *et al.*, 2018), root structure of barley seedlings (Pfeifer *et al.*, 2015), leaf anatomy in monocots and dicots (Mathers *et al.*, 2018) and dynamic starch accumulation in living grapevine stems (Earles *et al.*, 2018), among others. Most critically, whereas manual measurements can be laborious and destructive, non-destructive sampling for XRT analysis facilitates comprehensive quantification of complex morphological traits.

Quantifying complex shapes with XRT requires appropriate analytical approaches. Topological modeling, a mathematical field concerned with the connectedness of branching structures, can quantify inflorescence architecture by parsing geometric 3D structures into distinct, yet connected, components (Godin & Caraglio, 1998). Topological modeling has yielded important insights into inflorescence development, functional analysis, and crop improvement in a variety of plant species (e.g., *Arabidopsis thaliana*, *Capsicum annuum, Malus pumila*, and *Triticum*; Godin *et al.*, 1999; Letort *et al.*, 2006; Kang *et al.*, 2009). While powerful, these reductionist approaches rely on an a priori understanding of the mechanisms that contribute to complexity (e.g., branching patterns), and lose power when shapes vary drastically from one another (e.g., comparing a corn tassel to a grape cluster). Approaches that capture emergent properties of complex structures without presupposing the importance of individual structural components are complementary to traditional topological models (Bucksch *et al.*, 2017).

An emerging mathematical approach to interpret topological models is persistent homology (PH). PH extracts morphological features from two- or three-dimensional representations and can be used to compare very different shapes. PH has been applied to explain a wide range of features including atomic structures, urban and forested areas, cancers, cell shapes, and jaw shape, among others (Edelsbrunner & Morozov, 2013). In plants, PH has been used to estimate shapes that are otherwise difficult to measure including leaves, leaflet serration, spikelet shape, stomatal patterning, and root architecture (Li *et al.*, 2018a,b; Haus *et al.*, 2018; McAllister *et al.*, 2019; Migicovsky *et al.* 2018). Previous work showed that PH could capture more quantitative variation than traditional plant morphological measures (described above) resulting in the identification of otherwise latent quantitative trait loci (Li *et al.*, 2018b). PH is especially well-suited for quantifying branching topology as it can quantitatively summarize complex variation with a single measure (Li *et al.*, 2017; Delory *et al.*, 2018). Rachis, pedicel, and branches include inherently topological features that can be especially well-analyzed with PH-based methods.

Grape clusters (or bunches) are branched structures supporting berries produced by grapevines (*Vitis* spp.) and are an ideal system in which to apply XRT and PH. Grape infructescences are historically, culturally, and economically important and vary extensively in nature and in cultivation (Iland *et al.*, 2011). Cluster architecture determines bunch density, and is defined as “arrangement of berries in a cluster and the distribution of free space” (Richter *et al.*, 2018). The density of berries in a cluster is an important breeding feature because it determines yield, wine character, and disease resistance (amount of air flow between berries is a primary determinant of pests and pathogens on the fruit). Cluster density is a characteristic identified by the Organization Internationale de la Vigne et du Vin, and varies from “berries clearly separated” (loose clusters) to “berries deformed by compression” (very dense clusters; OIV, 2001). As one of the primary determinants of yield, end-product characteristics, and disease resistance cluster architecture has been studied extensively in grapevine (reviewed in Tello & Ibáñez, 2018). These studies have shown that wine grape cultivars (*Vitis vinifera*) display distinct bunch densities (Shavrukov *et al.*, 2004). However, less is known about cluster architecture in wild *Vitis* species, an important source of natural variation used by breeders in the development of hybrid grapevine varieties.

Historically, researchers have focused on a suite of cluster traits such as cluster size, shape, weight, and density/compactness to characterize bunch density quantified in grapevines (Rovasenda, 1881; Pulliat, 1888; Bioletti, 1938; Galet, 1979; Bettiga, 2003). Measurements are made primarily using traditional tools including rulers, digital calipers, volume displacement, and/or through human judging panels. More recently, automated image-based approaches have been implemented to capture aspects of cluster architecture in the lab and field (Ivorra *et al.*, 2015; Aquino *et al.*, 2017, 2018; Rist *et al.*, 2018). However, these image-based methods cannot penetrate the internal inflorescence structure. Therefore resulting models are based only the visible surface and the underlying topology cannot be fully captured, limiting an understanding of how inflorescence architecture and berry features co-vary. XRT and PH applications offer an important opportunity to understand grapevine bunch density through detailed analyses of inflorescence architecture. This work will deepen our understanding of natural variation of inflorescence structure, identify priority targets for breeding, and permit connecting 3D structure to underlying processes and genetics of inflorescence development.

We use X-ray tomography, geometric measurements, persistent homology, and structural simulation to characterize wild grapevine inflorescence architecture. We target the branching architecture of the mature inflorescence: the rachis and all branches that remain following the removal of ripe berries (Fig. 1). Specifically, we aim to: 1) characterize variation in component traits of inflorescence architecture within and among *Vitis* species; 2) assess phylogenetic signals underlying inflorescence architecture traits; and 3) interpret inflorescence trait variation in the context of breeding objectives. This work represents an important advance for the characterization of 3D plant architecture using a powerful combined imaging and computational approach.

**Fig. 1.**
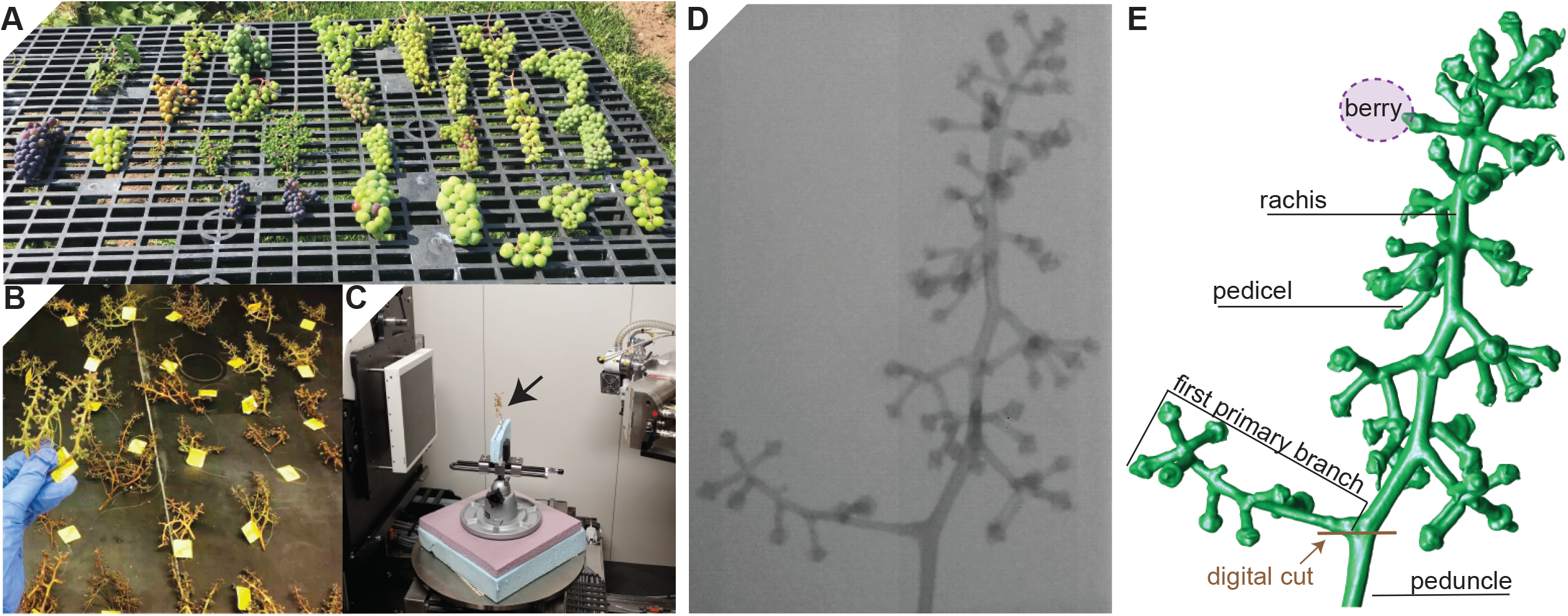
Sample preparation and imaging. (A) The ten *Vitis* species sampled for this study display diverse grape bunch morphology. (B) Inflorescence architectures after berry removal. (C) Inside the X-ray tomography instrument; the inflorescence is clamped in a panavise between two pieces of polystyrene on the X-ray turntable. (D) Two dimensional radiogram of grape inflorescence; X-rays, absorbed or passing through the inflorescence, are detected to create a silhouette. (E) Three dimensional reconstruction and the structure of the same inflorescence shown in (D) by taking radiograms at successive different angles and then computationally combining the images.

## Materials and methods

### Plant Material

In this study, we sampled grapevine bunches from 136 unique genotypes representing 10 wild *Vitis* species living in the USDA germplasm repository system (Geneva, NY; Table 1, Supplementary Fig. S1). Grapevines have a paniculate inflorescence that consists of a rachis with several primary and secondary branches, tapering towards the terminus of the organ (Iland *et al.*, 2011). Wild grapevines are dioecious; consequently, unbalanced sample sizes for different species reflect numbers of female genotypes available in the germplasm collection. Each unique genotype is represented in the germplasm collection by two clonally replicated vines. For most of the 136 genotypes, we collected a total of three clusters from the two clonal replicates combined, representing average cluster morphology. We avoided clusters that were visibly damaged or indirectly altered (e.g., tendril or trellis interference). For each vine, clusters were removed from separate canes at the point of peduncle attachment (Fig. 1A). In total, 392 clusters were collected in September 2016 when berries were soft, equivalent to EL38 developmental stage (Coombe, 1995; Fig. 1B). Berries were manually removed from clusters in the field, and the remaining inflorescence stalks (including rachis, branches, and pedicels; hereafter referred to as inflorescence or inflorescence architecture) were used to assess inflorescence architecture.

**Table 1.** Number of samples/individuals each species and berry information used in the study.

|  | Number (N) |  |  | Berry information (Galet (1988); Moore and Wen (2016)) |  |  |
| --- | --- | --- | --- | --- | --- | --- |
|  | Samples | Individuals | Individuals used in phylogenetic analysis | low diameter (mm) | High diameter (mm) | Berries per bunch |
| *V. acerifolia* | 32 | 11 | 9 | 8 | 12 | >25 |
| *V. aestivalis* | 5 | 2 | 1 | 8 | 20 | >25 |
| *V. amurensis* | 13 | 5 | 2 | 8 | 15 | NA |
| *V. cinerea* | 45 | 15 | 13 | 4 | 8 | >25 |
| *V. coignetiae* | 6 | 2 | 1 | NA | 8 | NA |
| *V. labrusca* | 62 | 22 | 12 | 12 | 23 | <25 |
| *V. palmata* | 3 | 1 | 1 | 8 | 10 | >25 |
| *V. riparia* | 158 | 53 | 48 | 8 | 12 | >25 |
| *V. rupestris* | 41 | 16 | 10 | 8 | 12 | <25 |
| *V. vulpina* | 27 | 9 | 2 | 8 | 12 | >25 |
| **Total** | 392 | 136 | 99 |  |  |  |

### X-ray tomography and data preprocessing

Grapevine inflorescences were scanned at the Donald Danforth Plant Science Center (St. Louis, MO) using a North Star Imaging X5000 X-ray tomography instrument (NSI; Rogers, MN) equipped with a 16-bit Varian flat panel detector (1536 × 1920 pixels with 127um pixel pitch) and 225kV microfocus reflection target X-ray source. Each inflorescence was held between two pieces of construction-grade expanded polystyrene, clamped in a panavise, and positioned on the X-ray turntable in one of two configurations (Fig. 1C): 725mm from the source, generating 1.26x magnification and 101um voxel resolution, or 766mm from the source, generating 1.19x magnification and 107um voxel resolution. Each scan used X-ray wattage set to 60kV and 1200uA at 10 frames per second, collecting 1200 16-bit TIFF projections over 360 degrees of rotation during a 2min continuous standard scan. Projections for each scan (Fig. 1D) were combined into a single 3D volume using NSI efX-CT software, converted to a density-based surface rendering Polygon file (PLY), and exported for analysis (Fig. 1E). The full PLY data set for this work is 7.85GB, and can be downloaded from: https://www.danforthcenter.org/scientists-research/principal-investigators/chris-topp/resources.

We exported the surface mesh data (.ply files) into Meshlab (v1.3.3, (Cignoni *et al.*, 2008) and performed the following processing steps to remove topological noise: 1) deleted the vertices where branches touch using “Select Vertexes” and “Delete Selected vertices” filters; 2) removed duplicates and isolated vertices and faces using the filters “Remove Duplicated Vertex,” “Remove Duplicate Faces,” “Remove Isolated pieces (wrt Diameter),” and “Remove Unreferenced Vertex.”

### Geometric inflorescence architecture traits

We extracted 15 geometric traits from scanned inflorescences (Fig. 2, Supplementary Fig. S2). Detailed trait descriptions and calculations are explained in Supplementary Table S1. Trait illustrations, including examples of low and high values for each trait, are available in Fig. 2 and Supplementary Fig. S2. Traits were organized in one of three trait groups: global-size features, local-branching features, and size-invariant features (Table 2). PedicelDiameter and PedicelBranchAngle were measured using the software DynamicRoots (Symonova et al. 2015) on a subset of detected pedicels from the raw 3D volume data. All other traits were derived from Matlab algorithms. Branch length traits (i.e., TotalBranchLength, RachisLength, PedicelLength, and AvgBranchLength) were derived from the persistence barcode (see next subsection).

**Table 2.**
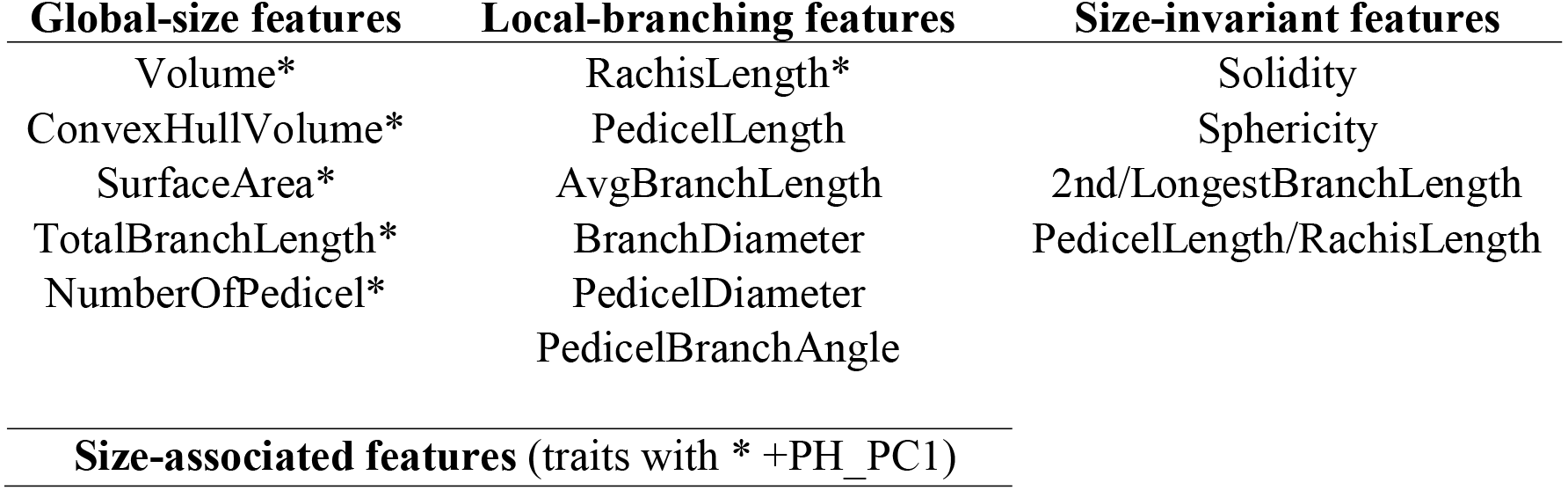
Fifteen geometric traits were organized into three categories based on the type of shape information captured by the trait. See Table S1 for a more detailed description of each trait.

**Fig. 2.**
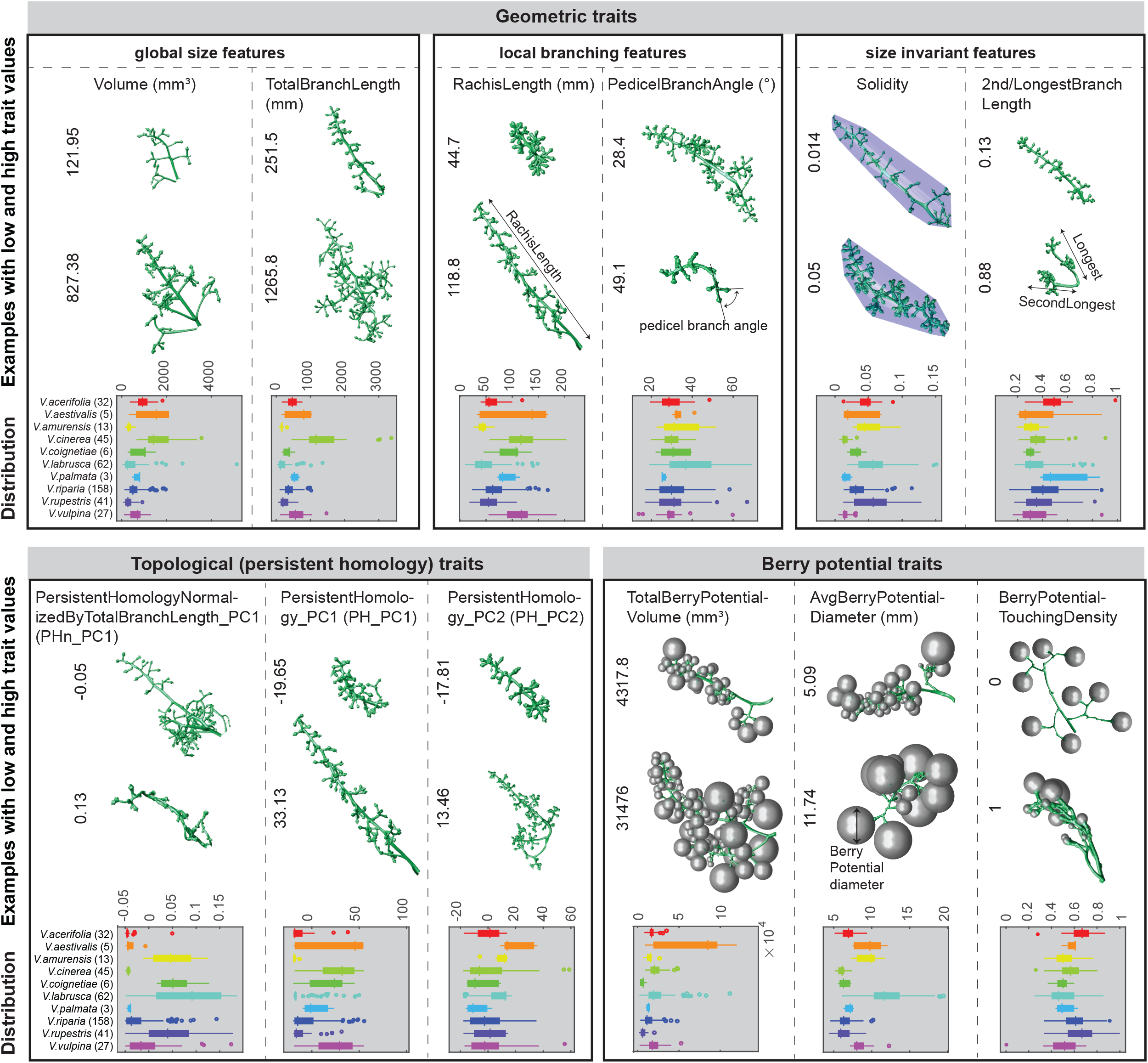
Examples of inflorescence geometric and topological traits and their distribution for ten *Vitis* species. Each panel shows one of the three traits categories (geometric traits, topological traits, and berry potential traits). Geometric traits are organized as global size features, local branching features, and size-invariant features. Each trait is listed at the top of the column and two inflorescence examples demonstrating low and high trait values listed to the left. At the bottom of each column is a boxplot indicating the distribution and variance within the ten *Vitis* species, represented in different colors. On each box, each dot indicates an outlier if it is more than 1.5 interquartile ranges; the central vertical line indicates the median; the left and right edges of the box represent the 25th and 75th percentiles; and the whiskers extend to the most extreme nonoutlier data. The label for each species is listed in the boxplot y axis of the leftmost plot, with the number of individuals sampled for each species shown in parentheses. For a more complete example and detailed description of each trait, see Fig. S2 and Table S1.

### Quantifying branching topology using persistent homology, a topological data analysis method

Persistent homology measures shapes based on a tailored mathematical function, such as geodesic distance, which we used here to capture both curved length and topology of the branches (Fig. 3, Supplementary Video S1). The geodesic distance of a point is the length of the shortest curve connecting the point and the base (e.g. purple curves, Fig. 3A), where the tailored base can be set as the first node or ground level (the brown line in Fig. 3A). For each branch, the tip always has the largest geodesic distance from the base (Fig. 3B). A level represents the collection of points whose geodesic distances are the same (e.g. geodesic distance=90, pink curve in Fig. 3A). A superlevel set, for example, at 90, is all the points whose geodesic distances are greater than 90 (black branch tips, Fig. 3A). Changing the level value from largest to smallest (x axis, Fig. 3C), the sequence of nesting superlevel sets can be formed, which is named superlevel set filtration (top panel, Fig. 3C). During the change of the level value, bars record the connected components for each of the superlevel sets. When a new component arises, a new bar starts (e.g. at level 112, purple branch, Fig. 3C). When two components merge (e.g. at level 65, orange branch merges into purple branch, Fig. 3C), the shorter bar stops (e.g. the orange bar stops at level 65, Fig. 3C). This bar graph, called the persistence barcode, summarizes topological information such as branching hierarchy, branch arrangement, and branch lengths. In our study, we set the base as the junction between peduncle and rachis (the lowermost node, indicated by a brown line in Fig. 1E, Fig. 3D, F) and use this base to compute the persistence barcode for the inflorescence architecture (Fig. 3E, G).

**Fig. 3.**
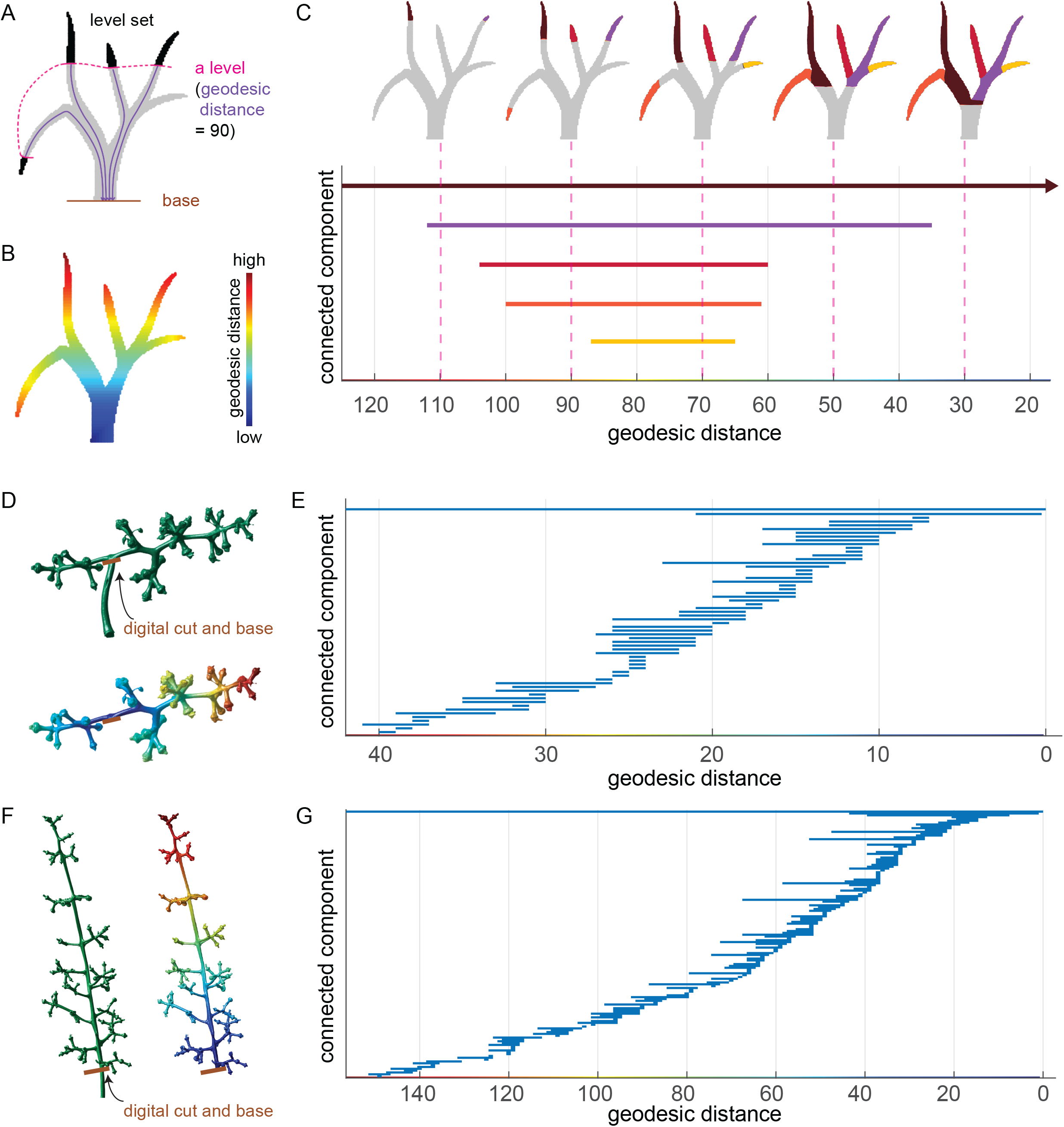
Persistent homology with geodesic distance comprehensively quantifies branching structures. (A) A level (pink solid line) defined by the same geodesic distance (length of any of the purple curves, in this case, set to 90) to the base of the inflorescence. The super level set is the pixels (in black) having greater geodesic distance than the pink level. (B) Pixels on a branching structure are colored by their geodesic distance to the base. They are colored with red representing the most distant through to blue for the closest ones. (C) A persistence barcode for each branching structure records the connected components for each level set at each geodesic distance value. The “birth” and “death” values for each bar represent the level where each branch starts and gets merged. Colored bars correspond to colored branches. (D) Above: example inflorescence. The stem is digitally cut at the base (brown line) where it meets the first branch. Below: 3D surface on the example inflorescence as in (B). (E) Persistence barcode for the inflorescence in (D). (F) and (G), similar to (D) and (E), show a different inflorescence architecture.

The persistence barcode can be used to compare topological similarity between any two inflorescences. To compute pairwise distance among persistence barcodes for the entire inflorescence population, we used the bottleneck distance (Cohen-Steiner *et al.*, 2007). Bottleneck distance is a robust metric that calculates the minimal cost to move bars from one persistence barcode to resemble another (Li *et al.*, 2017). We performed multidimensional scaling (MDS) on the pairwise bottleneck distance matrix and projected the data into lower dimensional Euclidean space by preserving the pairwise distance as well as possible. The Matlab (R2017a) MDS function cmdscale() projects the data so that MD1 acts as PC1 representing the most variation. The first three PCs (MDs) explained about 80% of the total variation and were included as traits: PersistentHomology_PC1 (PH_PC1, explained about 54% variation), PersistentHomology_PC2 (PH_PC2, explained about 20% variation), and PersistentHomology_PC3 (PH_PC3, explained about 6% variation). Those traits not only measure the topological structure, but also relate to geometric variation (e.g. global size) as the data were not normalized (Fig. 2, Supplementary Table S1).

Next, we normalized the persistence barcode by the TotalBranchLength (summation of the bar lengths) so that the TotalBranchLength was 1. By a similar procedure, we derived the first three PCs named PersistentHomologyNormalizedByTotalBranchLength_PC1 (PHn_PC1, explained about 45% variation), PersistentHomologyNormalizedByTotalBranchLength_PC2, (PHn_PC2, explained about 21% variation), and PersistentHomologyNormalizedByTotalBranchLength_PC3 (PHn_PC3, explained about 7% variation) for the normalized inflorescence topological structure (Fig. 2, Supplementary Table S1).

### Berry potential, an approach to indirectly explore the space limited by inflorescence architecture

An ongoing question in grapevine cluster architecture is the relationship between inflorescence architecture and berry number and size. Inflorescence architecture is one of several factors determining the number of berries that can form, due to the number of pedicels and the available space for berry development. In this study, berries were removed because of concerns about berry integrity during transport from New York to Missouri, and the time between harvest and scanning. Instead of looking directly at berries on the cluster, we used inflorescence architecture as a starting point to simulate potential space available for berry growth by evaluating expanding spheres attached to pedicels. The extent of sphere expansion allowed by each pedicel is referred to as “berry potential” (Fig. 4, Supplementary Video S2).

**Fig. 4.**
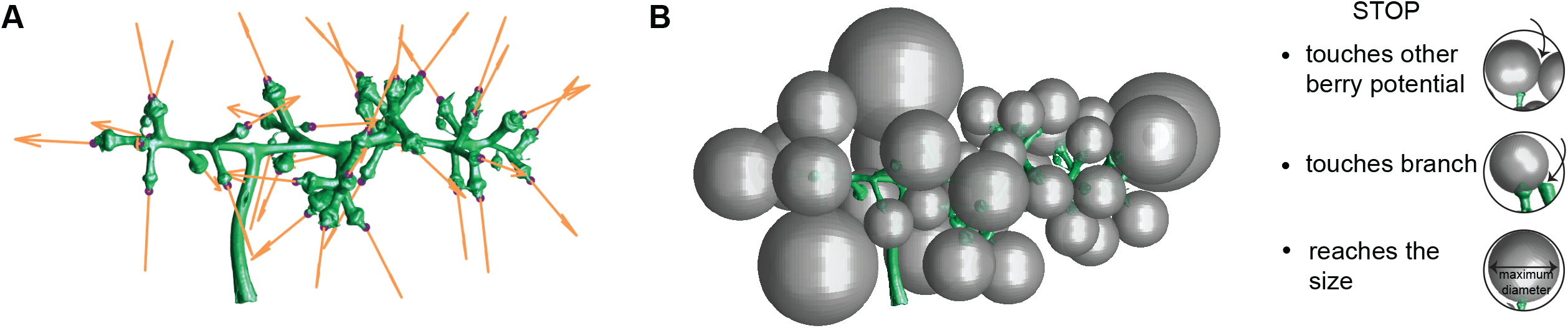
Berry potential simulation to explore the space determined by inflorescence architecture. (A) Determine the growth direction for each berry potential. (B) Expand berry potential by increasing the size and moving the center along the growth direction until it meets any of these three cases: 1) two berry potentials touch each other; 2) a berry potential touches any part of the inflorescence; 3) the diameter of the berry potential reaches the maximum for the species..

We first determined the growth direction for each berry potential based on the pedicel orientation. When spheres expand, the center moves along the pedicel direction (Fig. 4A). This step can be achieved by performing principal component analysis (PCA) on the near-berry segment of the pedicel. The first principal axis is the pedicel direction. We adjusted the arrow of the direction to make sure berry potential increases outward along the pedicel orientation. Then the berry potential increases until one of three situations is encountered (Fig. 4B): 1) if two berry potentials touch to each other, both berry potentials will stop increasing; 2) if a berry potential touches any part of the inflorescence, it will stop increasing; 3) if the diameter of the berry potential reaches the maximum size known for that species (Table 1), it will stop increasing. For each species, the maximum size is defined as the maximum berry diameter, a number estimated from known ranges of berry sizes for each species, based on values obtained from (Galet, 1988; Moore & Wen, 2016).

Berry potential does not reflect true berry growth; rather, berry potential is a derived attribute of inflorescence architecture, an indirect estimate of the space potentially available for berry growth. It also does not account for the possibility of branches bending or otherwise becoming re-oriented due to pressure from growing berries. Berry potential is based on the number of neighbor pedicels, neighbor pedicel lengths, and neighbor pedicel mutual angles. Larger values for berry potential are associated with fewer neighbor pedicels, and/or longer pedicel lengths, and/or larger mutual angles. From the berry potential simulation, we calculated three features, TotalBerryPotentialVolume, AvgBerryPotentialDiameter, and BerryPotentialTouchingDensity, which is the berry potential touching number (i.e., touching either another berry potential or any part of the inflorescence) divided by the number of berry potential (Fig. 2, Supplementary Table S1).

### Phylogenetic analysis

Phylogenetic analyses were conducted to understand evolutionary trends in inflorescence architecture in *Vitis*. Single nucleotide polymorphism (SNP) markers were generated as part of a separate study of the USDA Grapevine Germplasm Reserve in Geneva, NY (Klein *et al.*, 2018). The original dataset consisted of 304 individuals representing 19 species that were sequenced using genotyping-by-sequencing (GBS; Elshire *et al.*, 2011). Briefly, Klein *et al.* (2018) filtered data to retain biallelic sites with a minimum allele frequency of 0.01, a minimum mean depth of coverage of 10x, and only sites with <20% missing data and individuals with <20% missing data. SNP data for 99 individuals from this study that were also genotyped in (Klein *et al.*, 2018); Table 1) were extracted using custom scripts. We performed phylogenetic analysis on the sequence data extracted for 99 individuals using SVDquartets (Chifman & Kubatko, 2014), a maximum likelihood approach designed to address ascertainment bias associated with reduced representation sequencing techniques like GBS. We analyzed all possible quartets and carried out 100 bootstrap support runs (Supplementary Fig. S1) using PAUP* version 4.0a (Swofford, 2003). The three main clades recovered in the tree were consistent with previous phylogenetic work in *Vitis*: 1) an Asian Clade (*V. amurensis and V. coignetiae*), 2) North American Clade I (*V. riparia, V. acerifolia, and V. rupestris*), and 3) North American Clade II (*V. vulpina, V. cinerea, V. aestivalis, V. labrusca, and V. palmata*) (Tröndle *et al.*, 2010; Zecca *et al.*, 2012; Miller *et al.*, 2013; Zhang *et al.*, 2015; Klein *et al.*, 2018).

To visualize trait distributions on a phylogenetic tree using branch lengths, we used Mega X (Kumar *et al.*, 2018) to generate a neighbor joining tree with 2000 bootstrap replicates. All measurements were averaged across the three replicates per genotype to produce an average value for each trait for each genotype. We computed Pagel’s lambda to estimate phylogenetic signal for each morphological trait and mapped each trait onto the phylogeny (Supplementary Fig. S3A-X) using the R package phytools (v. 0.6-44; Revell, 2012). We calculated variation of each morphological trait for each clade based on the mean value for each species (Supplementary Fig. S4).

### Statistical analysis

PCA, MDS, and hierarchical cluster analysis generating a hierarchical tree were performed in Matlab using functions pca(), cmdscale(), and clustergram(). The R function cor.mtest() and package corrplot (Wei & Simko, 2017) were used for significance tests and correlation matrix visualization. The function lda() in R package MASS (Venables & Ripley, 2002) was used for the linear discriminant analysis (LDA) with a jackknifed ‘leave one out’ cross validation method.

### Code availability

All Matlab functions used to calculate persistence barcodes, bottleneck distances, simulation for berry potential, other geometric features used in this study, and the script for extracting phylogenetic information can be found at the following GitHub repository: https://github.com/Topp-Roots-Lab/Grapevine-inflorescence-architecture.

## Results

### Inflorescence morphological variation and trait correlation within *Vitis* species

We investigated 24 morphological traits (15 geometric traits, six PH traits, and three berry potential traits) of inflorescence architecture in 10 wild *Vitis* species (136 genotypes, 392 samples) and detected wide variation in morphological features within and between species (Fig. 2, Supplementary Fig. S2 and Table S2). In particular, of all the species examined, *V. aestivalis* has the largest variance for TotalBerryPotentialVolume. *V. labrusca* has the largest variance for ten traits (i.e., pedicel features, Sphericity, AvgBranchDiameter, AvgBerryPotentialDiameter, and normalized topological traits). *V. cinerea* has the largest variance for six traits (i.e., most global-size features, PH_PC2, and PH_PC3). In comparison, *V. palmata* has smallest variance for eight traits (i.e. pedicel features, Sphericity, AvgBranchDiameter, TotalBerryPotentialVolume, PH_PC3, and PHn_PC3), as does *V. amurensis* (global-size features, RachisLength, PH_PC1, and PH_PC2).

All traits were hierarchically clustered based on the mean trait values for each species, classifying traits into two main categories: mostly size-invariant + local-branching features (PHn_PC3 to PedicelLength), versus global-size features (AvgBranchLength to BerryPotentialTouchingDensity) (Fig. 5A). Hierarchical clustering (Fig. 5A) and pairwise correlation for morphological traits (Fig. 5B) show that global-size features (ConvexHullVolume, SurfaceArea, Volume, NumberOfPedicel, and TotalBranchLength), PH_PC1, and RachisLength are all highly positively correlated. We refer to these seven traits as size-associated features. Size-associated features are negatively correlated with PedicelLength/RachisLength, Solidity, Sphericity, and PHn_PC1. Some traits are relatively independent such as 2nd/LongestBranchLength, PedicelLength, PedicelBranchAngle, PH_PC2, PHn_PC2, and PHn_PC3 (Fig. 5B). PH_PC3 has some negative relation with size-invariant features. PHn_PC1 positively correlates with Sphericity, Solidity, and AvgeBerryPotentialDiameter (Fig. 5B). Pairwise correlations of morphological features (allometric relationships) for each of the species vary widely (Fig. 5C; for all traits see Supplementary Fig. S5A-X). For example, more pedicels typically result in smaller berry potential diameters, except for *V. aestivalis*. Longer branches tend to be thinner, except for *V. coignetiae,* and correlate with larger inflorescences, except in *V. acerifolia*.

**Fig. 5.**
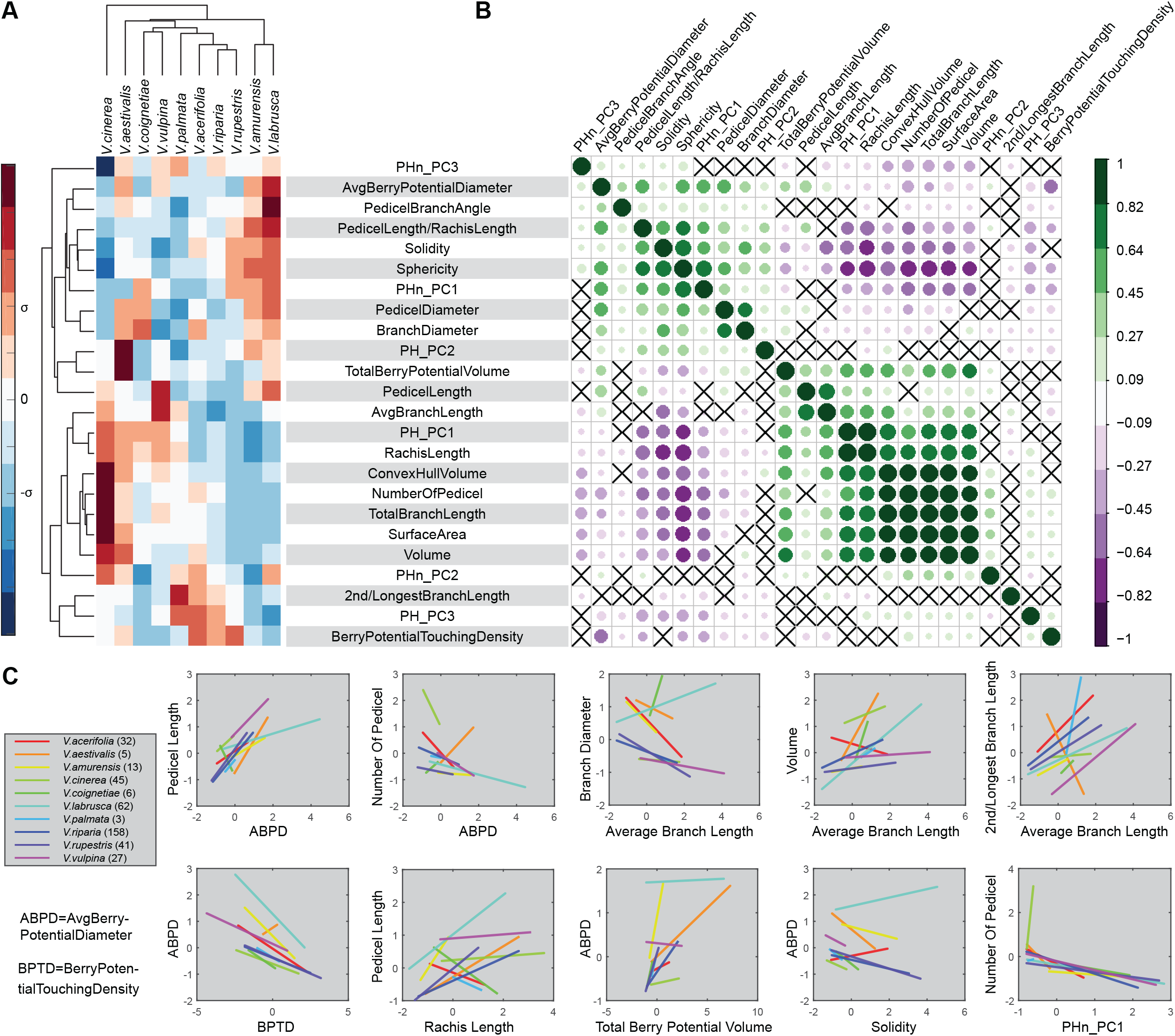
Hierarchical cluster analysis and correlation analysis. (A) Cluster analysis based the mean value for each trait of 10 *Vitis* species. The heatmap shows values above (red) or below (blue) the mean for each trait. The morphological traits (rows) are clustered hierarchically with the name shown on the right and hierarchical tree listed on the left. The species (columns) are also clustered hierarchically with the name and hierarchical tree shown at the top. (B) Correlation matrix plot shows pairwise positively stronger correlation (green and larger circle) or negatively stronger correlation (purple and larger circle). Non-significant correlations (p>0.05) are crossed out. The traits are ordered in the same way as (A). (C) Selected pairs of traits showing linear regression lines for each species.

Hierarchical clustering of 10 *Vitis* species based on the 24 morphological traits resolved four groups: 1) *V. cinerea*, 2) *V. aestivalis,* 3) *V. coignetiae/ V. vulpina/ V. palmata/ V. acerifolia/ V. riparia/ V. rupestris,* and 4) *V. amurensis/ V. labrusca* (Fig. 5A). Among the 10 *Vitis* species examined in this study, the largest variance in mean trait values are seen in *V. cinerea* (Fig. 5A). *V. cinerea* samples are generally larger than those from the other species, as reflected in size-associated traits. Topology traits such as PHn_PC3 and size-invariant traits like Sphericity and Solidity are lower in the mean trait value for *V. cinerea* than for other species. Similarly, mean trait values are larger for size-associated traits in *V. aestivalis* (Fig. 5A). Compared to other species, topology and berry potential traits are larger in *V. aestivalis*. Mean trait values of the third group (*V. coignetiae/ V. vulpina/ V. palmata/ V. acerifolia/ V. riparia/ V. rupestris*, Fig. 5A) tend to be nearer to middle values compared to the other species. Within this group, *V. acerifolia/ V. riparia/ V. rupestris* typically are larger in the mean trait value for berry potential touching (i.e., denser berry potentials). These three species and *V. palmata* tend to have large, first primary branches (i.e., wings; Fig. 1E). *V. coignetiae* has thicker branches and *V. vulpina* has longer pedicels compared to other species in this group. The final group, *V. amurensis* and *V. labrusca,* have relatively smaller inflorescences with thicker branches compared to the other species sampled here. These general features are reflected in larger mean values for several size-invariant and local-branching features and smaller mean values for many branch length dependent and size-associated features, respectively (Fig. 5A).

### Multivariate, discriminant analysis of Vitis species based on inflorescence architecture

In order to understand how overall inflorescence architecture varies among *Vitis* species, we performed PCA using all 24 morphological features and all samples. PC1 explained 37.12% of the total variation in the measured architecture (Fig. 6A). The traits with the largest values for PC1 loadings, indicating that they contributed most to variation, are size-associated features, Solidity and Sphericity. PC2 explained 15.4% of the total variation in the measured inflorescence architecture, with variation primarily explained by local-branching features such as PedicalDiameter, PedicelLength, PedicelLength/RachisLength, AvgBranchLength, BranchDiameter, three berry potential traits, and PHn_PC1 (Fig. 6A). Although inflorescences from each species occupy different regions of morphospace, these regions overlap considerably.

**Fig. 6.**
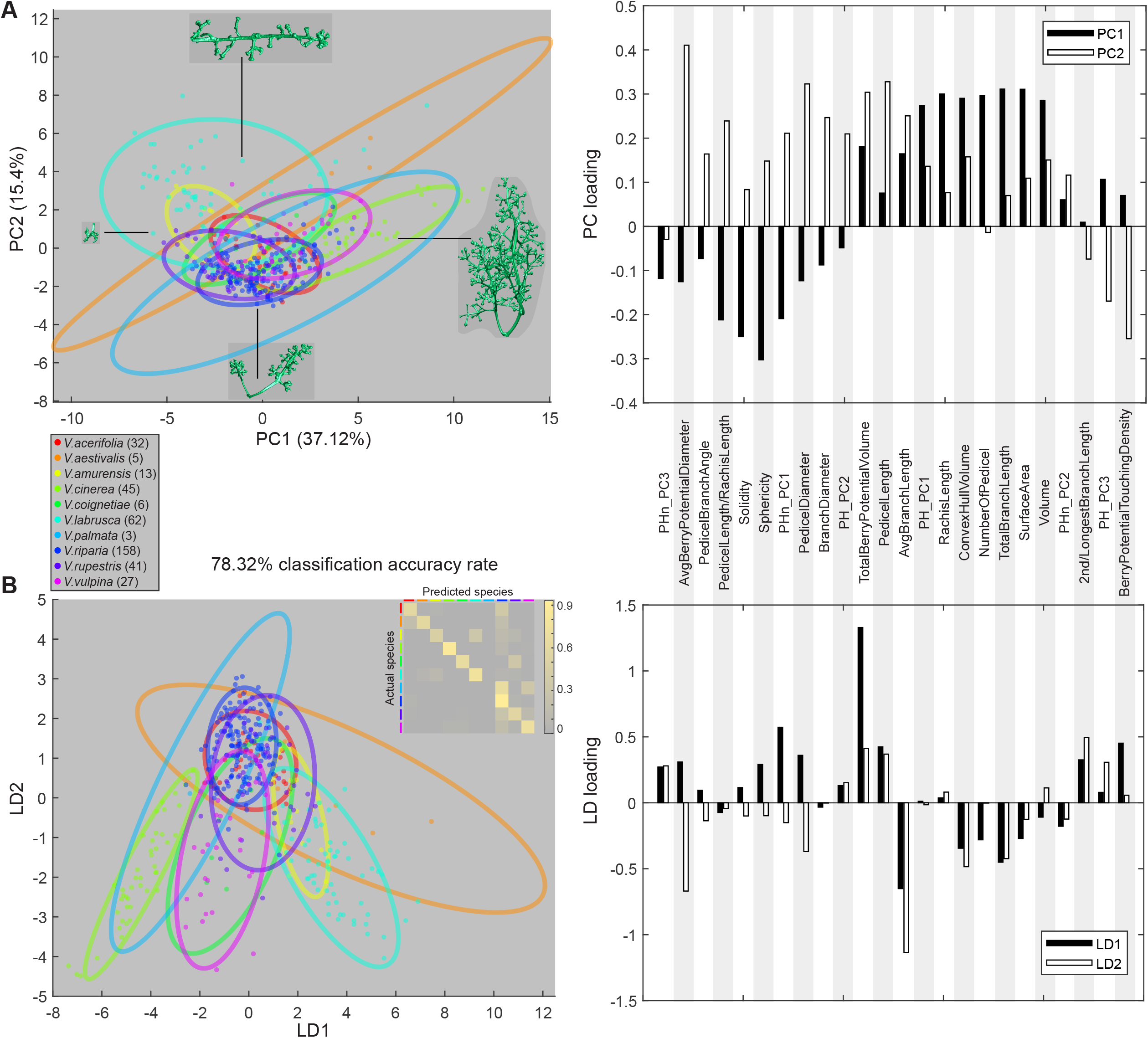
Classification for ten *Vitis* species based on inflorescence architecture. (A) Left: Principal component analysis (PCA) plot on 24 morphological traits. The percent variance for each PC explained is shown in parentheses. Species are shown in different colors. Right: The loadings for the traits that contribute to the variance are shown. (B) Left: Linear discriminant analysis (LDA) plot on the first 18 PCs (99.5% variance). Species are shown in different colors. The confusion matrix for predicted species is shown in the upper right corner. Right: The loadings for the traits that best distinguish species from each other are shown. Using a jacknifed ‘leave one out’ cross validation, we obtain a 78.32% classification accuracy rate.

LDA performed on the first 18 PCs, explaining 99.5% of the variation, distinguished between species with a classification accuracy rate of 78.32%. A confusion matrix (Fig. 6B) shows the proportion of samples correctly predicted for each species. LD1 primarily separates *V. cinerea*, *V. labrusca*, and *V. amurensis* from the other species while LD2 primarily separates *V. vulpina* and *V. coignetiae*. The traits that are most important for distinguishing these species, as indicated by LD loadings, are TotalBerryPotentialVolume and PHn_PC1 for LD1, and AvgBranchLength and AvgBerryPotentialDiameter for LD2 (Fig. 6B). The most important predictors for correctly separating any two species are shown as the grey scaled boxes in Supplementary Fig. S6 and Table S3. For example, BranchDiameter and PedicelDiameter are key when contrasting *V. coignetiae* and *V. vulpina*, suggesting that different branch thickness easily distinguishes these two species. This method correctly determined species classifications with 100% accuracy when contrasting *V. aestivalis* and *V.cinerea, V. aestivalis* and *V.palmata*, *V. aestivalis* and *V. vulpina*, *V. amurensis* and *V. cinerea*, *V. amurensis* and *V. palmata*, *V. cinerea* and *V. coignetiae*. Other combinations of species are harder to distinguish on the basis of inflorescence characters. For example, the classification accuracy rate was only 80% when distinguishing between *V. amurensis* and *V. labrusca* and 82% for *V. aestivalis* and *V. coignetiae*.

### Phylogenetic signal of inflorescence architecture within clades

The phylogeny dataset (N=99) is generally well-supported at the species level and correlates well with current taxonomy. Using average trait values per individual, Pagel’s lambda shows 12 morphological traits (seven size-associated features along with PedicelDiameter, TotalBerryPotentialVolume, Sphericity, PH_PC2, PHn_PC1) have strong phylogenetic signal (lambda>0.8, Fig. 7, Supplementary Table S4). While most species sampled tend to have small values for the seven size-associated features, *V. aestivalis, V. cinerea,* and *V. vulpina* tend to have values that are either close to median, or larger. On average, *V. labrusca* has larger values for Sphericity and PHn_PC1 compared to other species sampled, while *V. cinerea* generally has some of the smallest values for these traits. Only two morphological traits (2nd/LongBranchLength, lambda=0.06 and BerryPotentialTouchingDensity, lambda=0.25) lack phylogenetic signal (Fig.7, Supplementary Table S4).

**Fig. 7.**
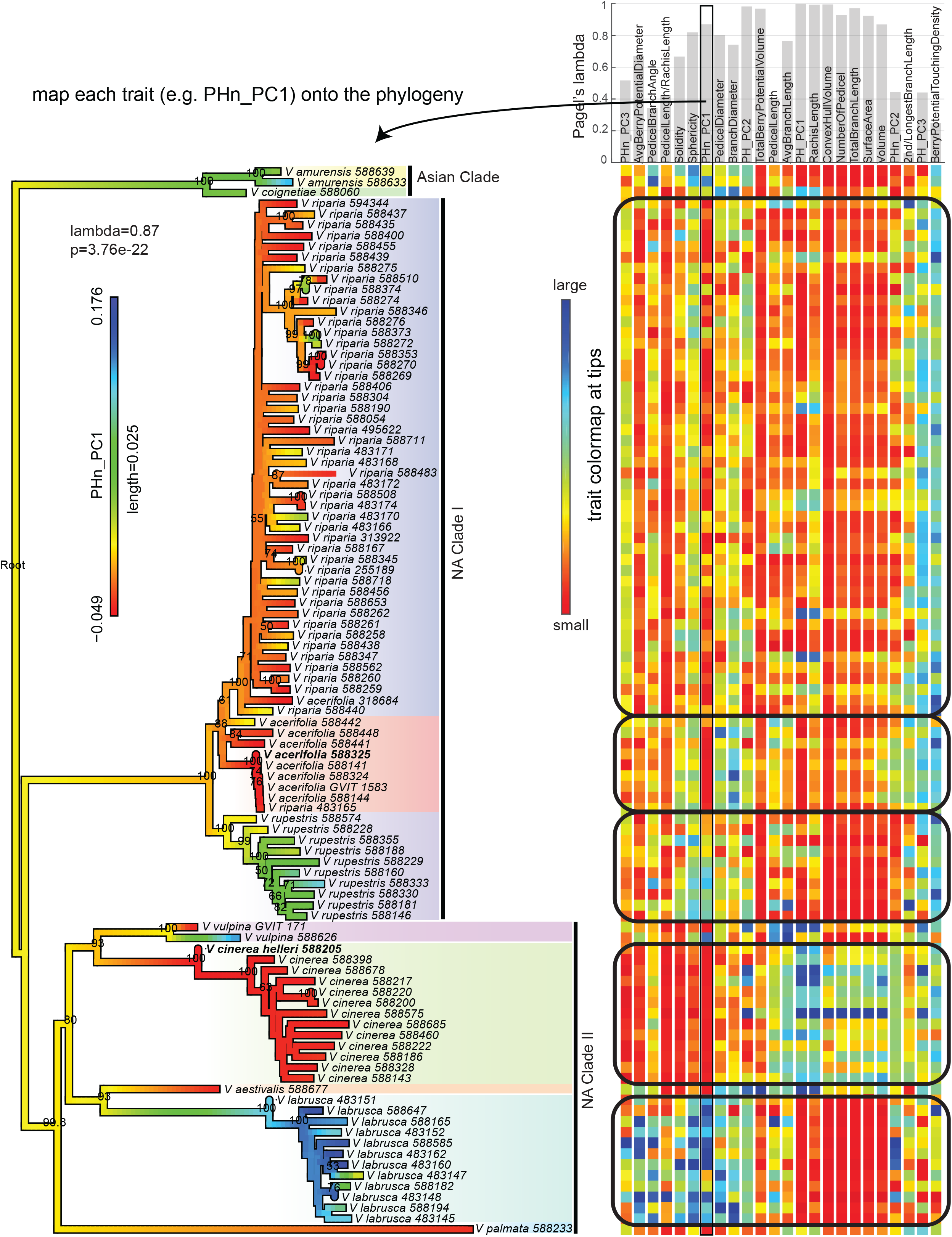
Phylogenetic analysis. A Neighbor Joining phylogenetic tree for a subset of the *Vitis* data set (n=99). Node values denote bootstrap support for values greater than or equal to 50. Ten *Vitis* species are highlighted in different colored backgrounds. Three clades (Asian Clade, NA Clade I, and NA Clade II) are labeled and marked by vertical bars. The barplot showing values of Pagel’s lambda, an estimate of phylogenetic signal, overlaps with the trait name on the right top panel. Below each trait, a rainbow colormap shows the values for individuals (small values in red to large values in blue). Rectangles surround the trait value map for species with more than five individuals. One trait (PHn_PC1) was randomly selected to be projected onto the phylogenetic tree branches, and indicates trait variation (red, lower values; blue, higher values) within individuals and among clades.

We observe differences in *Vitis* inflorescence architecture among clades and between species. For North American (NA) clade I (*V. acerifolia, V. riparia, V. rupestris*), variation in the 24 morphological traits measured have similarly small values among species, particularly for several size-associated traits, although there is relatively large variation for PH_PC3 and BerryPotentialTouchingDensity (Fig. 7). Within NA Clade I, we observe differences among clade members for traits such as Sphericity and PHn_PC1 (larger in *V. rupestris* compared to other clade members) and PedicelDiameter and BranchDiameter (slightly larger in *V. acerifolia* compared to other clade members; Fig. 7). NA Clade II appears to be more variable among clade members. *V. cinerea* has larger values for size-associated traits compared to clade members *V. labrusca*, *V. palmata,* and *V. vulpina*. Meanwhile, *V. labrusca* typically has larger values for local features (e.g., Sphericity, PedicelDiameter, AvgBerryPotentialDiameter, PedicelBranchAngle) compared to the other clade members (Fig. 7).

We calculated the mean value for each species of each morphological trait to study variation within the three clades and detect subtle signatures (Fig. 7). We computed the variance for the multivariate trait (combining all the 24 traits), and each of these 24 traits for each clade (Supplementary Fig. S4, Supplementary Table S5). Overall, based on the samples used in this analysis, variance of the multivariate trait for the NA Clade I (variation=0.14) is much smaller than the NA Clade II (variation=0.64), while the variation for Asian Clade is 0.39. Some traits have almost no variance in Asian Clade such as PedicelDiameter, PHn_PC2, PH_PC3, and 2nd/LongestBranchLength. However, North American species (8/~19 taxa) in this study are better represented than Asian species (2/~37 taxa), so we are cautious not to overinterpret this finding. Traits with the greatest variance in the Asian Clade included PedicelLength/RachisLength, RachisLength, and PH_PC1, while NA Clade I has greatest variance in PHn_PC2. All the other traits have greatest variance in the NA Clade II (Supplementary Fig. S4, Supplementary Table S5). Traits with the smallest variance in the Asian Clade included PHn_PC3, PHn_PC1, PedicelDiameter, BranchDiameter, NumberOfPedicel, 2nd/LongestBranchLength, PH_PC3, and BerryPotentialTouchingDensity. The other traits had small variance in NA Clades I (Supplementary Fig. S4, Supplementary Table S5). Our results highlight clade-specific variation in inflorescence architecture for previously undescribed traits.

## Discussion

Inflorescence architecture provides the scaffold on which flowers and fruits develop, and consequently is a primary trait under investigation in many crop systems. Studies extend into interspecific variation, pollen dispersal, genetic architecture, evolution, regulation, and development of inflorescence structures (e.g., Bradley *et al.*, 1996; Friedman & Harder, 2004; Kellogg, 2007; Morris *et al.*, 2013; Han *et al.*, 2014; Hodge & Kellogg, 2015; Whipple, 2017; Stitzer & Ross-Ibarra, 2018; Ta *et al.*, 2018; Richter *et al.*, 2018). Yet the challenge remains to analyze these complex 3D branching structures with appropriate tools. High resolution data sets are required to represent the actual structure and comprehensive analysis of both the geometric and topological features relevant to phenotypic variation and to clarify evolutionary and developmental inflorescence patterns.

Our results demonstrate the power and potential of X-ray imaging and advanced morphometric analysis for investigating complex 3D phenotypic features. We analyzed the phenotypic variation in inflorescence architecture of 10 wild *Vitis* species using computer vision and an emerging biological shape analysis method, persistent homology, which allowed comprehensive comparisons of shape. Although samples analyzed here represent only a subset of the known variation in *Vitis*, which includes an estimated 60 species, our analyses demonstrate significant variation within and among *Vitis* species and among clades. Correlation analysis (Fig. 5B) revealed some unexpected relationships, for example pedicel branch angles were largely independent of other traits. It also shows that PH is a complementary feature, as it is relatively independent from most geometric features. We were able to assign widely differing architectures to biological species with high accuracy (Fig. 6) from the 24 different morphometric traits surveyed in this study. PH provides an important contribution to this discriminatory power, as does berry potential (Fig. 6B). We observed that traits such as the rachis length, the sum of all branches, the space encompassing the inflorescence architecture (ConvexHullVolume), and PH can be indicative of species and clade (Fig. 7). Our results suggest meaningful, comprehensive information about the inflorescence structure was captured with a single measure (i.e., the persistence barcode) and that PH is a valuable method for quantifying and summarizing topological information.

Persistent homology analysis has led to a deeper understanding of trait genetic variation and architecture in plants. Li *et al.* (2018a) used PH to analyze two-dimensional (2D) leaf shape and predicted family identity with accuracy greater than expected by chance in over 140 plant families, outperforming other widely-used methods of digital shape analysis. Li *et al.* (2018b) showed that PH-based, topological data analysis distinguished between genotypes and identified many new quantitative trait loci (QTL) with 2D tomato leaf shape and root architecture data. This work sets a precedent for measuring observable, yet previously undescribed, phenotypes. In grapevine, QTL analysis indicates a genetic basis to inflorescence architecture and berry compactness (Correa *et al.*, 2014; Richter *et al.*, 2018). Deploying PH-based, topological modeling to grapevine mapping populations could lead to the rapid identification of additional inflorescence trait QTL for breeding. For example, we observed total branch length (a proxy for bigger or smaller clusters) correlates with number of pedicels (a proxy for berry number; Fig. 5), an informative relationship to assess potential yield. However, selecting for total branch length might lead to a negative correlation with the average berry potential diameter (i.e., smaller berries). Although this correlation may be desirable for wine grapes, it is not for table grapes.

Grapevine cluster architecture is a composite feature that reflects multiple subtraits including stalk traits (inflorescence architecture) and berry features (Richter *et al.*, 2018). OIV 204 uses “bunch: density” to describe variation in clusters, ranging from (1) berries clearly separated with many visible pedicels to (9) berries deformed by compression (OIV, 2001; Rombough, 2002). Other authors have deconstructed traits contributing to cluster architecture primarily through individual measurements taken by hand (e.g., Shavrukov *et al.*, 2004; Tello *et al.*, 2015; Zdunić *et al.*, 2015; Tello & Ibáñez, 2018) and more recently, with image-based technologies (Cubero *et al.*, 2014; Roscher *et al.*, 2014; Ivorra *et al.*, 2015; Aquino *et al.*, 2017, 2018; Rist *et al.*, 2018). Here, we are able to describe traits of interest that contribute greatly to the morphological features captured by the OIV scale (e.g., NumberOfPedicel, PedicelLength, PedicelBranchAngle, RachisLength, overall shape using PH; Fig. 2, Supplementary Fig. S2). This method could facilitate precision breeding for both whole inflorescence structure topology and specific desirable geometric traits.

While several studies have quantified cluster structure in cultivated grapevines, similar studies of wild *Vitis* inflorescence architecture are lacking. Munson (1909) and Galet (1979) describe North American *Vitis* cluster structure qualitatively, commenting on compactness, size, shape, and the presence of large first primary branches (wings/shoulders). Taxonomic descriptions typically do not examine inflorescence architecture beyond categorical type, position on the vine, and the average number of berries per cluster (Comeaux *et al.*, 1987; Moore, 1991; Moore & Wen, 2016). Descriptions of the position of the inflorescence are useful for identification and are included in dichotomous keys; however, to our knowledge, other inflorescence architecture traits have not been rigorously quantified among wild *Vitis* species. Although qualitative descriptions are valuable and accessible, powerful phenotyping tools are required to associate complex phenotypes with evolutionary and developmental patterns.

Using 3D imaging and PH with a topological modeling approach, we identified attributes of inflorescence architecture that vary within and among *Vitis* species that, to our knowledge, have not been previously described. Differences in inflorescence architecture among clades mirror other phenotypic differences among members of North American *Vitis*. For example, members of NA Clade I (*V. acerifolia*, *V. riparia*, and *V. rupestris*) have small values for size-associated features (e.g., RachisLength, ConvexHullVolume, NumberOfPedicel, TotalBranchLength, SurfaceArea, Volume) and relatively large values for PH_PC3 and BerryPotentialTouchingDensity (Fig. 7). These species share suites of other morphological characters (nodal diaphragm, branch, and leaf surface traits, and large stipules; Moore 1991, Moore and Wen 2016, Klein *et al.*, 2018). It is possible that among closely related species conserved pathways generate vegetative and reproductive similarities.

Sample size is low for the Asian Clade and most of NA Clade II, limiting our ability to assess variation in these species; however, members of NA Clade II do not have suites of shared inflorescence traits (*V. aestivalis, V. cinerea, V. labrusca, V. vulpina*; Klein *et al.*, 2018). Rather, *V. labrusca* has very small values for size-associated traits and larger values for local features compared to the other clade members, whereas *V. cinerea* has larger values for size-associated features and smaller values for local features (Fig. 7). This is consistent with the observation that aside from core phenotypic synapomorphies in the genus (tendril, bark, lenticel, and nodal diaphragm characters), members of NA Clade IIb (*V. aestivalis, V. cinerea, V. labrusca,* and *V. vulpina*) do not share morphological traits unique to the clade (Klein *et al.*, 2018). These species mostly co-occur across their distributions (Callen *et al.*, 2016) and additional sampling of *Vitis* taxa is necessary to further explore these complex evolutionary patterns. We observe *V. amurensis* grouping with *V. labrusca* and *V. coignetiae* grouping with North American species in hierarchical cluster analysis (Fig. 5A). The former two species have relatively smaller inflorescence architectures with thicker branches compared to the other species sampled here. Taxonomic relationships among North American and Asian *Vitis* species have been historically challenging, with clades comprised of species with disjunct distributions (Mullins *et al.*, 1992). Since current taxonomy resolves separate Asian and North American clades (Klein *et al.*, 2018), morphological similarity between these species likely reflects convergent evolution.

### Future Directions

Three-dimensional imaging through XRT and advanced mathematical approaches like persistent homology provide new ways to visualize and interpret complex biological structures including inflorescences, and to understand the genetic and environmental factors underlying variation in their architecture. In grapevines, cluster density is an important trait that is used to assess grapevine crop quality and to forecast yield, in part because of the association between bunch density and fungal infestations such as *Botrytis* (Hed *et al.*, 2009; Iland *et al.*, 2011; Molitor & Beyer, 2014; Molitor *et al.*, 2018). This study expands on previous work identifying variation in inflorescence architecture among cultivars (Shavrukov *et al.*, 2004), finding notable differences in cluster architecture among species. A logical next step may be to use 3D images and PH with topological modeling to trace the development of inflorescences across multiple growing seasons in a mapping population. Methods presented here are also amenable to scanning with berries, provided some noteworthy technical challenges are first addressed (e.g. minimizing berry damage and rotting during transportation, cluster stabilization during scanning, and segmentation of 3D volumes with features that vary widely in their X-ray absorbance). This work would provide a more complete representation of cluster structure, as well as inform our berry potential simulation with genotype-specific empirical data. We plan to develop predictive structural models of grapevine cluster development using these techniques.

Imaging and shape analysis approaches presented here can also be used to tease apart subtle environmental influences on inflorescence architecture, and the major agronomic trait of bunch density. Identifying environmental effects on phenotypic variation has important implications both for vineyard management and the assessment of intra-clone variation across geographic space. Cluster compactness can be manipulated through a variety of agronomic practices (Molitor et al. 2012; Gil et al. 2013; Frioni et al. 2017; Gourieroux et al. 2017; Poni et al. 2018; Reeve et al. 2018). Techniques described here can be used to quantify influences of specific treatments on cluster architecture. In addition, because grapevines are clonally propagated, clusters from the same widespread clones can be collected from different geographic locations, scanned and analyzed for variation. High resolution assessment of inflorescence architecture offers important insights into natural variation in bunch density and the genetic and environmental factors that influence it. The capacity to capture 3D variation in this complex trait over space and time represents a promising advance for a valuable potential target of selection in one of the most economically important berry crops in the world.

## Supporting information

SVideo1

SVideo2

## Supplementary data

**Fig. S1** A maximum likelihood phylogenetic tree for ten *Vitis* species.

**Fig. S2** Summary of inflorescence geometric and topological traits and the distribution for ten *Vitis* species.

**Fig. S3** Morphological traits mapped on the phylogenetic tree.

**Fig. S4.** Variation for each clade.

**Fig. S5** Pairwise correlations of morphological traits (allometric relationships) showing linear regression lines for each species.

**Fig. S6** Pairwise species classification.

**Table S1.** Trait description and calculation.

**Table S2.** Trait variance for each species.

**Table S3.** Trait loadings for two species classification.

**Table S4.** Trait Pagel’s lambda for phylogenetic analysis.

**Table S5.** Trait variation for each clade.

**Video S1** Illustration of quantifying branching topology using persistent homology.

**Video S2** Berry potential simulation

## Acknowledgements

The authors would like to acknowledge Elizabeth A. Kellogg (DDPSC) for valuable comments, particularly on phylogenetic analysis and inflorescence anatomy. We thank Noah Fahlgren (DDPSC) for computational assistance and Kari Miller (Washington University) for scanning assistance. We thank Zoë Migicovsky (Dalhousie University) for valuable comments.

This work was supported by funding from the United States National Science Foundation projects IIA-1355406, IOS-1638507, and DBI-1759796.

## Author contributions

CNT, DHC and JL designed the research; JL collected the samples and consulted on the biology; KD generated the X-ray data; LLK and AJM provided phylogenetic data and consulted for the biology; NJ and ML extracted pedicel diameter and angle; ML developed and extracted all the traits and conducted all the analysis and figures; ML, LLK, KD, JL, AJM, and CNT wrote the manuscript.

## Notes

#### Summary of Updates

Daniel H Chitwood was added as an author

