## SupplementaryData for "Characterizing grapevine 3D inflorescence architecture using X-ray imaging and advanced morphometrics: implications for understanding cluster density"

**Supplementary data**

Article title: Characterizing grapevine (*Vitis* spp.) inflorescence architecture using X-ray imaging: implications for understanding cluster density

Authors: Mao Li, Laura L. Klein, Keith E. Duncan, Ni Jiang, Daniel H. Chitwood, Jason Londo, Allison J. Miller, Christopher N. Topp

The following supplementary data is available for this article:

**Fig. S1** A maximum likelihood phylogenetic tree for ten *Vitis* species. An example for each species is shown with the same color highlighting the species name.

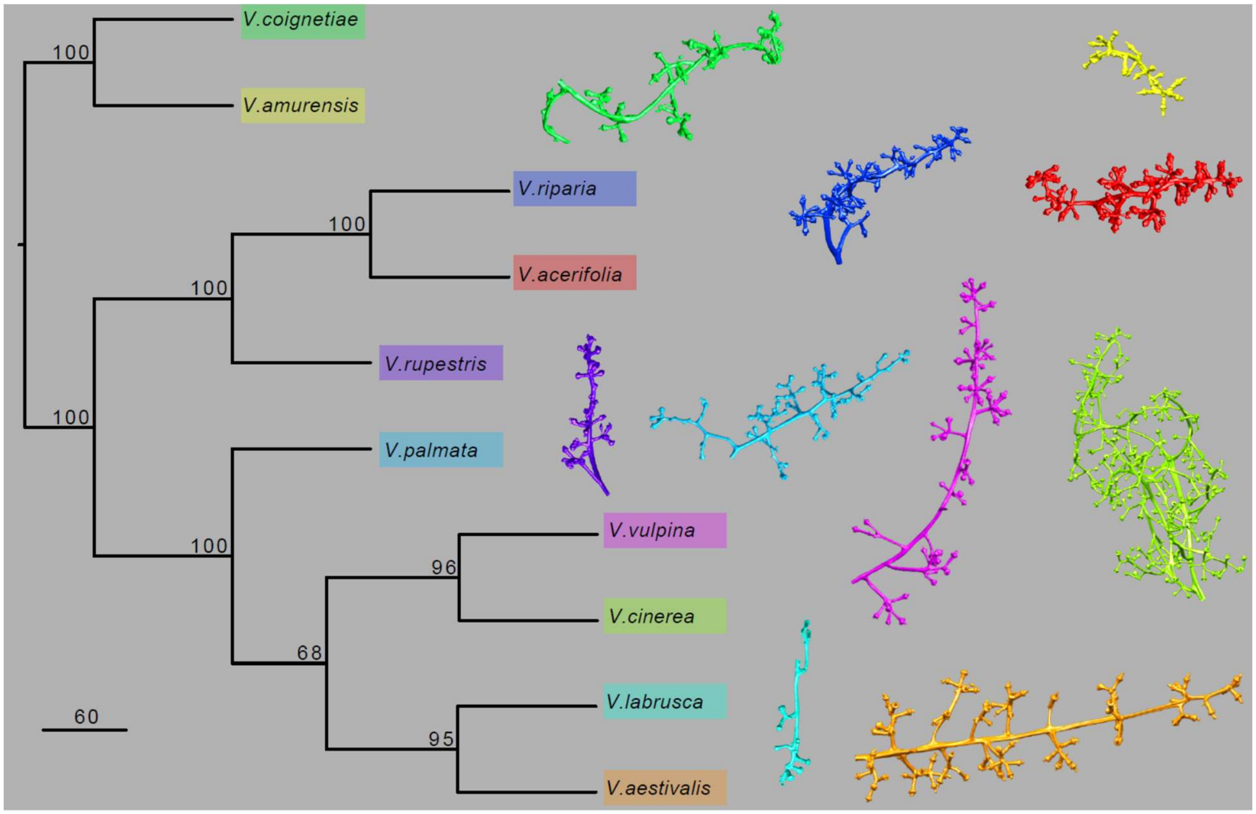


**Fig. S2** Summary of inflorescence geometric and topological traits and the distribution for ten *Vitis* species. Each panel shows seven traits, with each trait listed at the top of the column and two inflorescence examples demonstrating low and high trait values listed to the left. At the bottom of each column is a boxplot indicating the distribution and variance within 10 *Vitis* species, represented in different colors. On each box, each dot indicates an outlier if it is more than 1.5 interquartile ranges; the central vertical line indicates the median; the left and right edges of the box represent the 25th and 75th percentiles; and the whiskers extend to the most extreme nonoutlier data. The label for each species is listed in the boxplot y axis of the leftmost plot, with the number of individuals sampled for each species shown in parentheses. For a more detailed description of each trait, see Table S1.

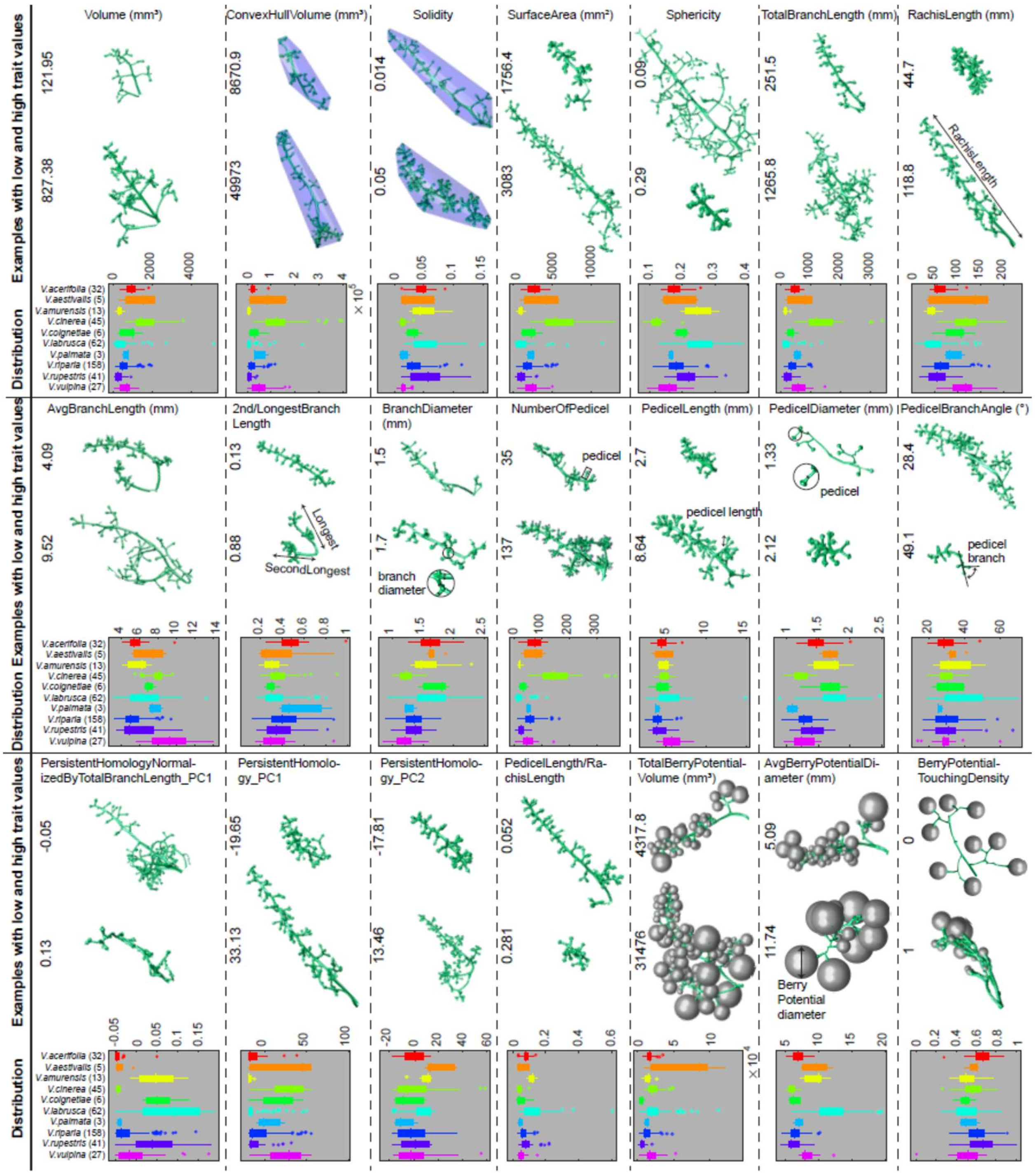

**Fig. S3** A Neighbor Joining phylogenetic tree for a subset of the *Vitis* data set (n=99). Node values denote bootstrap support for values. Ten *Vitis* species are highlighted in different colored backgrounds. Three clades (Asian Clade, NA Clade I, and NA Clade II) are labeled and marked by vertical bars. (a-x) Each trait is projected onto the phylogenetic tree branches, and indicates trait variation (red, lower values; blue, higher values) within individuals and among clades. Pagel's lambda value, an estimate of phylogenetic signal, and the p value are shown on the left top.

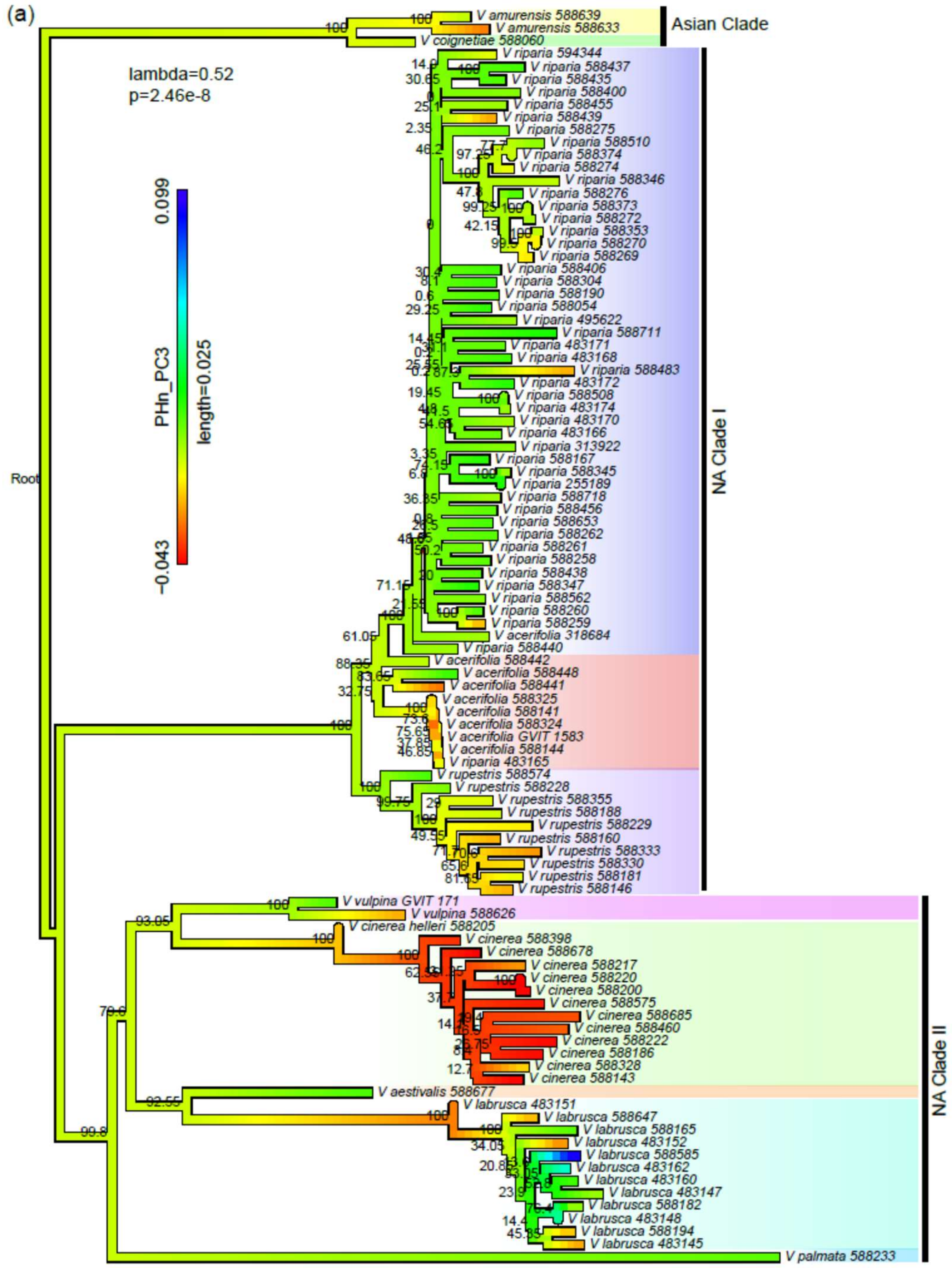

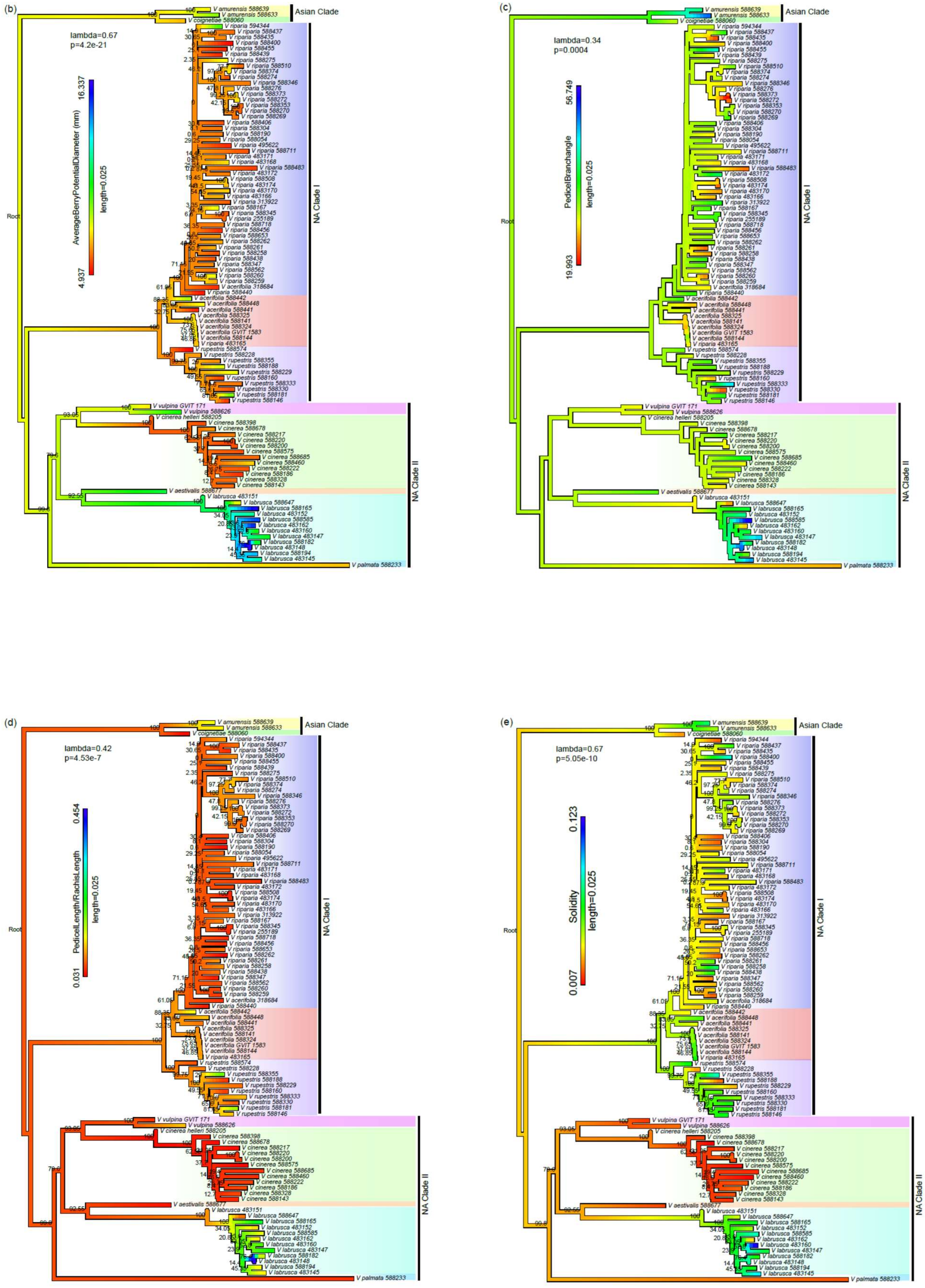
(b)
lambda=0.67
p=4.2e-21
AverageBerryPotentialDiameter (mm)
16.337
4.937
length=0.025
Root
Asian Clade
NA Clade I
NA Clade II
(c)
lambda=0.34
p=0.0004
PedicelBranchangle
56.749
19.993
length=0.025
Root
Asian Clade
NA Clade I
NA Clade II
(d)
lambda=0.42
p=4.53e-7
PedicelLength/RachisLength
0.454
0.031
length=0.025
Root
Asian Clade
NA Clade I
NA Clade II
(e)
lambda=0.67
p=5.05e-10
Solidity
0.123
0.007
length=0.025
Root
Asian Clade
NA Clade I
NA Clade II

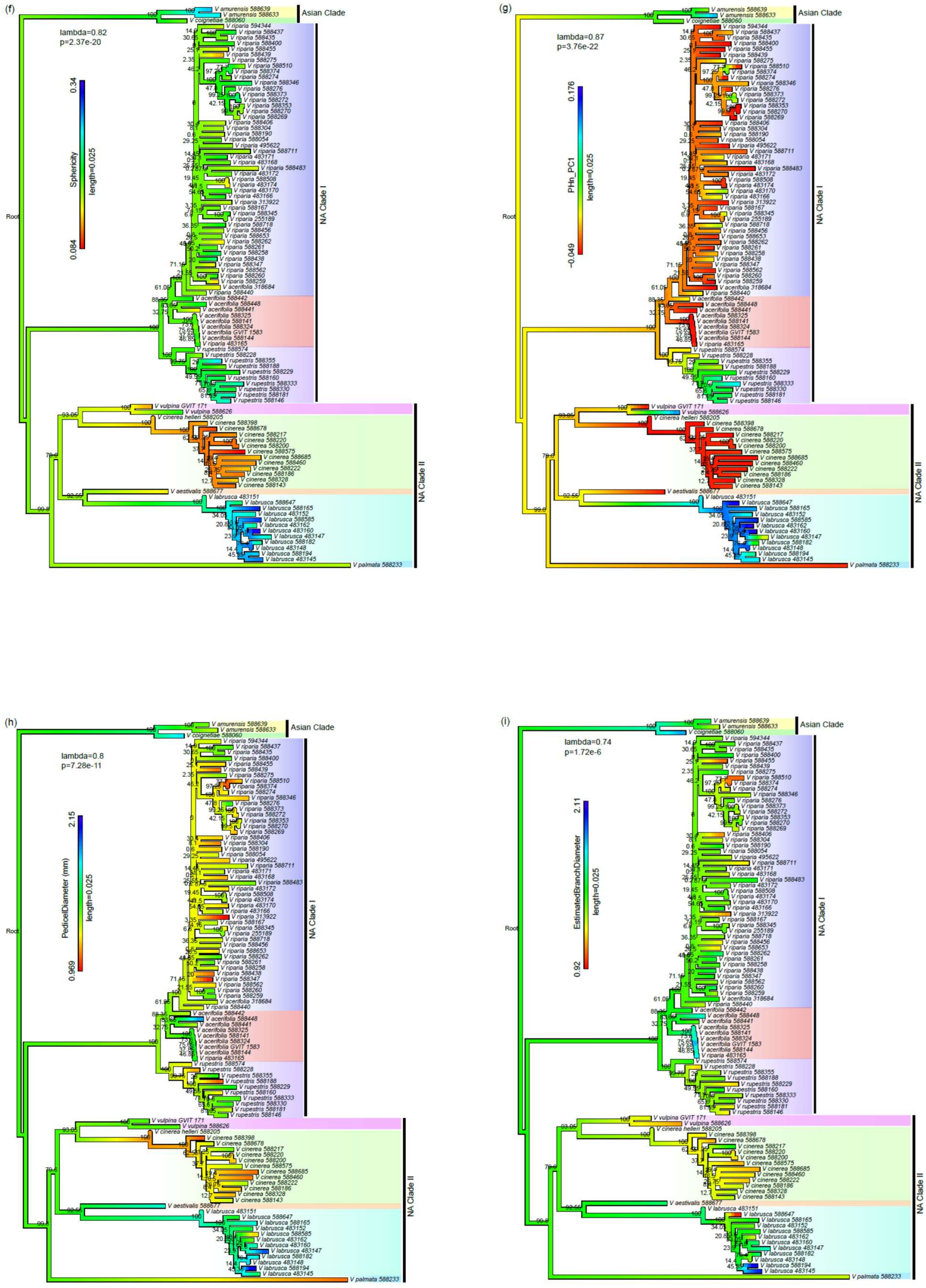

(f)
lambda=0.82
p=2.37e-20
0.34
Sphericity
length=0.025
0.084
Asian Clade
NA Clade I
NA Clade II
Root
(g)
lambda=0.87
p=3.76e-22
0.176
PHn_PC1
length=0.025
-0.049
Asian Clade
NA Clade I
NA Clade II
Root
(h)
lambda=0.8
p=7.28e-11
2.15
PedicelDiameter (mm)
length=0.025
0.969
Asian Clade
NA Clade I
NA Clade II
Root
(i)
lambda=0.74
p=1.72e-6
2.11
EstimatedBranchDiameter
length=0.025
0.92
Asian Clade
NA Clade I
NA Clade II
Root

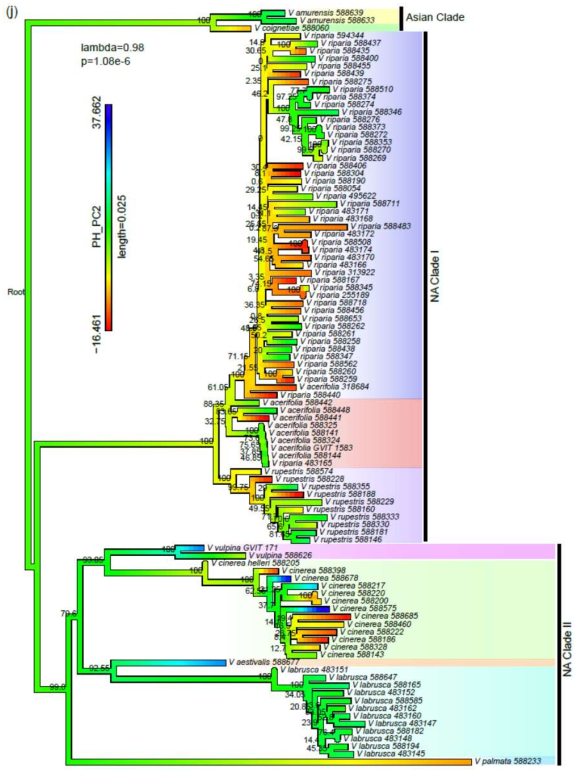
(j)
lambda=0.98
p=1.08e-6
37.662
PH_PC2
length=0.025
-16.461
Root
Asian Clade
NA Clade I
NA Clade II

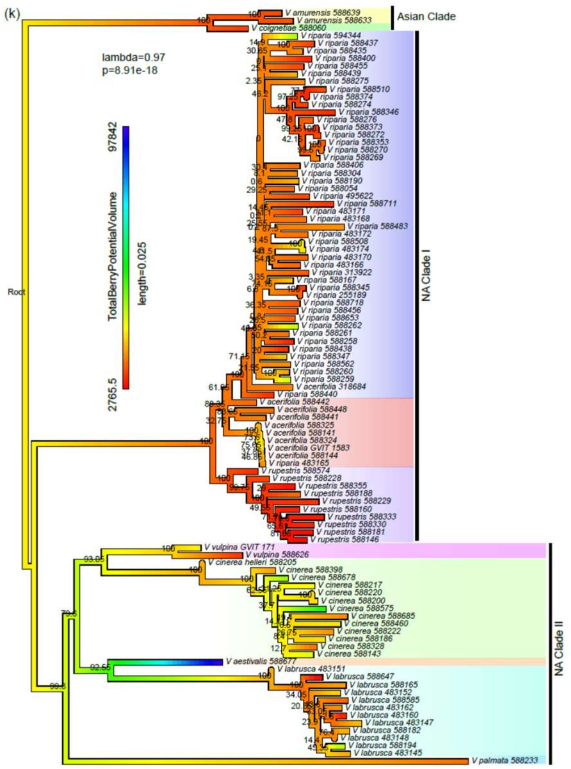
(k)
lambda=0.97
p=8.91e-18
97842
TotalBerryPotentialVolume
length=0.025
2765.5
Root
Asian Clade
NA Clade I
NA Clade II

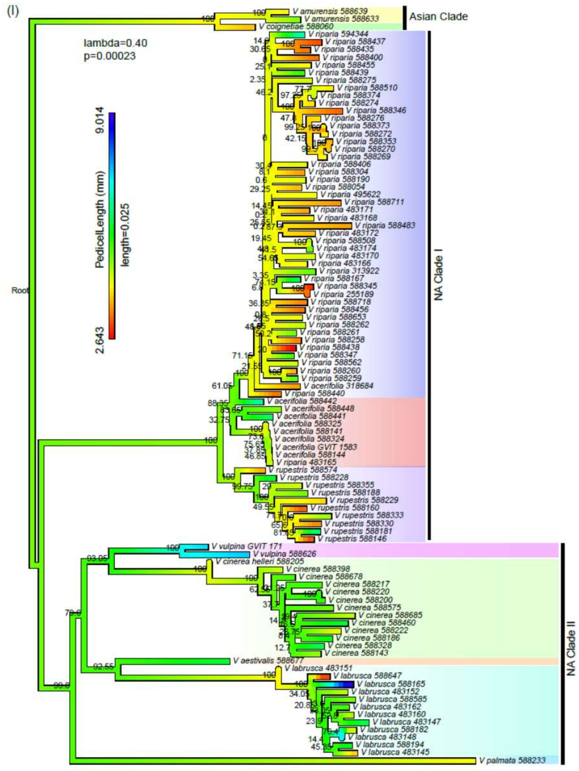
(l)
lambda=0.40
p=0.00023
9.014
PedicelLength (mm)
length=0.025
2.643
Root
Asian Clade
NA Clade I
NA Clade II

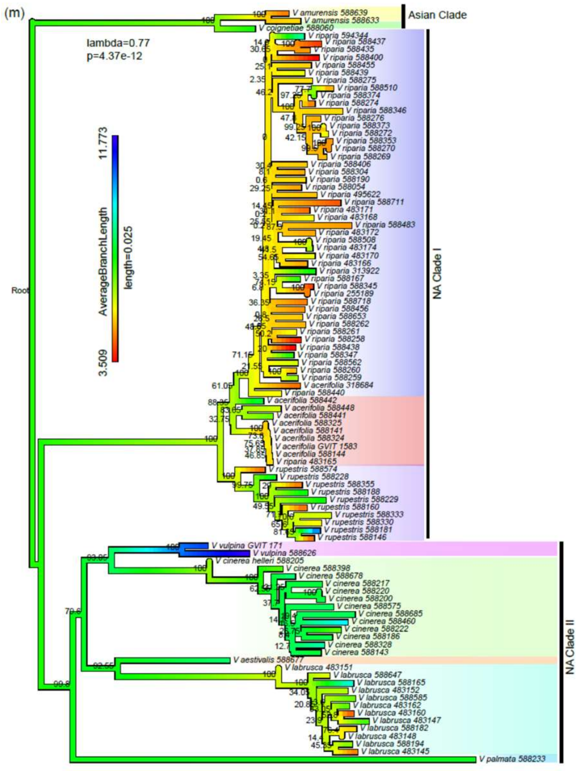
(m)
lambda=0.77
p=4.37e-12
11.773
AverageBranchLength
length=0.025
3.509
Root
Asian Clade
NA Clade I
NA Clade II

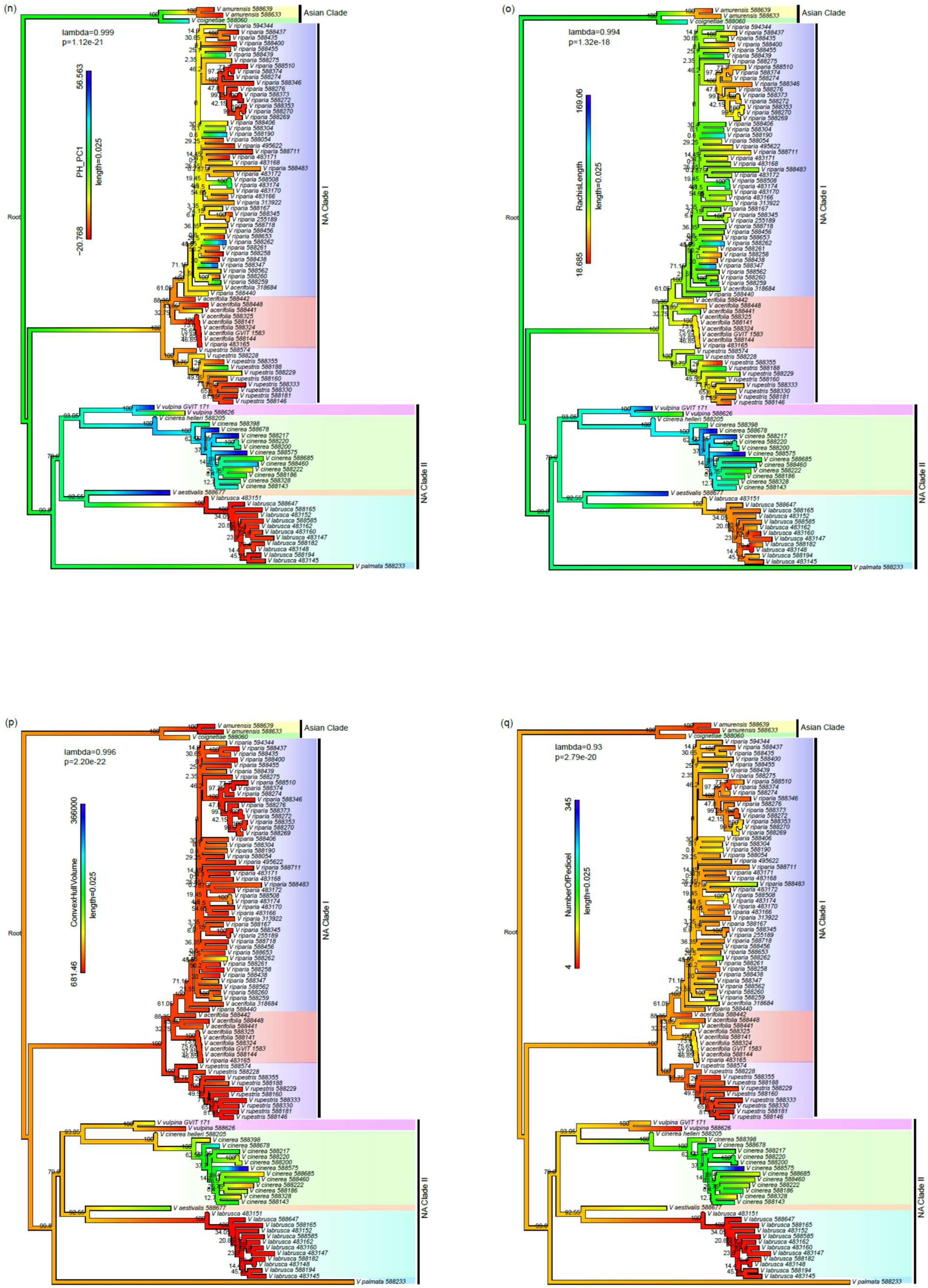
(n)
lambda=0.999
p=1.12e-21
PH_PC1
length=0.025
56.563
−20.768
Asian Clade
NA Clade I
NA Clade II
Root
(o)
lambda=0.994
p=1.32e-18
RachisLength
length=0.025
169.06
18.685
Asian Clade
NA Clade I
NA Clade II
Root
(p)
lambda=0.996
p=2.20e-22
ConvexHullVolume
length=0.025
366000
681.46
Asian Clade
NA Clade I
NA Clade II
Root
(q)
lambda=0.93
p=2.79e-20
NumberOfPedicel
length=0.025
345
4
Asian Clade
NA Clade I
NA Clade II
Root

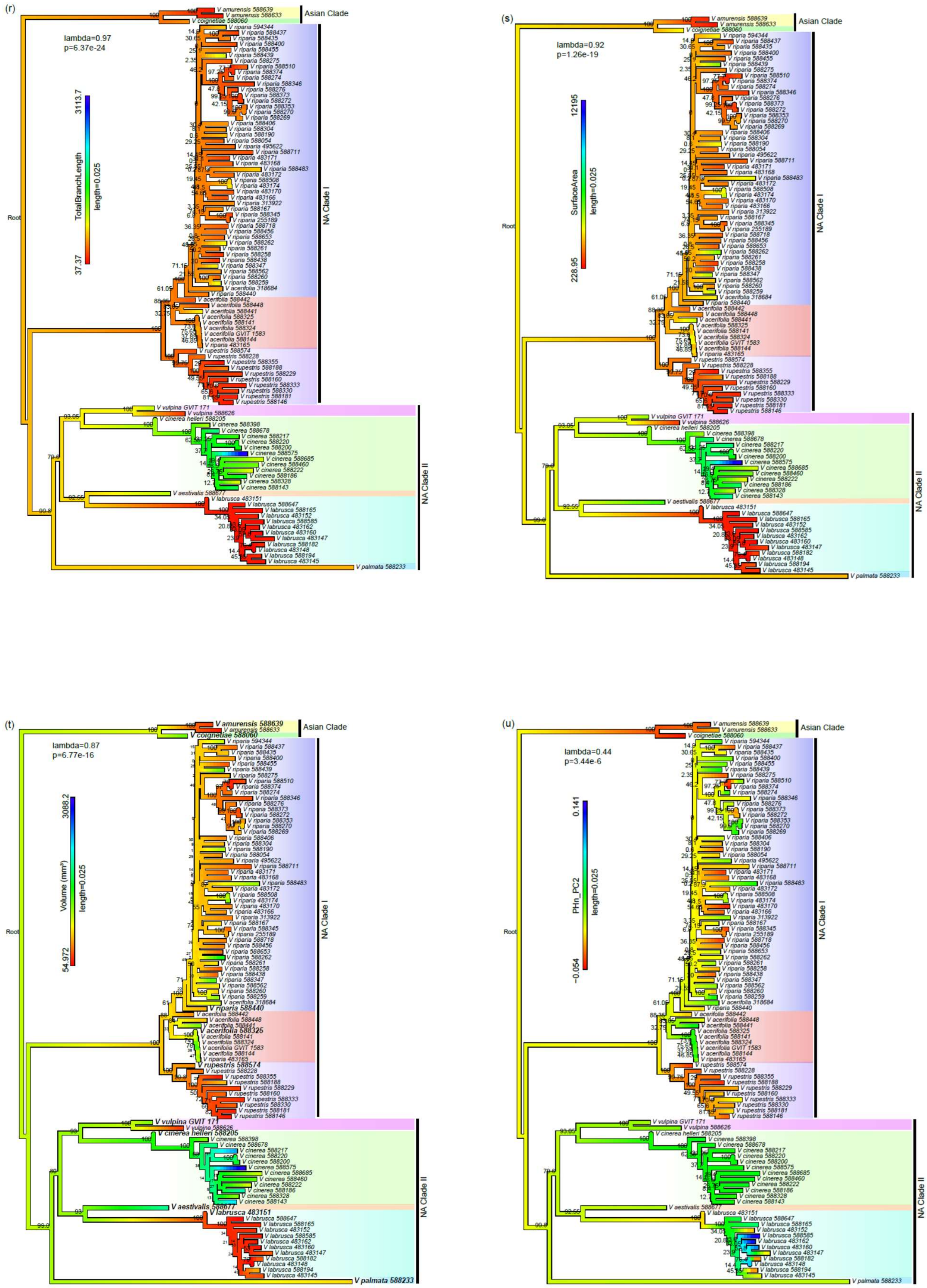
(r)
lambda=0.97
p=6.37e-24
TotalBranchLength
length=0.025
3113.7
37.37
Root
Asian Clade
NA Clade I
NA Clade II
(s)
lambda=0.92
p=1.26e-19
SurfaceArea
length=0.025
12195
228.95
Root
Asian Clade
NA Clade I
NA Clade II
(t)
lambda=0.87
p=6.77e-16
Volume (mm³)
length=0.025
3088.2
54.972
Root
Asian Clade
NA Clade I
NA Clade II
(u)
lambda=0.44
p=3.44e-6
PHn_PC2
length=0.025
0.141
−0.054
Root
Asian Clade
NA Clade I
NA Clade II

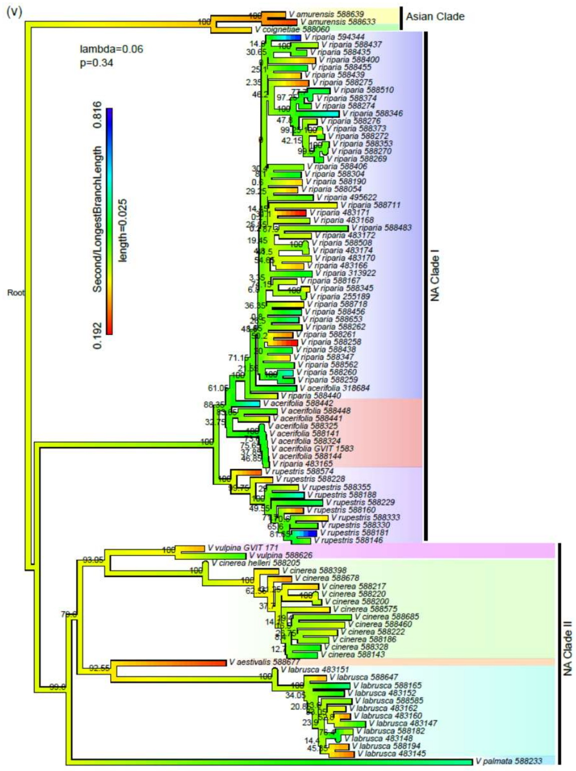
(v)
lambda=0.06
p=0.34
Second/LongestBranchLength
0.816
0.192
length=0.025
Root
Asian Clade
NA Clade I
NA Clade II

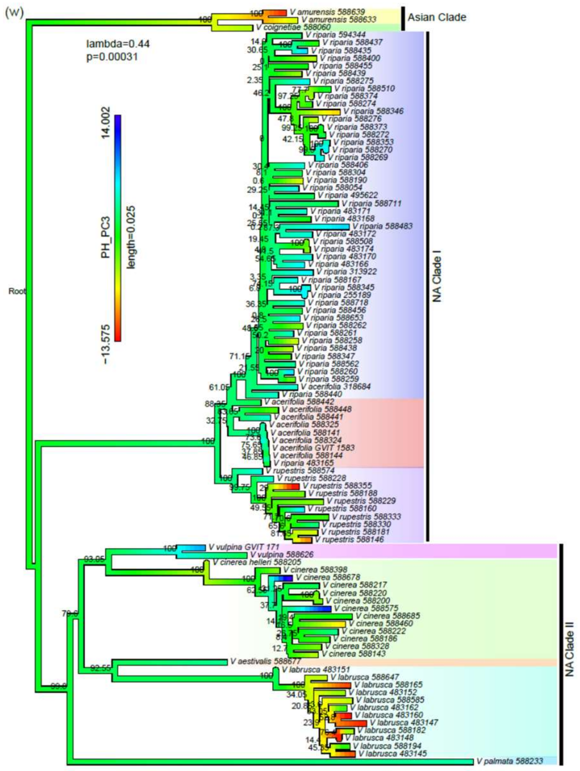
(w)
lambda=0.44
p=0.00031
PH_PC3
14.002
−13.575
length=0.025
Root
Asian Clade
NA Clade I
NA Clade II

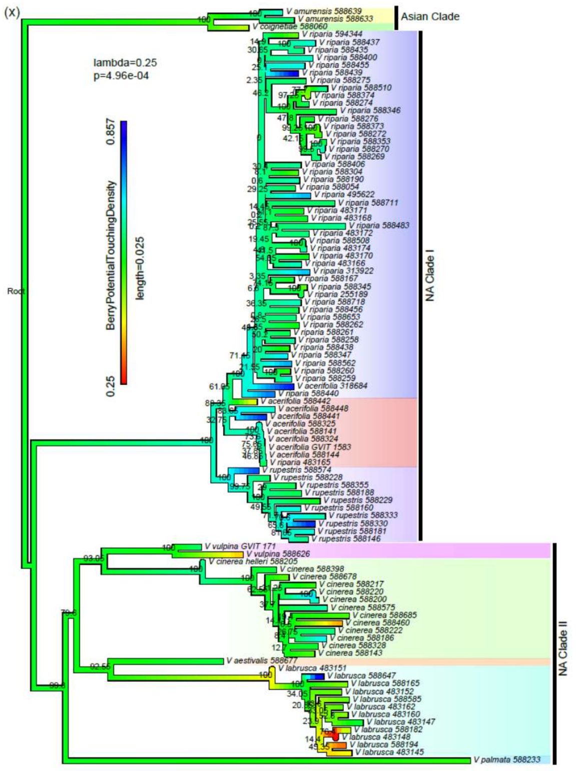
(x)
lambda=0.25
p=4.96e-04
BerryPotentialTouchingDensity
0.857
0.25
length=0.025
Root
Asian Clade
NA Clade I
NA Clade II

**Fig. S4** Variation for each clade. The variance of multivariate (top) and each rachis morphological trait for each clade based on the mean value for each species.

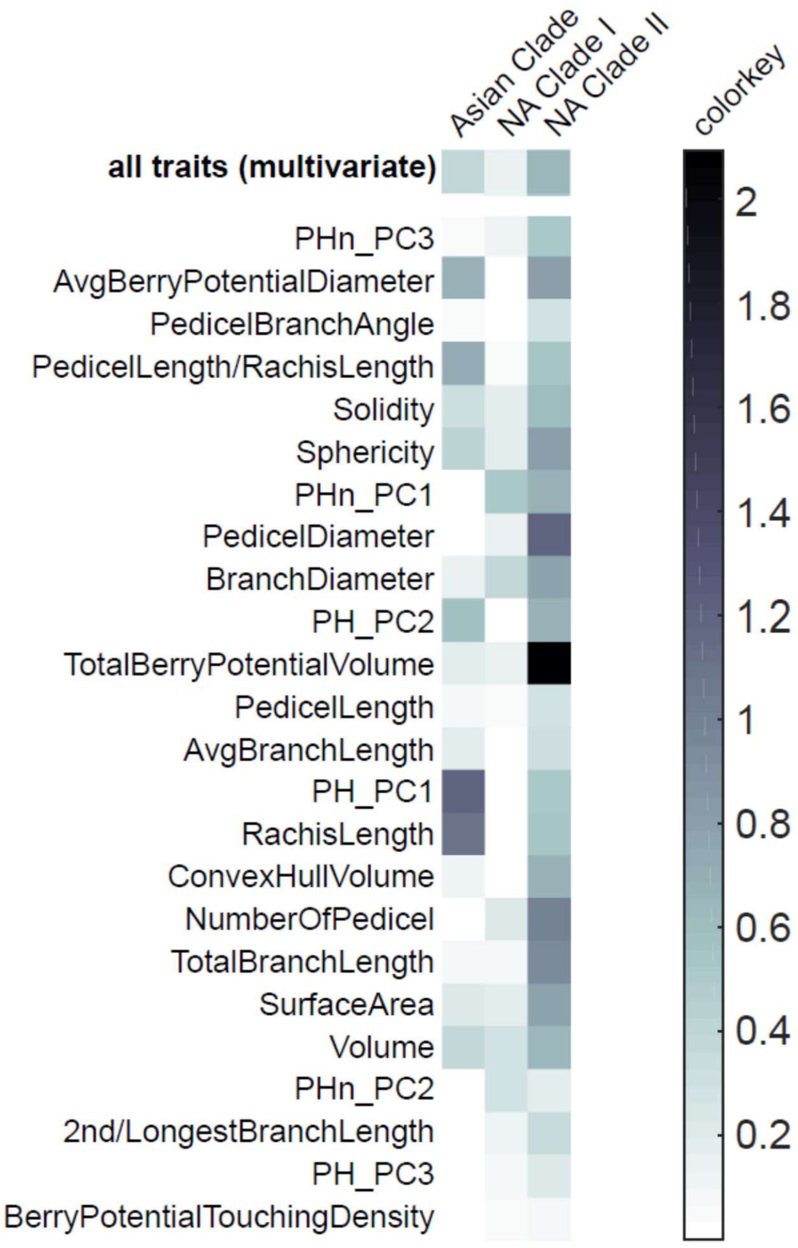


**Fig. S5** Pairwise correlations of morphological traits (allometric relationships) showing linear regression lines for each species.

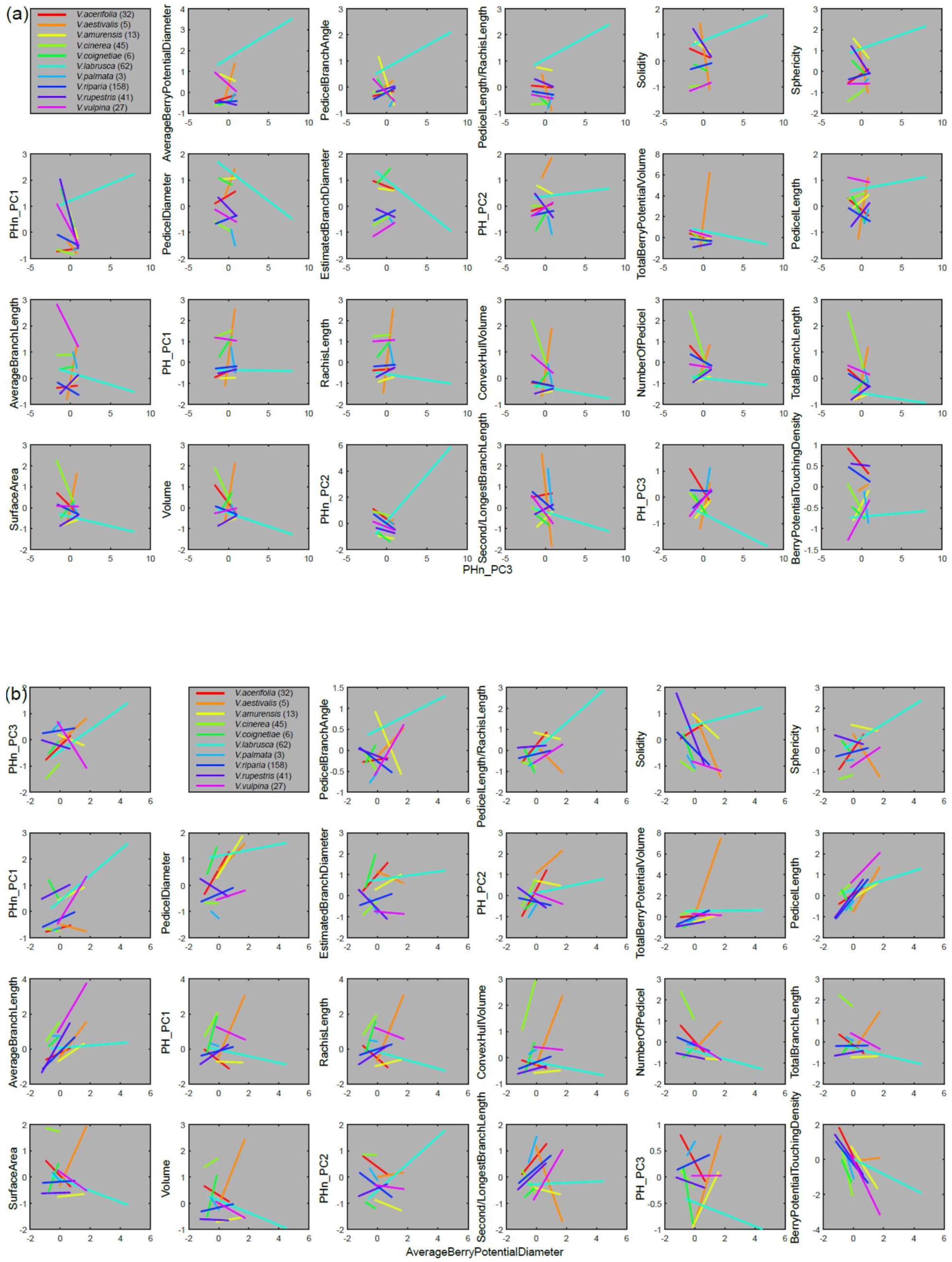
(a)
V.acerifolia (32)
V.aestivalis (5)
V.amurensis (13)
V.cinerea (45)
V.coignetiae (6)
V.labrusca (62)
V.palmata (3)
V.riparia (158)
V.rupestris (41)
V.vulpina (27)
AverageBerryPotentialDiameter
PedicelBranchAngle
PedicelLength/RachisLength
Solidity
Sphericity
PHn_PC1
PedicelDiameter
EstimatedBranchDiameter
PH_PC2
TotalBerryPotentialVolume
PedicelLength
AverageBranchLength
PH_PC1
RachisLength
ConvexHullVolume
NumberOfPedicel
TotalBranchLength
SurfaceArea
Volume
PHn_PC2
Second/LongestBranchLength
PH_PC3
BerryPotentialTouchingDensity
PHn_PC3
(b)
PHn_PC3
V.acerifolia (32)
V.aestivalis (5)
V.amurensis (13)
V.cinerea (45)
V.coignetiae (6)
V.labrusca (62)
V.palmata (3)
V.riparia (158)
V.rupestris (41)
V.vulpina (27)
PedicelBranchAngle
PedicelLength/RachisLength
Solidity
Sphericity
PHn_PC1
PedicelDiameter
EstimatedBranchDiameter
PH_PC2
TotalBerryPotentialVolume
PedicelLength
AverageBranchLength
PH_PC1
RachisLength
ConvexHullVolume
NumberOfPedicel
TotalBranchLength
SurfaceArea
Volume
PHn_PC2
Second/LongestBranchLength
PH_PC3
BerryPotentialTouchingDensity
AverageBerryPotentialDiameter

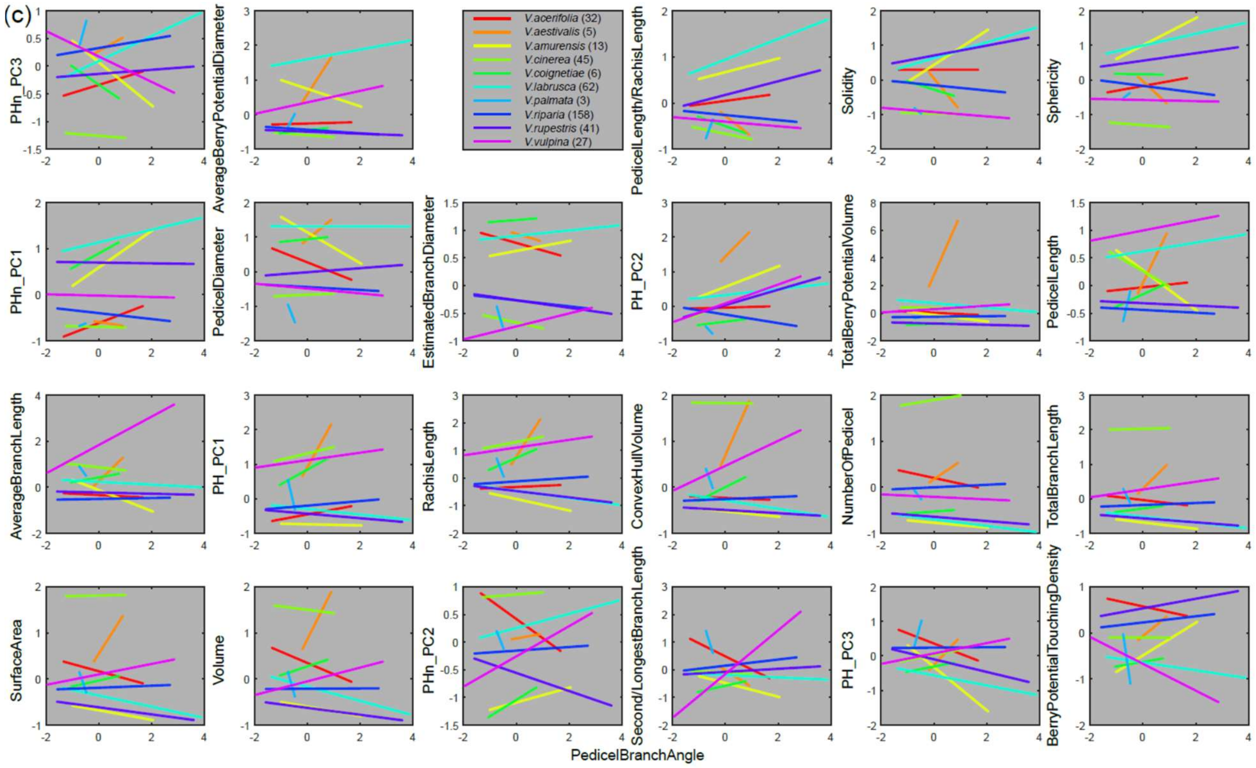
(c)
V.acerifolia (32)
V.aestivalis (5)
V.amurensis (13)
V.cinerea (45)
V.coignetiae (6)
V.labrusca (62)
V.palmata (3)
V.riparia (158)
V.rupestris (41)
V.vulpina (27)
PHn_PC3
AverageBerryPotentialDiameter
PedicelLength/RachisLength
Solidity
Sphericity
PHn_PC1
PedicelDiameter
EstimatedBranchDiameter
PH_PC2
TotalBerryPotentialVolume
PedicelLength
AverageBranchLength
PH_PC1
RachisLength
ConvexHullVolume
NumberOfPedicel
TotalBranchLength
SurfaceArea
Volume
PHn_PC2
Second/LongestBranchLength
PH_PC3
BerryPotentialTouchingDensity
PedicelBranchAngle

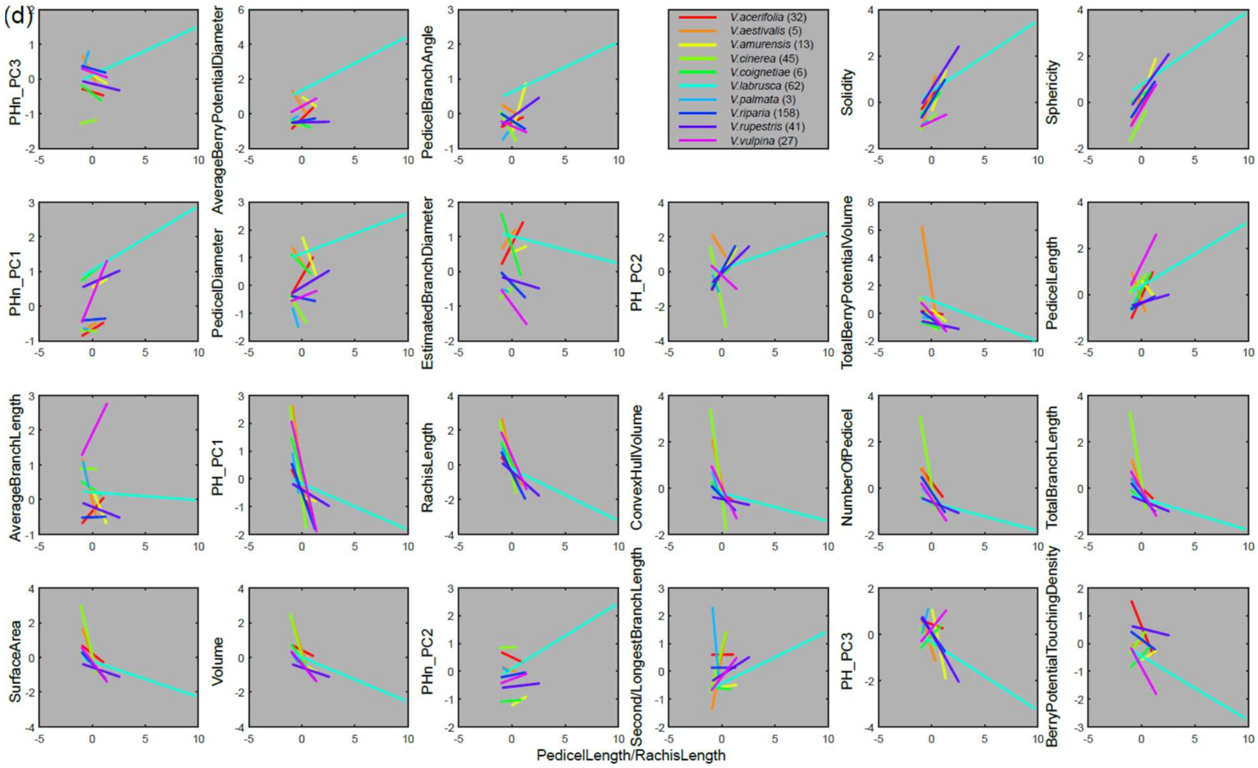
(d)
V.acerifolia (32)
V.aestivalis (5)
V.amurensis (13)
V.cinerea (45)
V.coignetiae (6)
V.labrusca (62)
V.palmata (3)
V.riparia (158)
V.rupestris (41)
V.vulpina (27)
PHn_PC3
AverageBerryPotentialDiameter
PedicelBranchAngle
Solidity
Sphericity
PHn_PC1
PedicelDiameter
EstimatedBranchDiameter
PH_PC2
TotalBerryPotentialVolume
PedicelLength
AverageBranchLength
PH_PC1
RachisLength
ConvexHullVolume
NumberOfPedicel
TotalBranchLength
SurfaceArea
Volume
PHn_PC2
Second/LongestBranchLength
PH_PC3
BerryPotentialTouchingDensity
PedicelLength/RachisLength

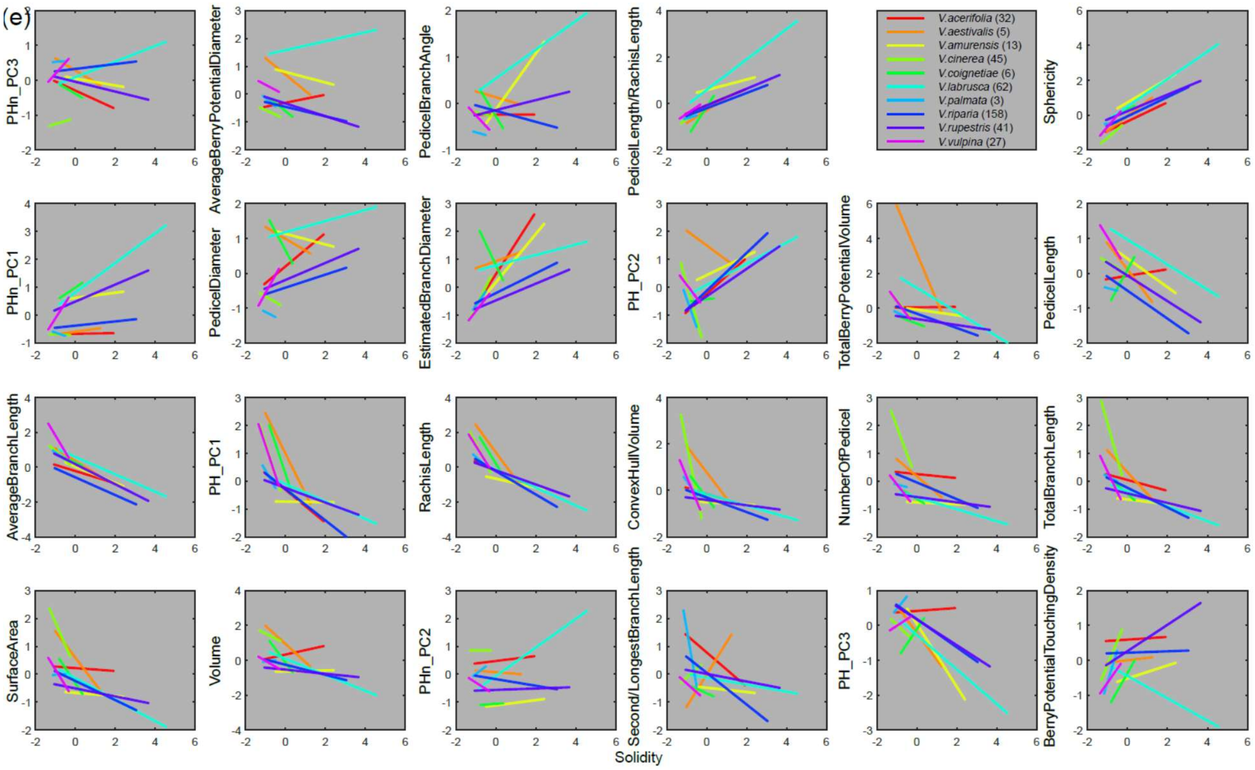
(e)
PHn_PC3
AverageBerryPotentialDiameter
PedicelBranchAngle
PedicelLength/RachisLength
V.acerifolia (32)
V.aestivalis (5)
V.amurensis (13)
V.cinerea (45)
V.coignetiae (6)
V.labrusca (62)
V.palmata (3)
V.riparia (158)
V.rupestris (41)
V.vulpina (27)
Sphericity
PHn_PC1
PedicelDiameter
EstimatedBranchDiameter
PH_PC2
TotalBerryPotentialVolume
PedicelLength
AverageBranchLength
PH_PC1
RachisLength
ConvexHullVolume
NumberOfPedicel
TotalBranchLength
SurfaceArea
Volume
PHn_PC2
Second/LongestBranchLength
PH_PC3
BerryPotentialTouchingDensity
Solidity

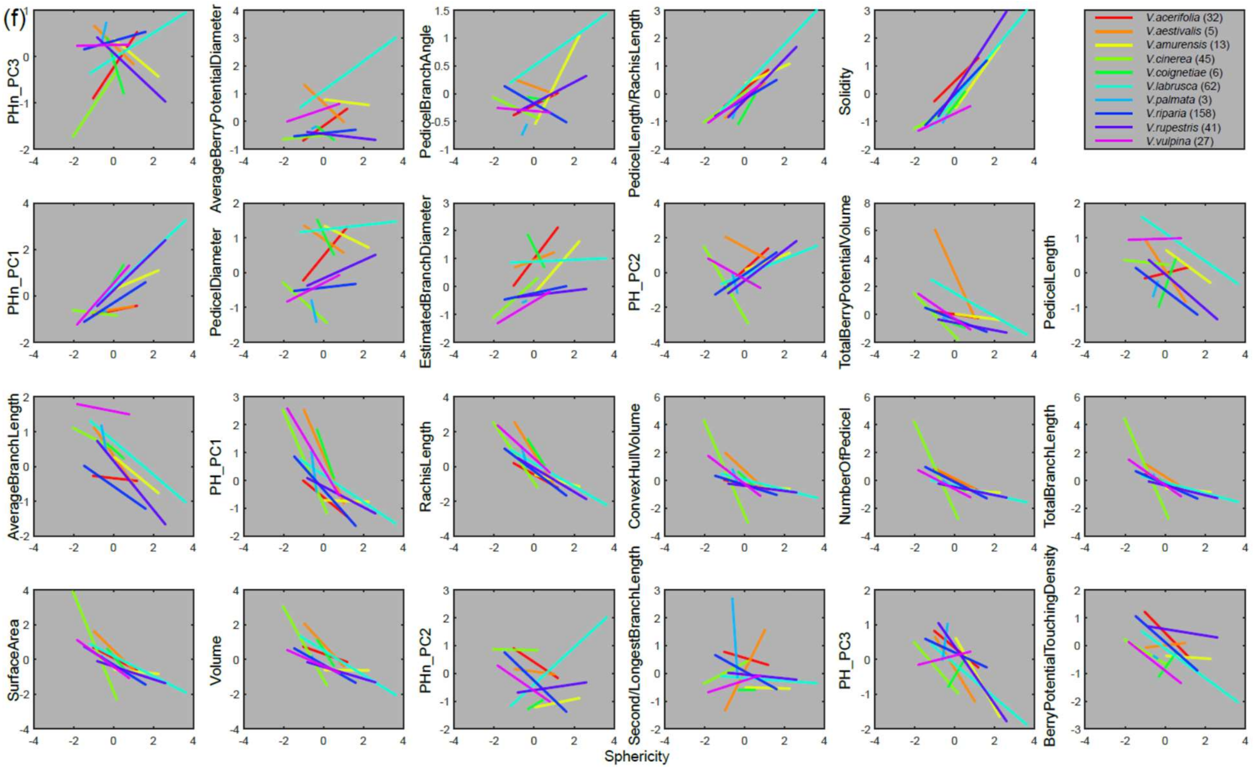
(f)
PHn_PC3
AverageBerryPotentialDiameter
PedicelBranchAngle
PedicelLength/RachisLength
Solidity
V.acerifolia (32)
V.aestivalis (5)
V.amurensis (13)
V.cinerea (45)
V.coignetiae (6)
V.labrusca (62)
V.palmata (3)
V.riparia (158)
V.rupestris (41)
V.vulpina (27)
PHn_PC1
PedicelDiameter
EstimatedBranchDiameter
PH_PC2
TotalBerryPotentialVolume
PedicelLength
AverageBranchLength
PH_PC1
RachisLength
ConvexHullVolume
NumberOfPedicel
TotalBranchLength
SurfaceArea
Volume
PHn_PC2
Second/LongestBranchLength
PH_PC3
BerryPotentialTouchingDensity
Sphericity

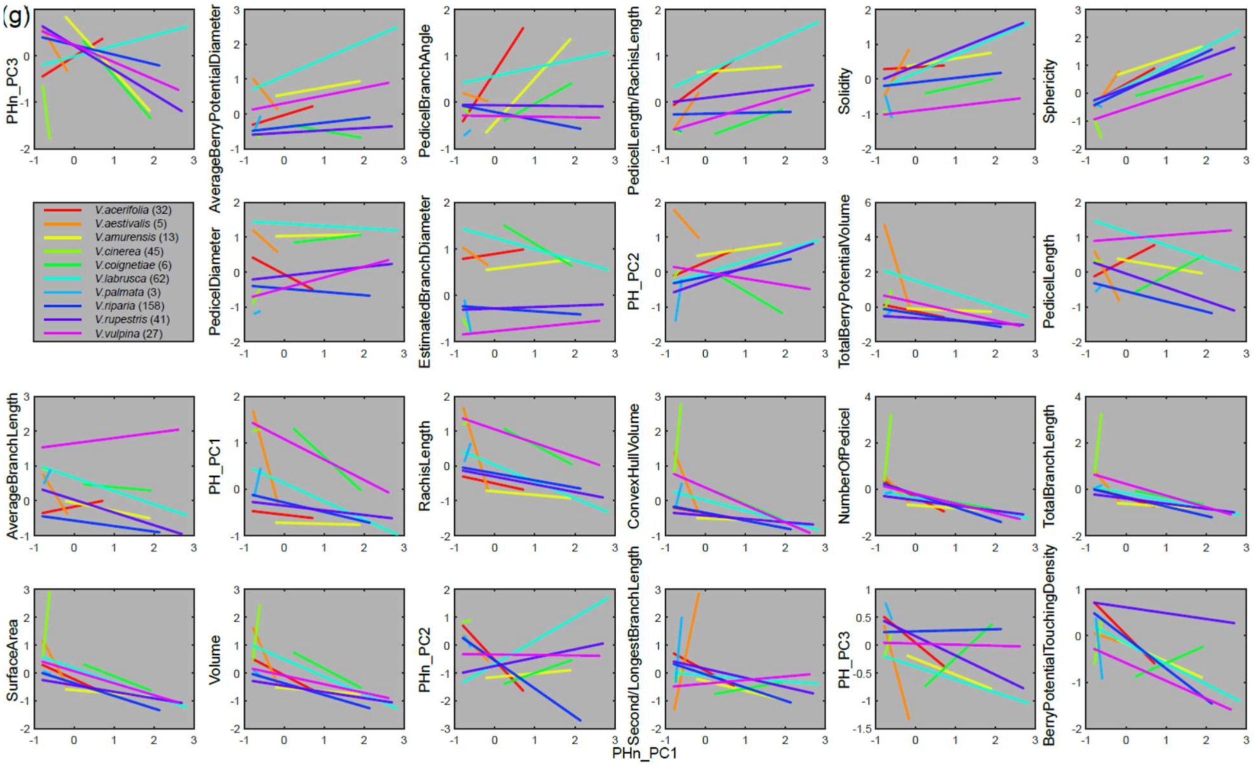
(g)
PHn_PC3
AveragePotentialBerryDiameter
PedicelBranchAngle
PedicelLength/RachisLength
Solidity
Sphericity
V.acerifolia (32)
V.aestivalis (5)
V.amurensis (13)
V.cinerea (45)
V.coignetiae (6)
V.labrusca (62)
V.palmata (3)
V.riparia (158)
V.rupestris (41)
V.vulpina (27)
PedicelDiameter
EstimatedBranchDiameter
PH_PC2
TotalBerryPotentialVolume
PedicelLength
AverageBranchLength
PH_PC1
RachisLength
ConvexHullVolume
NumberOfPedicel
TotalBranchLength
SurfaceArea
Volume
PHn_PC2
Second/LongestBranchLength
PH_PC3
BerryPotentialTouchingDensity
PHn_PC1

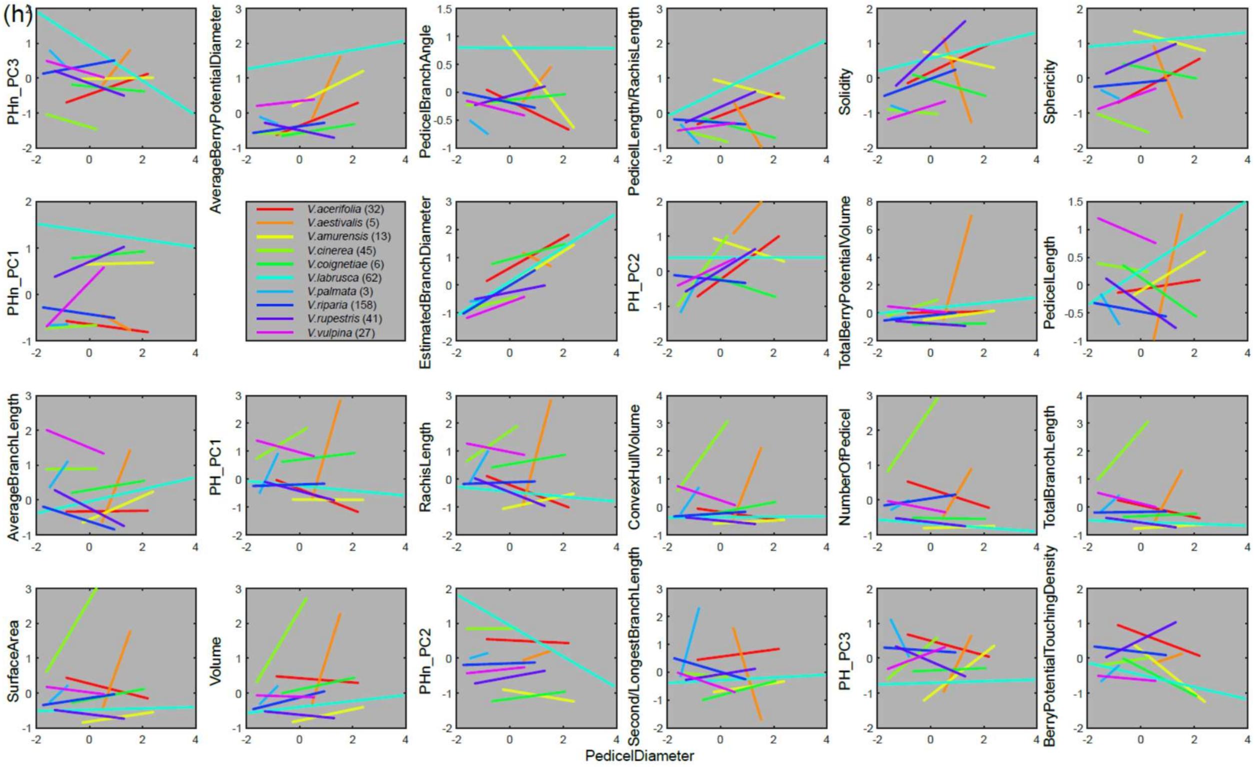
(h)
PHn_PC3
AveragePotentialBerryDiameter
PedicelBranchAngle
PedicelLength/RachisLength
Solidity
Sphericity
PHn_PC1
V.acerifolia (32)
V.aestivalis (5)
V.amurensis (13)
V.cinerea (45)
V.coignetiae (6)
V.labrusca (62)
V.palmata (3)
V.riparia (158)
V.rupestris (41)
V.vulpina (27)
EstimatedBranchDiameter
PH_PC2
TotalBerryPotentialVolume
PedicelLength
AverageBranchLength
PH_PC1
RachisLength
ConvexHullVolume
NumberOfPedicel
TotalBranchLength
SurfaceArea
Volume
PHn_PC2
Second/LongestBranchLength
PH_PC3
BerryPotentialTouchingDensity
PedicelDiameter

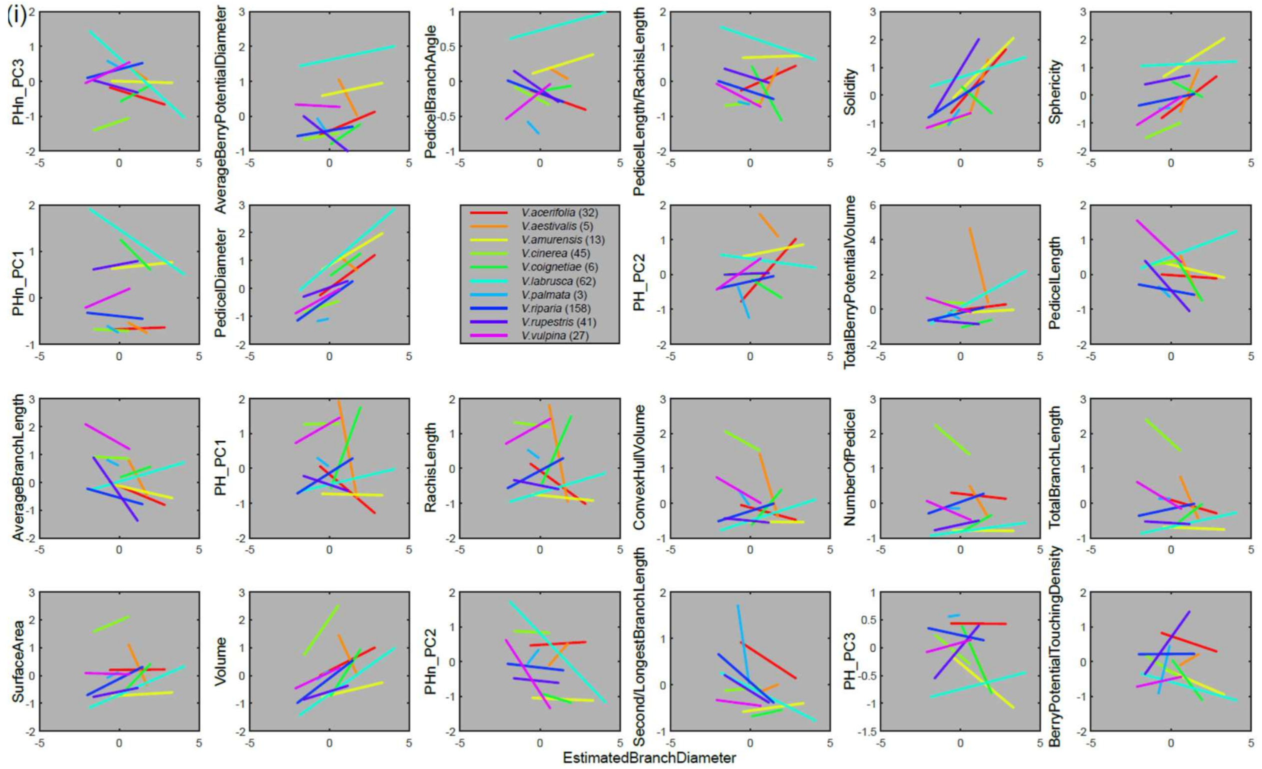
(i)
PHn_PC3
AverageBerryPotentialDiameter
PedicelBranchAngle
PedicelLength/RachisLength
Solidity
Sphericity
PHn_PC1
PedicelDiameter
V.acerifolia (32)
V.aestivalis (5)
V.amurensis (13)
V.cinerea (45)
V.coignetiae (6)
V.labrusca (62)
V.palmata (3)
V.riparia (158)
V.rupestris (41)
V.vulpina (27)
PH_PC2
TotalBerryPotentialVolume
PedicelLength
AverageBranchLength
PH_PC1
RachisLength
ConvexHullVolume
NumberOfPedicel
TotalBranchLength
SurfaceArea
Volume
PHn_PC2
Second/LongestBranchLength
PH_PC3
BerryPotentialTouchingDensity
EstimatedBranchDiameter

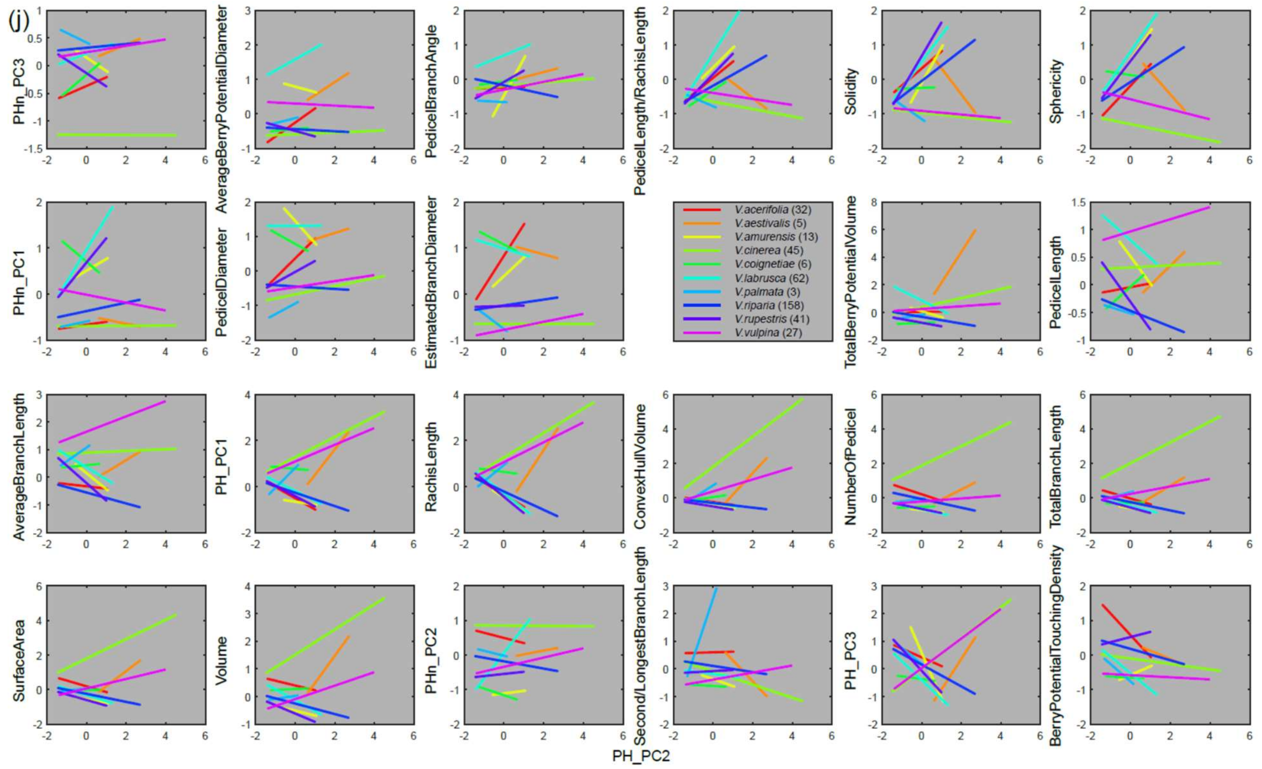
(j)
PHn_PC3
AverageBerryPotentialDiameter
PedicelBranchAngle
PedicelLength/RachisLength
Solidity
Sphericity
PHn_PC1
PedicelDiameter
EstimatedBranchDiameter
V.acerifolia (32)
V.aestivalis (5)
V.amurensis (13)
V.cinerea (45)
V.coignetiae (6)
V.labrusca (62)
V.palmata (3)
V.riparia (158)
V.rupestris (41)
V.vulpina (27)
TotalBerryPotentialVolume
PedicelLength
AverageBranchLength
PH_PC1
RachisLength
ConvexHullVolume
NumberOfPedicel
TotalBranchLength
SurfaceArea
Volume
PHn_PC2
Second/LongestBranchLength
PH_PC3
BerryPotentialTouchingDensity
PH_PC2

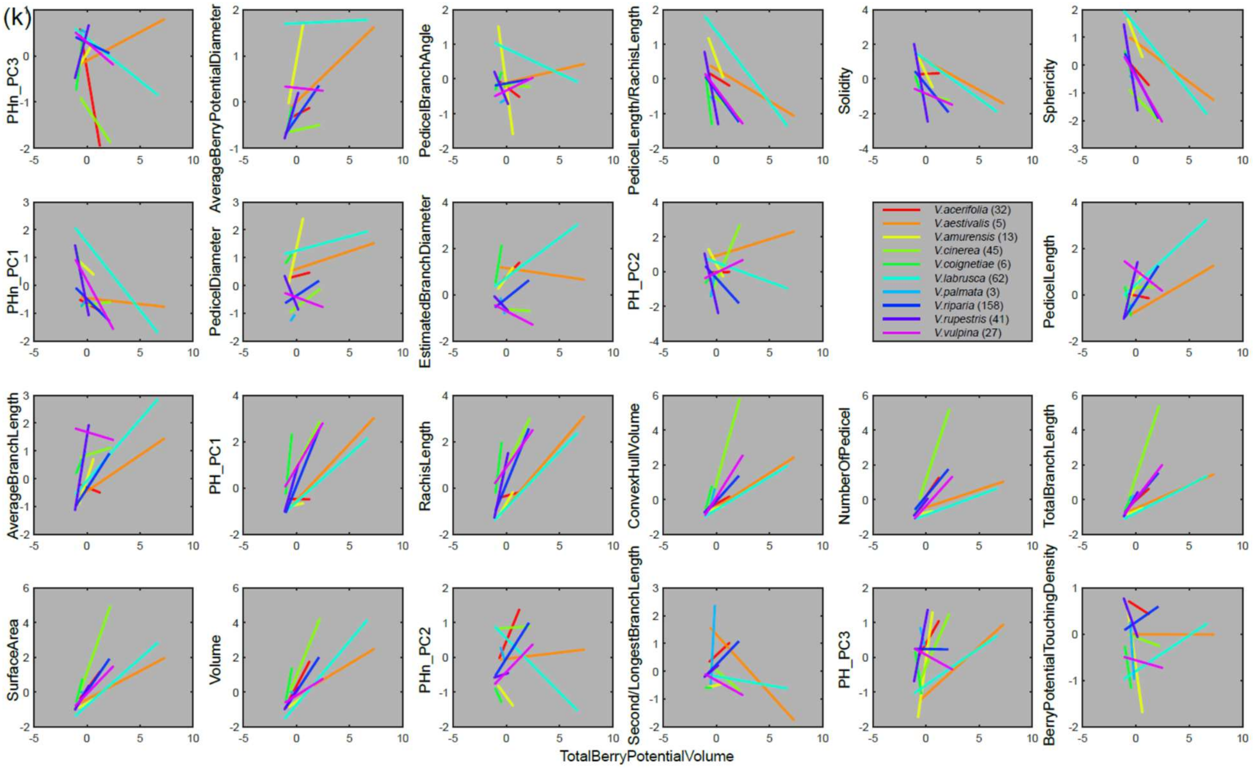
(k)
PHn_PC3
AverageBerryPotentialDiameter
PedicelBranchAngle
PedicelLength/RachisLength
Solidity
Sphericity
PHn_PC1
PedicelDiameter
EstimatedBranchDiameter
PH_PC2
PedicelLength
AverageBranchLength
PH_PC1
RachisLength
ConvexHullVolume
NumberOfPedicel
TotalBranchLength
SurfaceArea
Volume
PHn_PC2
Second/LongestBranchLength
PH_PC3
BerryPotentialTouchingDensity
TotalBerryPotentialVolume
V.acerifolia (32)
V.aestivalis (5)
V.amurensis (13)
V.cinerea (45)
V.coignetiae (6)
V.labrusca (62)
V.palmata (3)
V.riparia (158)
V.rupestris (41)
V.vulpina (27)

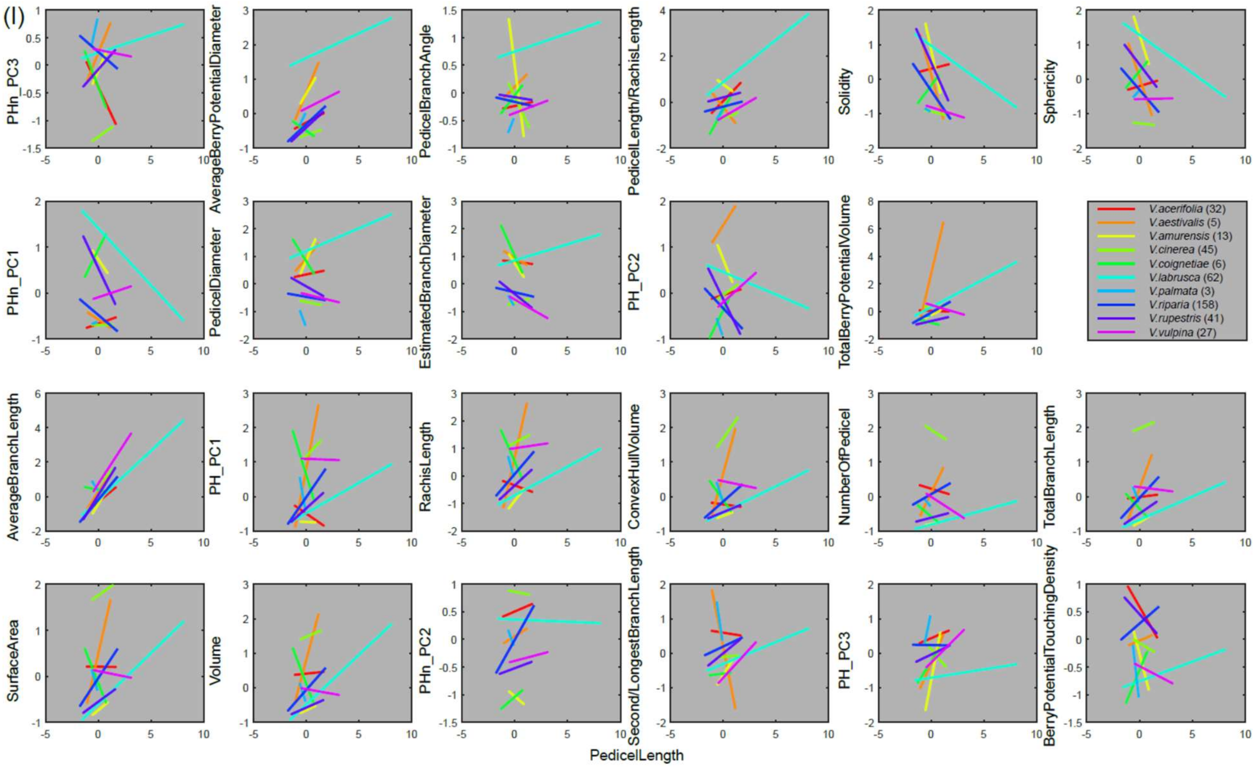
(l)
PHn_PC3
AverageBerryPotentialDiameter
PedicelBranchAngle
PedicelLength/RachisLength
Solidity
Sphericity
PHn_PC1
PedicelDiameter
EstimatedBranchDiameter
PH_PC2
TotalBerryPotentialVolume
AverageBranchLength
PH_PC1
RachisLength
ConvexHullVolume
NumberOfPedicel
TotalBranchLength
SurfaceArea
Volume
PHn_PC2
Second/LongestBranchLength
PH_PC3
BerryPotentialTouchingDensity
PedicelLength
V.acerifolia (32)
V.aestivalis (5)
V.amurensis (13)
V.cinerea (45)
V.coignetiae (6)
V.labrusca (62)
V.palmata (3)
V.riparia (158)
V.rupestris (41)
V.vulpina (27)

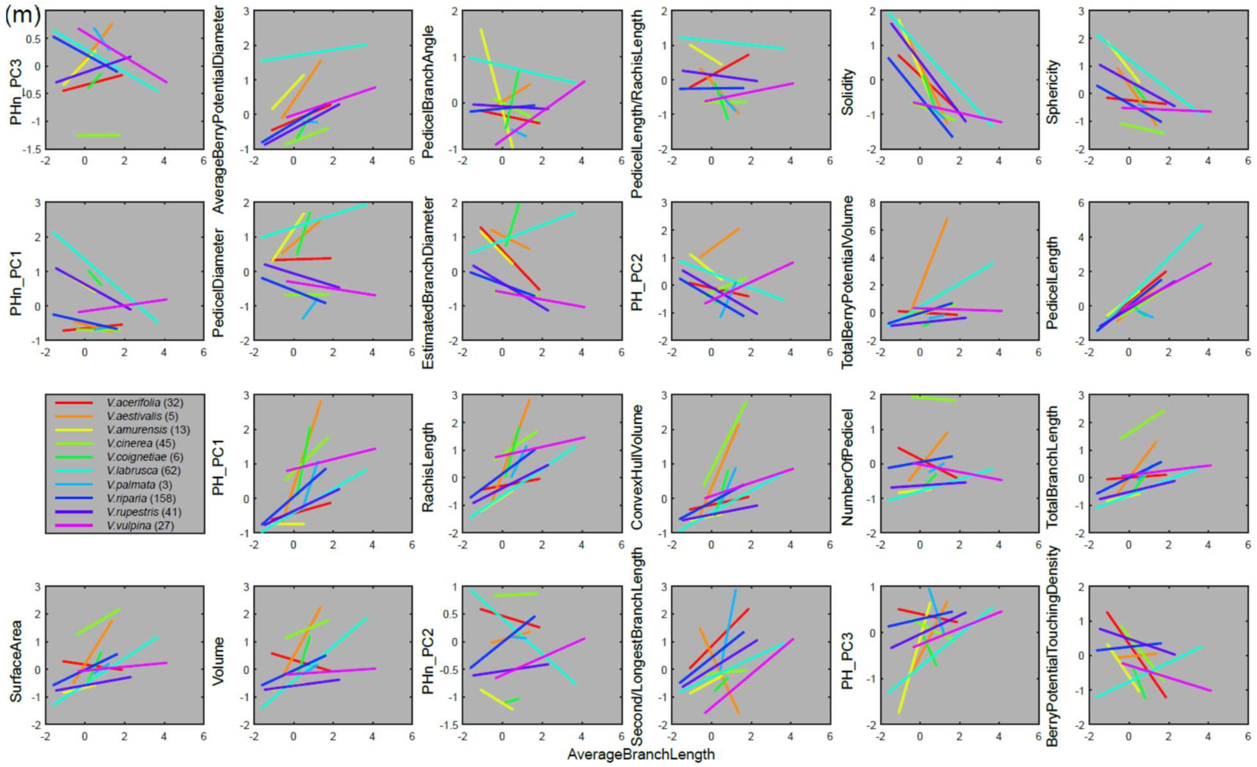

(m)
PHn_PC3
AverageBerryPotentialDiameter
PedicelBranchAngle
PedicelLength/RachisLength
Solidity
Sphericity
PHn_PC1
PedicelDiameter
EstimatedBranchDiameter
PH_PC2
TotalBerryPotentialVolume
PedicelLength
V.acerifolia (32)
V.aestivalis (5)
V.amurensis (13)
V.cinerea (45)
V.coignetiae (6)
V.labrusca (62)
V.palmata (3)
V.riparia (158)
V.rupestris (41)
V.vulpina (27)
PH_PC1
RachisLength
ConvexHullVolume
NumberOfPedicel
TotalBranchLength
SurfaceArea
Volume
PHn_PC2
Second/LongestBranchLength
PH_PC3
BerryPotentialTouchingDensity
AverageBranchLength


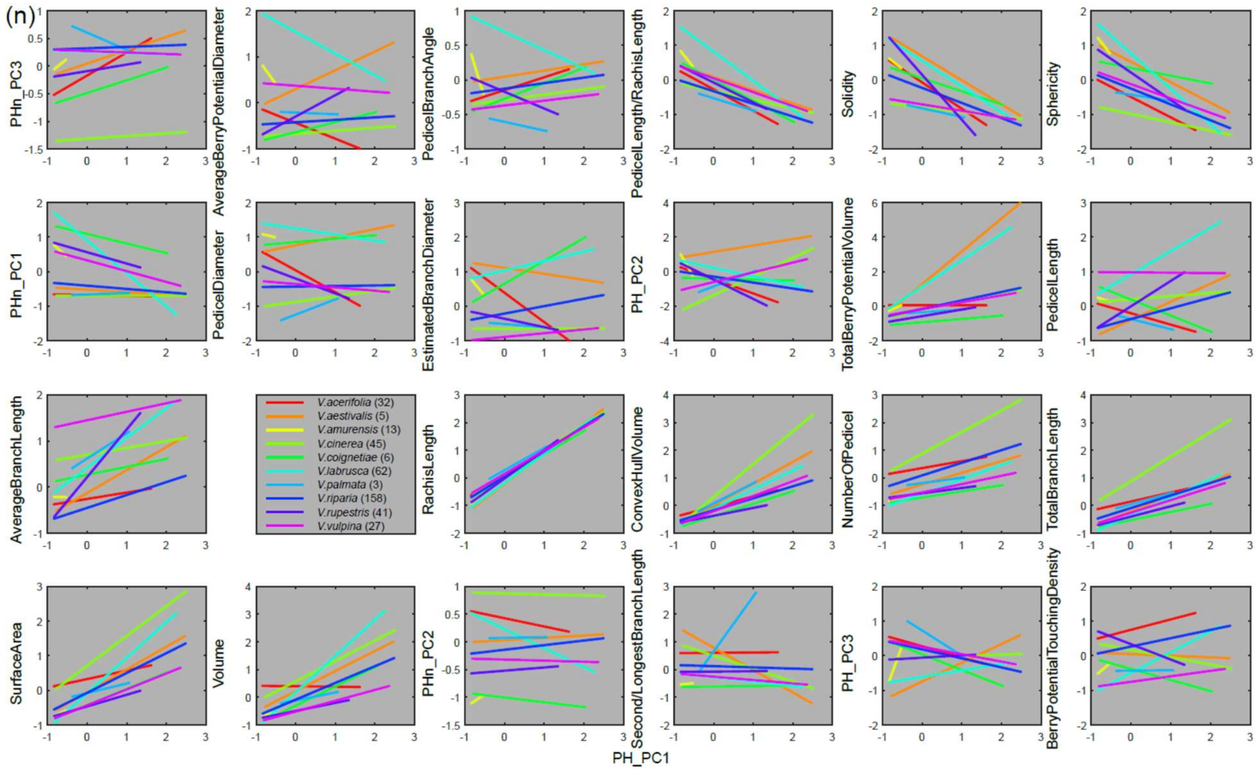

(n)
PHn_PC3
AverageBerryPotentialDiameter
PedicelBranchAngle
PedicelLength/RachisLength
Solidity
Sphericity
PHn_PC1
PedicelDiameter
EstimatedBranchDiameter
PH_PC2
TotalBerryPotentialVolume
PedicelLength
AverageBranchLength
V.acerifolia (32)
V.aestivalis (5)
V.amurensis (13)
V.cinerea (45)
V.coignetiae (6)
V.labrusca (62)
V.palmata (3)
V.riparia (158)
V.rupestris (41)
V.vulpina (27)
RachisLength
ConvexHullVolume
NumberOfPedicel
TotalBranchLength
SurfaceArea
Volume
PHn_PC2
Second/LongestBranchLength
PH_PC3
BerryPotentialTouchingDensity
PH_PC1

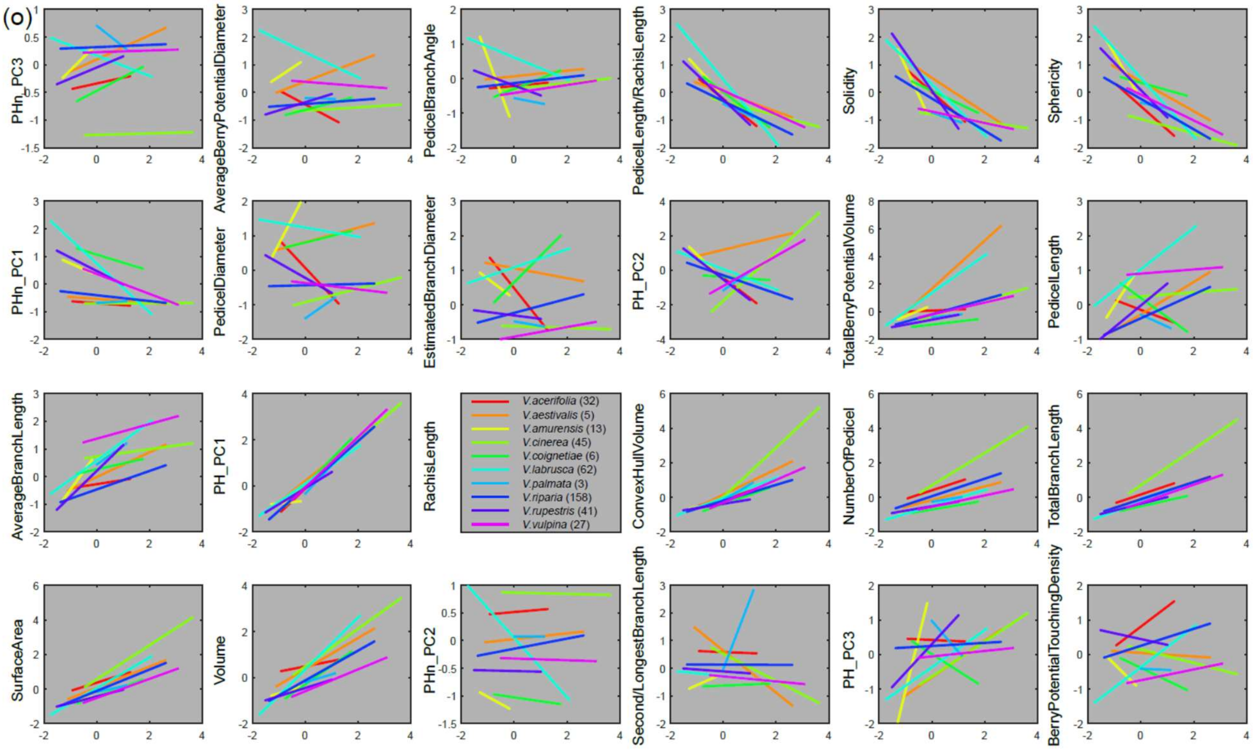
(o)
PHn_PC3
AveragBerryPotentialDiameter
PedicelBranchAngle
PedicelLength/RachisLength
Solidity
Sphericity
PHn_PC1
PedicelDiameter
EstimatedBranchDiameter
PH_PC2
TotalBerryPotentialVolume
PedicelLength
AverageBranchLength
PH_PC1
RachisLength
ConvexHullVolume
NumberOfPedicel
TotalBranchLength
SurfaceArea
Volume
PHn_PC2
Second/LongestBranchLength
PH_PC3
BerryPotentialTouchingDensity
V.acerifolia (32)
V.aestivalis (5)
V.amurensis (13)
V.cinerea (45)
V.coignetiae (6)
V.labrusca (62)
V.palmata (3)
V.riparia (158)
V.rupestris (41)
V.vulpina (27)

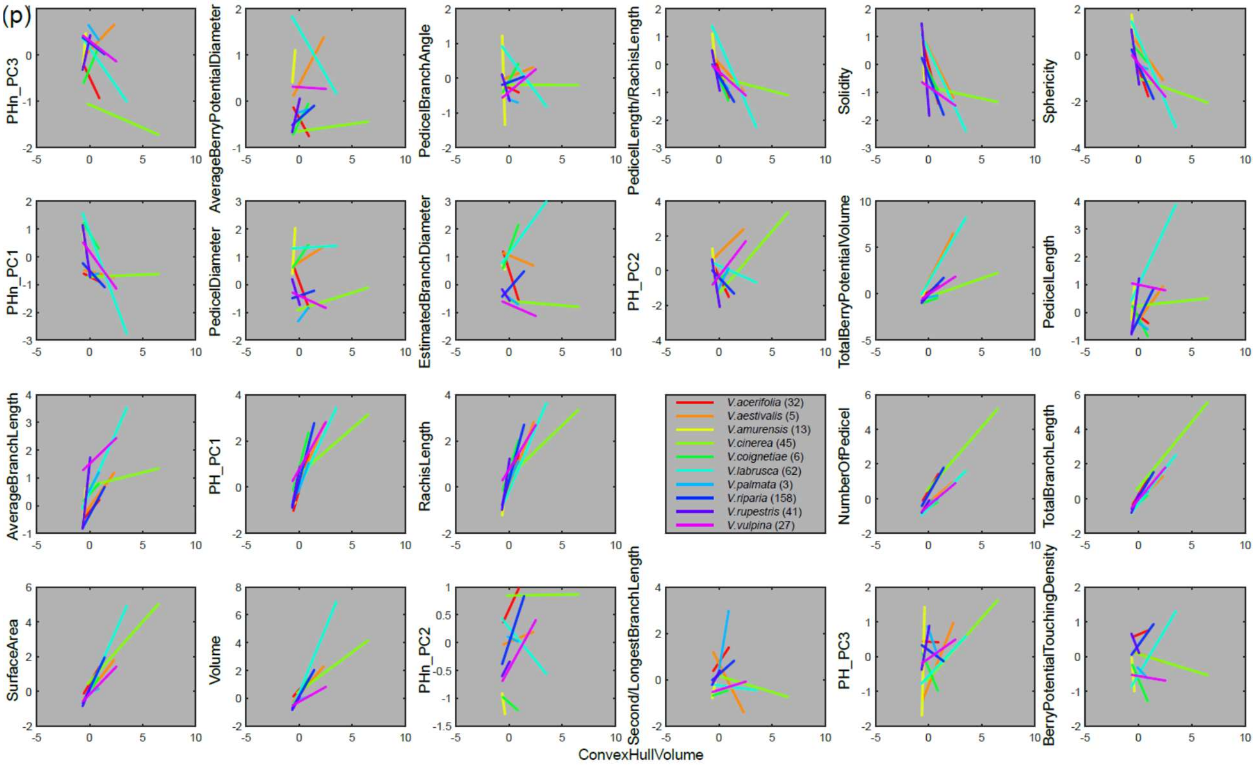
(p)
PHn_PC3
AverageBerryPotentialDiameter
PedicelBranchAngle
PedicelLength/RachisLength
Solidity
Sphericity
PHn_PC1
PedicelDiameter
EstimatedBranchDiameter
PH_PC2
TotalBerryPotentialVolume
PedicelLength
AverageBranchLength
PH_PC1
RachisLength
NumberOfPedicel
TotalBranchLength
SurfaceArea
Volume
PHn_PC2
Second/LongestBranchLength
PH_PC3
BerryPotentialTouchingDensity
V.acerifolia (32)
V.aestivalis (5)
V.amurensis (13)
V.cinerea (45)
V.coignetiae (6)
V.labrusca (62)
V.palmata (3)
V.riparia (158)
V.rupestris (41)
V.vulpina (27)
ConvexHullVolume

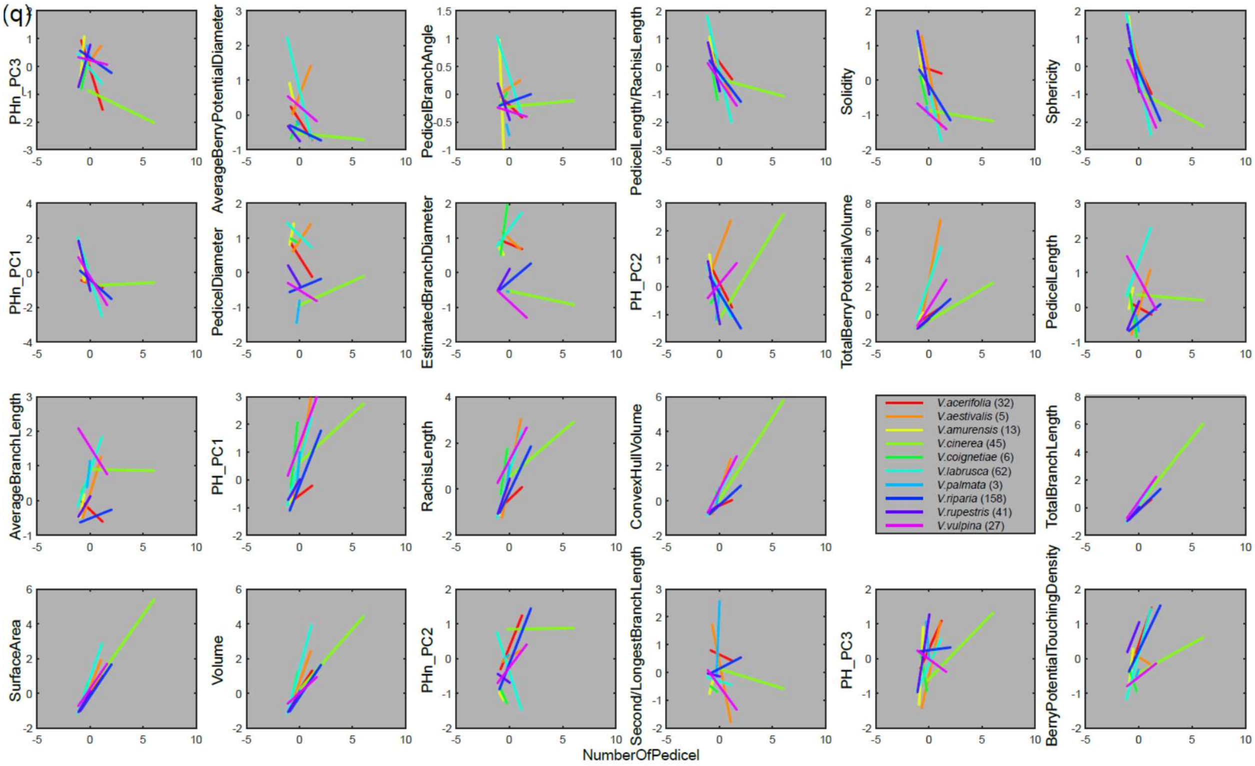
(q)
PHn_PC3
AverageBerryPotentialDiameter
PedicelBranchAngle
PedicelLength/RachisLength
Solidity
Sphericity
PHn_PC1
PedicelDiameter
EstimatedBranchDiameter
PH_PC2
TotalBerryPotentialVolume
PedicelLength
AverageBranchLength
PH_PC1
RachisLength
ConvexHullVolume
TotalBranchLength
SurfaceArea
Volume
PHn_PC2
Second/LongestBranchLength
PH_PC3
BerryPotentialTouchingDensity
NumberOfPedicel
V.acerifolia (32)
V.aestivalis (5)
V.amurensis (13)
V.cinerea (45)
V.coignetiae (6)
V.labrusca (62)
V.palmata (3)
V.riparia (158)
V.rupestris (41)
V.vulpina (27)

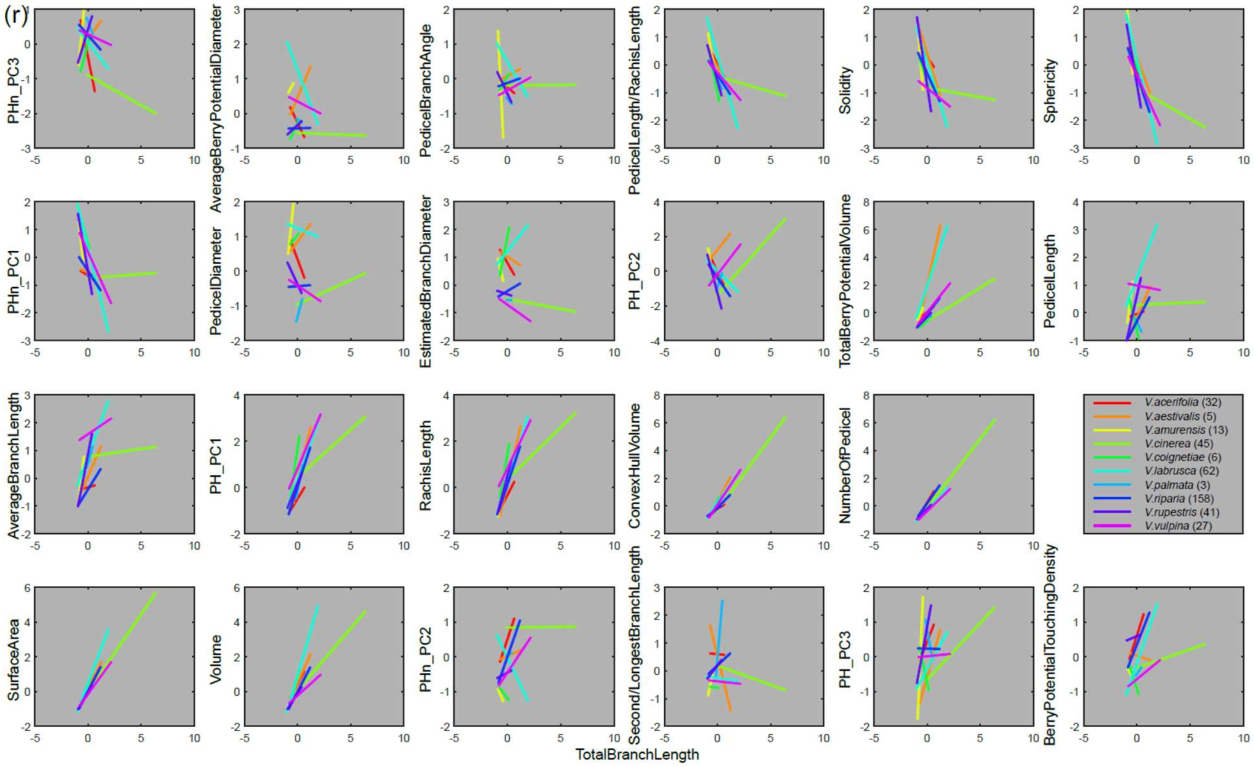
(r)
PHn_PC3
AverageBerryPotentialDiameter
PedicelBranchAngle
PedicelLength/RachisLength
Solidity
Sphericity
PHn_PC1
PedicelDiameter
EstimatedBranchDiameter
PH_PC2
TotalBerryPotentialVolume
PedicelLength
AverageBranchLength
PH_PC1
RachisLength
ConvexHullVolume
NumberOfPedicel
SurfaceArea
Volume
PHn_PC2
Second/LongestBranchLength
PH_PC3
BerryPotentialTouchingDensity
TotalBranchLength
V.acerifolia (32)
V.aestivalis (5)
V.amurensis (13)
V.cinerea (45)
V.coignetiae (6)
V.labrusca (62)
V.palmata (3)
V.riparia (158)
V.rupestris (41)
V.vulpina (27)

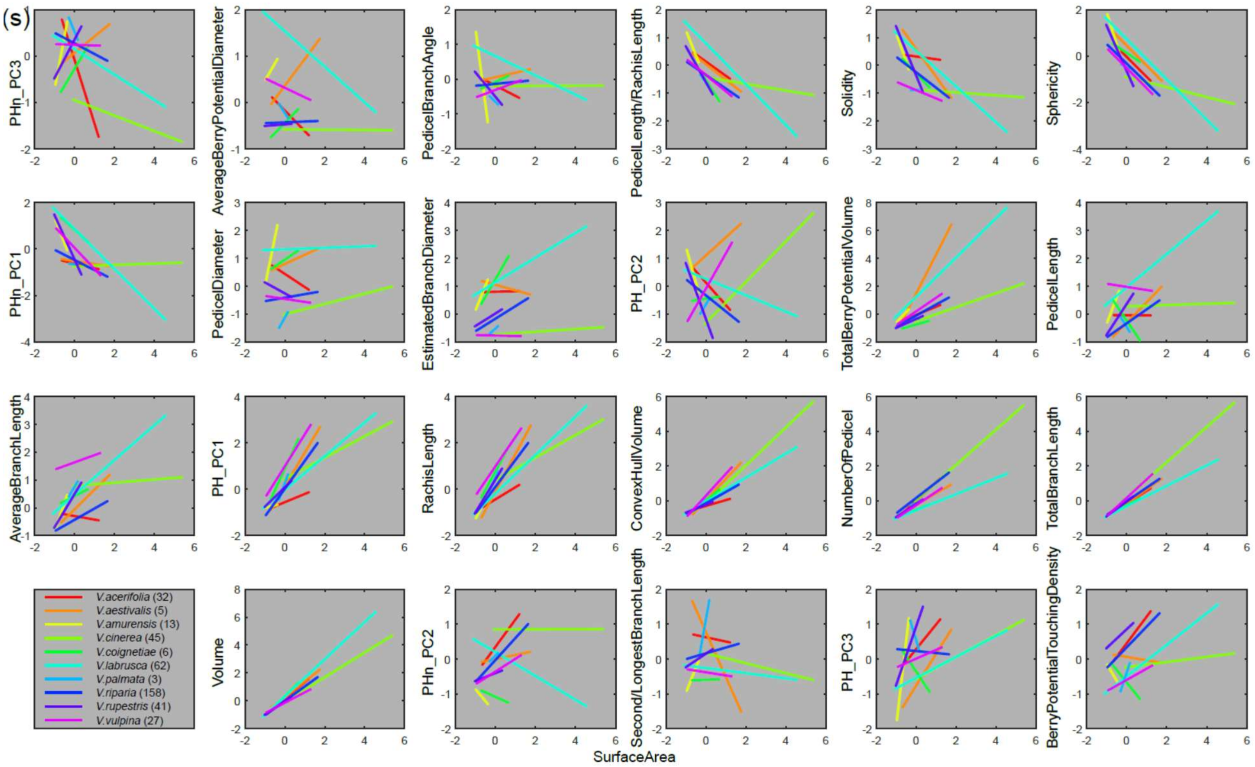
(s)
PHn_PC3
AverageBerryPotentialDiameter
PedicelBranchAngle
PedicelLength/RachisLength
Solidity
Sphericity
PHn_PC1
PedicelDiameter
EstimatedBranchDiameter
PH_PC2
TotalBerryPotentialVolume
PedicelLength
AverageBranchLength
PH_PC1
RachisLength
ConvexHullVolume
NumberOfPedicel
TotalBranchLength
V.acerifolia (32)
V.aestivalis (5)
V.amurensis (13)
V.cinerea (45)
V.coignetiae (6)
V.labrusca (62)
V.palmata (3)
V.riparia (158)
V.rupestris (41)
V.vulpina (27)
Volume
PHn_PC2
Second/LongestBranchLength
PH_PC3
BerryPotentialTouchingDensity
SurfaceArea

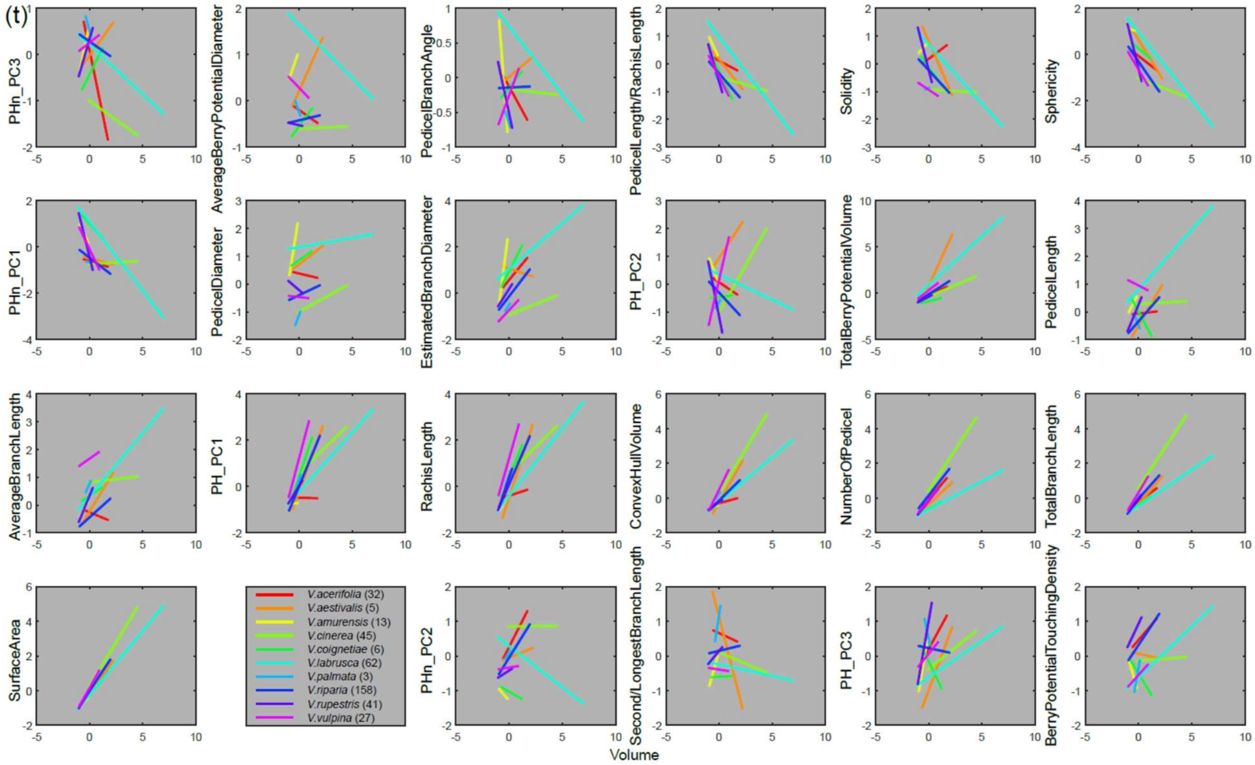
(t)
PHn_PC3
AverageBerryPotentialDiameter
PedicelBranchAngle
PedicelLength/RachisLength
Solidity
Sphericity
PHn_PC1
PedicelDiameter
EstimatedBranchDiameter
PH_PC2
TotalBerryPotentialVolume
PedicelLength
AverageBranchLength
PH_PC1
RachisLength
ConvexHullVolume
NumberOfPedicel
TotalBranchLength
SurfaceArea
V.acerifolia (32)
V.aestivalis (5)
V.amurensis (13)
V.cinerea (45)
V.coignetiae (6)
V.labrusca (62)
V.palmata (3)
V.riparia (158)
V.rupestris (41)
V.vulpina (27)
PHn_PC2
Second/LongestBranchLength
PH_PC3
BerryPotentialTouchingDensity
Volume

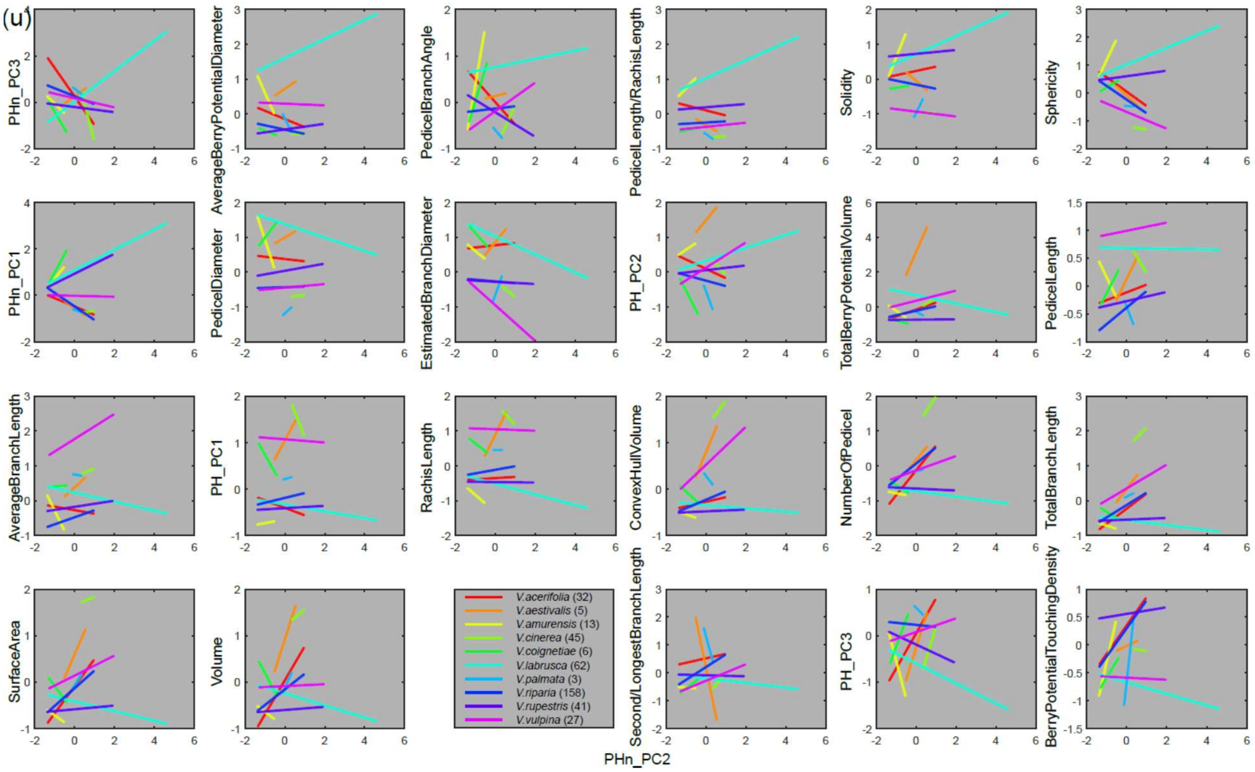
(u)
PHn_PC3
AverageBerryPotentialDiameter
PedicelBranchAngle
PedicelLength/RachisLength
Solidity
Sphericity
PHn_PC1
PedicelDiameter
EstimatedBranchDiameter
PH_PC2
TotalBerryPotentialVolume
PedicelLength
AverageBranchLength
PH_PC1
RachisLength
ConvexHullVolume
NumberOfPedicel
TotalBranchLength
SurfaceArea
Volume
Second/LongestBranchLength
PH_PC3
BerryPotentialTouchingDensity
V.acerifolia (32)
V.aestivalis (5)
V.amurensis (13)
V.cinerea (45)
V.coignetiae (6)
V.labrusca (62)
V.palmata (3)
V.riparia (158)
V.rupestris (41)
V.vulpina (27)
PHn_PC2

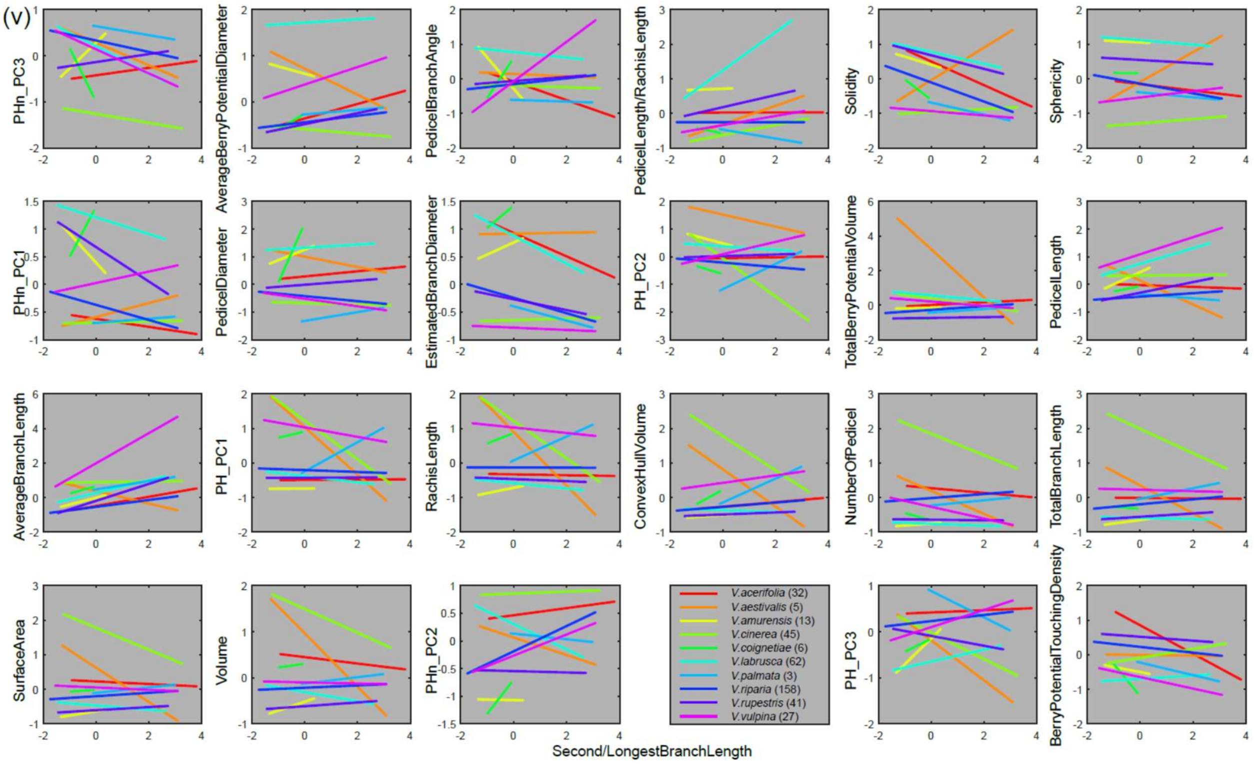
(v)
PHn_PC3
AverageBerryPotentialDiameter
PedicelBranchAngle
PedicelLength/RachisLength
Solidity
Sphericity
PHn_PC1
PedicelDiameter
EstimatedBranchDiameter
PH_PC2
TotalBerryPotentialVolume
PedicelLength
AverageBranchLength
PH_PC1
RachisLength
ConvexHullVolume
NumberOfPedicel
TotalBranchLength
SurfaceArea
Volume
PHn_PC2
V.acerifolia (32)
V.aestivalis (5)
V.amurensis (13)
V.cinerea (45)
V.coignetiae (6)
V.labrusca (62)
V.palmata (3)
V.riparia (158)
V.rupestris (41)
V.vulpina (27)
PH_PC3
BerryPotentialTouchingDensity
Second/LongestBranchLength

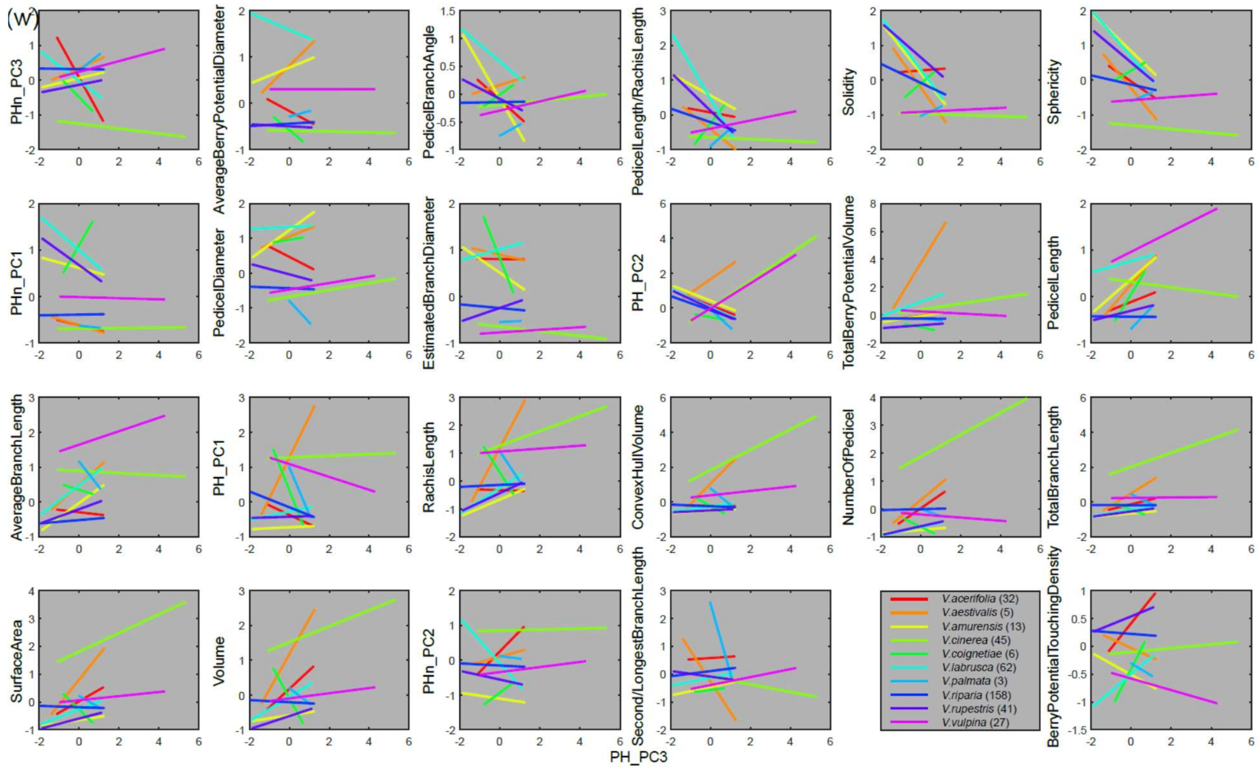
(w)
PHn_PC3
AverageBerryPotentialDiameter
PedicelBranchAngle
PedicelLength/RachisLength
Solidity
Sphericity
PHn_PC1
PedicelDiameter
EstimatedBranchDiameter
PH_PC2
TotalBerryPotentialVolume
PedicelLength
AverageBranchLength
PH_PC1
RachisLength
ConvexHullVolume
NumberOfPedicel
TotalBranchLength
SurfaceArea
Volume
PHn_PC2
Second/LongestBranchLength
BerryPotentialTouchingDensity
V.acerifolia (32)
V.aestivalis (5)
V.amurensis (13)
V.cinerea (45)
V.coignetiae (6)
V.labrusca (62)
V.palmata (3)
V.riparia (158)
V.rupestris (41)
V.vulpina (27)
PH_PC3

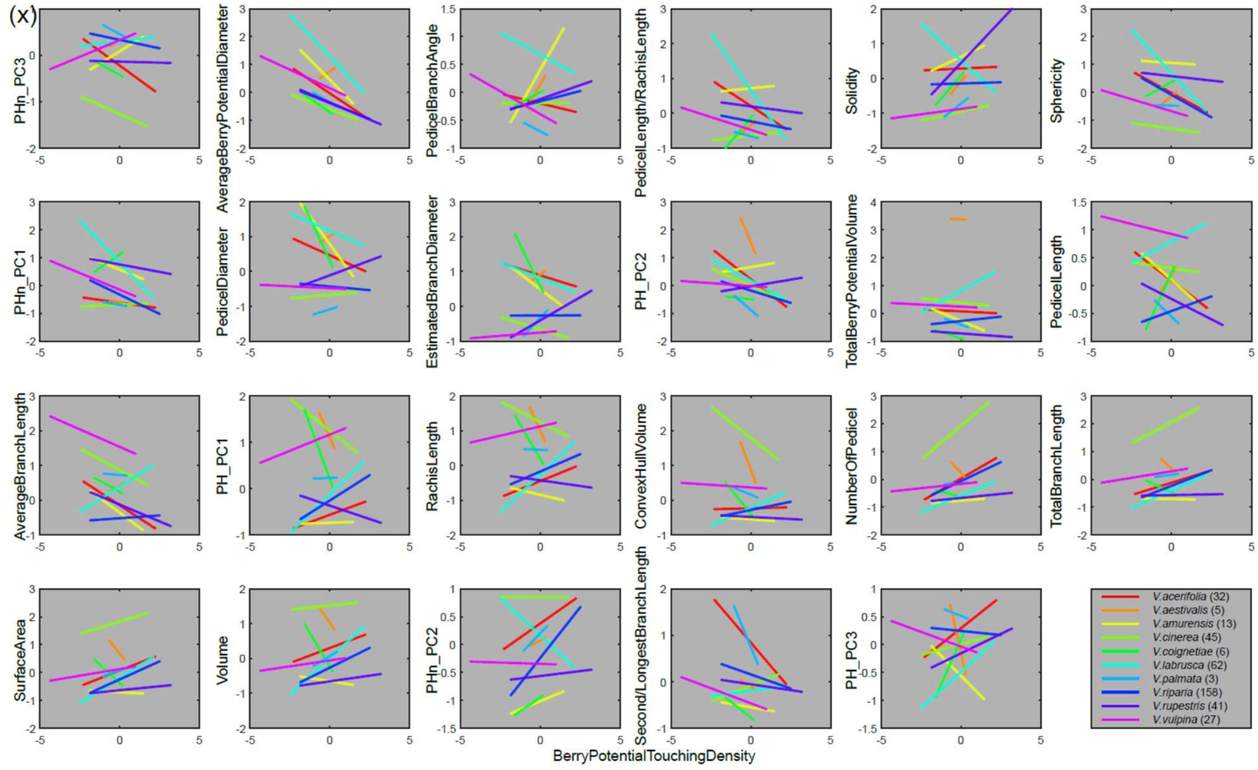
(x)
PHn_PC3
AverageBerryPotentialDiameter
PedicelBranchAngle
PedicelLength/RachisLength
Solidity
Sphericity
PHn_PC1
PedicelDiameter
EstimatedBranchDiameter
PH_PC2
TotalBerryPotentialVolume
PedicelLength
AverageBranchLength
PH_PC1
RachisLength
ConvexHullVolume
NumberOfPedicel
TotalBranchLength
SurfaceArea
Volume
PHn_PC2
Second/LongestBranchLength
PH_PC3
V.acerifolia (32)
V.aestivalis (5)
V.amurensis (13)
V.cinerea (45)
V.coignetiae (6)
V.labrusca (62)
V.palmata (3)
V.riparia (158)
V.rupestris (41)
V.vulpina (27)
BerryPotentialTouchingDensity

**Fig. S6** Pairwise species classification. On the upper right triangle, each purple square shows the leave one out classification accuracy rate based on PCA and LDA for the two *Vitis* species naming for the corresponding row and column. The colorbar indicating the accuracy rate. More purple means more accurate in classification. On the lower left triangle, the 24 gray scaled boxes showing the weight/ loading for the 24 traits to best distinguish the two *Vitis* species naming for the corresponding row and column. The trait name for each box is listed in the bottom enlarged example for *V. coignetiae* v.s. *V. vulpina* with the colorbar. Darker represents larger weight. The rachis examples for *V. coignetiae* and *V. vulpina* are shown in the right panel.

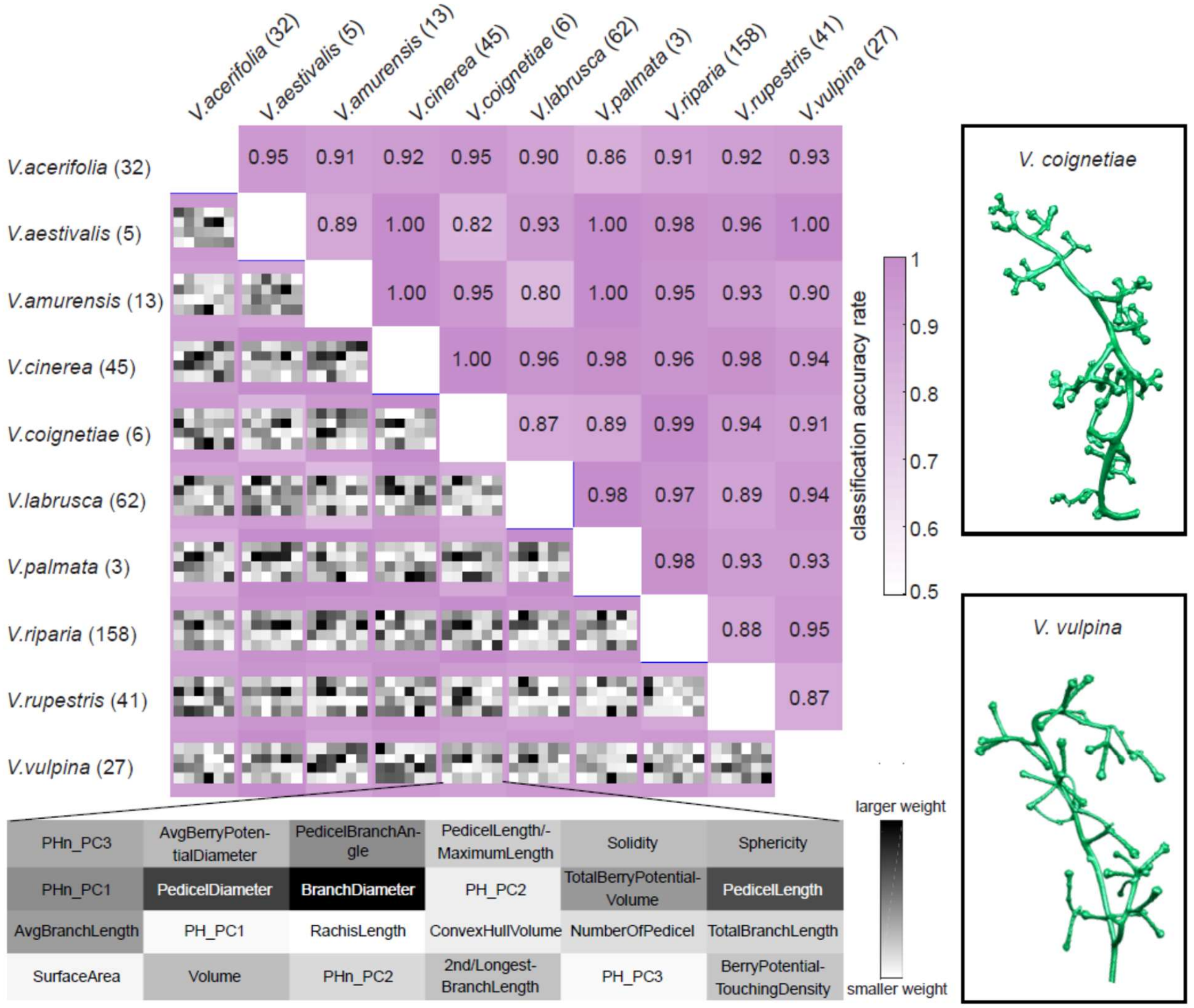

**Table S1** Trait description and calculation

| Trait | Unit | Description and Calculation |
|---|---|---|
| Volume | $mm^3$ | Larger value of the volume measures larger size of the infloresence architecture. It is the total number of voxels comprising the infloresence architecture multiplying by the $unit^3$ of each voxel side |
| ConvexHullVolume | $mm^3$ | Larger value of the convex hull volume measures larger space that the infloresence architecture occupied. It is calculated from the volume of the smallest convex set that contains all the infloresence architecture voxels |
| Solidity | NA | Larger value of the solidity usually shows densier or more compact of the infloresence architecture occupied in the space. It is calculated as the ratio Volume/ConvexHullVolume |
| SurfaceArea | $mm^2$ | The area of the surface of the infloresence architecture. Larger size of the infloresence architecture usually has larger value of the surface area. It is the sum of area of the triangle faces on the surface mesh |
| Sphericity | NA | Sphericity measures how closely the shape of infloresence architecture is to a sphere. Larger value usually has relatively thick or less branches. It is calculated by the formula $6\pi(\text{Volume})^2/(\text{SurfaceArea})^3$ |
| TotalBranchLength | mm | TotalBranchLength is the sum of the length of all branches. Larger value usually shows more branches or longer branches. We first compute persistent barcode with geodesic distance to the base in which each bar represents a branch with its length information recorded. Then we sum the length of all bars. |
| RachisLength | mm | RachisLength is the length of the main branch/rachis. It is the length of the longestbars in the barcode |
| AvgBranchLength | mm | Larger value of the average branch length usually means overall longer branches in the infloresence architecture. It is calculated by the ratio TotalBranchLength/number of branches where number of branches is equal to the number of bars in the barcode |
| 2nd/LongestBranchLength | NA | The longest branch length usually is the length of the rachis. If there is a shoulder/first primary branch in the rachi, the second longest branch length is usually the length of the shoulder. The value can be used to imply the possible shoulder structure in the infloresence architecture |
| BranchDiameter | mm | Larger value shows thicker branches. Instead direcly measuring from the rachis branches, the branch diameter is estimated using the formular 2√( Volume/(π*TotalBranchLength ) |
| NumberOfPedicel | no. | The number of pedicels which are the branches might hold berries. It is almost similar to the number of tips expect a few exception |
| PedicelLength | mm | It measures the average length of pedicels. It is calculated by the maximum value in the density estimator of the branch length as we assume the pedicels appear in the highest frequency in all types of branches in the infloresence architecture |
| PedicelDiameter | mm | It measures the average diameter of pedicels. Larger value means thicker of the pedicels. Each diameter is derived from the Dynamic Roots software. We computed the mean for the subset of the detected pedicels |
| PedicelBranchAngle | deg | It measures the average angle of pedicel to its parent branch. Larger value means more open of the pedicels. Each angle is derived from the Dynamic Roots software. The PedicelBranchAngle is the mean for the subset of the detected pedicels |
| PedicelLength/RachisLength | NA | The ratio between the PedicelLength and RachisLength. Smaller value ususally shows relatively longer rachis to the pedicel |
| PH_PC1 | NA | First few PCs of persistent homology can reveal the most variance in the topological and geometric branch strucuture. This traits is the PC1 of persistent homology with geodesic distance to the base. |
| PH_PC2 | NA | This traits is the PC2 of persistent homology with geodesic distance to the base. |
| PH_PC3 | NA | This traits is the PC3 of persistent homology with geodesic distance to the base. |
| PHn_PC1 | NA | This traits is the PC1 of persistent homology with geodesic distance to the base after scaling the infloresence architecture so that the total branch length is 1 |
| PHn_PC2 | NA | This is the PC2 of persistent homology with geodesic distance to the base after scaling the infloresence architecture so that the total branch length is 1 |
| PHn_PC3 | NA | This is the PC3 of persistent homology with geodesic distance to the base after scaling the infloresence architecture so that the total branch length is 1 |
| TotalBerryPotentialVolume | $mm^3$ | The total number of voxels comprising all berry potential. The berry potential is the simulated balls at the tip of the pedicel that stop growing until it either touches another ball, touches branch, or reach the size specifically for different species. This indirectly reflects the comprehensive relationship among number of pedicel, pedicel mutural angles, and pedicle length, exploring the space limited by the infloresence architecture |
| AvgBerryPotentialDiameter | mm | This average diameter reveals the size of the berry potential. Larger value ususally is ususally due to the sparse of the pedicel, larger pedicel mutural angle, and longer pedicel length, or larger size upper bound |
| BerryPotentialTouchingDensity | NA | Smaller number usually shows more loose bunch. It is calculated by number of Berry Potential touching/number of Berry Potential. We account the touching either by touching with other Berry Potential or touching with any part of the inflorescence. |

**Table S2** Trait variance for each species

| | V_acerifolia | V_aestivalis | V_amurensis | V_cinerea | V_coignetiae | V_labrusca | V_palmata | V_riparia | V_rupestris | V_vulpina |
|---|---|---|---|---|---|---|---|---|---|---|
| Volume | 0.36163 | **1.6499** | **0.051843** | 1.4951 | 0.557 | 1.5391 | 0.094832 | 0.32651 | 0.11575 | 0.24412 |
| ConvexHullVolume | 0.07061 | 2.0185 | **0.0064221** | **2.6835** | 0.28557 | 0.42078 | 0.34163 | 0.1467 | 0.035538 | 0.71689 |
| Solidity | 0.41036 | 1.385 | 0.88764 | **0.051833** | 0.19146 | 1.4162 | 0.11597 | 0.46664 | **1.4728** | 0.092489 |
| SurfaceArea | 0.22079 | 1.3023 | **0.025379** | **1.6971** | 0.27295 | 0.85053 | 0.068723 | 0.28847 | 0.12104 | 0.36081 |
| Sphericity | 0.3203 | 1.014 | 0.59095 | 0.12997 | 0.1174 | **1.2753** | **0.018163** | 0.34116 | 0.63822 | 0.37292 |
| TotalBranchLength | 0.13838 | 0.86932 | **0.015806** | **1.9434** | 0.10187 | 0.30946 | 0.095864 | 0.19231 | 0.099153 | 0.45407 |
| RachisLength | 0.24198 | **3.4664** | **0.10882** | 0.83641 | 0.79229 | 0.79693 | 0.38751 | 0.54 | 0.4435 | 0.82968 |
| AvgBranchLength | 0.27085 | 0.79809 | 0.30275 | 0.23008 | **0.06206** | 1.1369 | 0.17938 | 0.33645 | 1.1262 | **1.3784** |
| 2nd/LongestBranchLength | 0.73711 | **3.3919** | 0.29573 | 0.75725 | **0.10645** | 1.0339 | 2.7824 | 0.96208 | 1.0307 | 0.95123 |
| BranchDiameter | 0.694 | 0.2073 | 1.3529 | 0.33466 | 0.57055 | **1.5063** | **0.14978** | 0.44582 | 0.47249 | 0.4452 |
| NumberOfPedicel | 0.27597 | 0.56436 | **0.011567** | **1.9936** | 0.063841 | 0.22395 | 0.022477 | 0.28112 | 0.08554 | 0.31066 |
| PedicelLength | 0.49891 | 0.95011 | 0.25795 | 0.25323 | 0.50247 | **2.3012** | **0.097169** | 0.41766 | 0.79138 | 0.88714 |
| PedicleDiameter | 0.539 | 0.2017 | 0.7756 | 0.2451 | 0.916 | **1.2044** | **0.1259** | 0.3115 | 0.5061 | 0.3993 |
| PedicelBranchAngle | 0.4172 | 0.1885 | 1.0819 | 0.3737 | 0.5776 | **1.9396** | **0.023** | 0.7288 | 1.1074 | 0.689 |
| PedicelLength/RachisLength | 0.23656 | 0.41967 | 0.14525 | **0.076333** | 0.48704 | **3.0349** | 0.081875 | 0.20494 | 0.73452 | 0.25544 |
| TotalBerryPotentialVolume | 0.15897 | **11.22** | 0.11083 | 0.49162 | 0.046818 | 2.2124 | **0.038094** | 0.25665 | 0.072486 | 0.55551 |
| AvgBerryPotentialDiameter | 0.16317 | 0.58725 | 0.31597 | **0.039747** | 0.072778 | **1.4938** | 0.073299 | 0.13597 | 0.23024 | 0.18297 |
| BerryPotentialTouchingDensity | 0.75372 | **0.17144** | 0.75605 | 0.69907 | 0.36946 | 1.0743 | 0.60217 | 0.66532 | 1.1486 | **1.2247** |
| PH_PC1 | 0.3184 | **2.8847** | **0.0096105** | 0.90197 | 1.0849 | 0.59902 | 0.59107 | 0.60222 | 0.29921 | 1.0759 |
| PH_PC2 | 0.53731 | 0.96919 | **0.30062** | **2.214** | 0.71213 | 0.69814 | 0.58958 | 0.71487 | 0.74374 | 1.6619 |
| PH_PC3 | 0.38162 | 1.3361 | 1.0924 | **1.5948** | 0.33731 | 0.92022 | **0.31658** | 0.69013 | 1.0904 | 1.244 |
| PHn_PC1 | 0.068755 | 0.063323 | 0.44906 | **0.0037512** | 0.38184 | **1.4648** | 0.0069755 | 0.27759 | 0.97863 | 0.76306 |
| PHn_PC2 | 0.29708 | 0.15429 | 0.057095 | **0.015133** | 0.12193 | **2.8024** | 0.054566 | 0.47877 | 0.68431 | 0.61346 |
| PHn_PC3 | 0.79849 | 0.23151 | 0.43607 | 0.29702 | 0.40699 | **2.5777** | **0.077309** | 0.36682 | 0.43925 | 0.49579 |
| Summation | 8.911165 | **36.044853** | 9.4382126 | 19.358377 | 9.138707 | **32.83223** | **6.9343165** | 10.1785 | 14.467197 | 16.204639 |

**Table S3** Trait loadings for two species classification

| Species 1 | Species 2 | PHn_PC3 | AvgBerryP | PedicelB | PedicelLe | Solidity | Sphericit | PHn_PC1 | PedicleD | BranchDi | PH_PC2 | TotalBerr | PedicelLe | AverageE | PH_PC1 | RachisLe | ConvexHu | NumberC | TotalBra | SurfaceA | Volume | PHn_PC2 | 2nd/Long | PH_PC3 | BerryPote |
|---|---|---|---|---|---|---|---|---|---|---|---|---|---|---|---|---|---|---|---|---|---|---|---|---|---|
| V.acerifolia | V.aestivalis | 0.23556 | 0.1723 | 0.09009 | 0.00957 | 0.01406 | 0.1002 | 0.02763 | 0.15876 | 0.07839 | 0.27261 | 0.32813 | 0.03376 | 0.08463 | 0.15131 | 0.10692 | 0.10011 | 0.09172 | 0.01948 | 0.02445 | 0.0048 | 0.1348 | 0.13381 | 0.14485 | 0.16373 |
| V.acerifolia | V.amurensi | 0.00693 | 0.24916 | 0.07481 | 0.07484 | 0.06949 | 0.16364 | 0.43133 | 0.2569 | 0.16156 | 0.06772 | 0.02911 | 0.23477 | 0.08326 | 0.02745 | 0.00584 | 0.07863 | 0.20424 | 0.12092 | 0.17179 | 0.22782 | 0.4751 | 0.73544 | 0.05461 | 0.13469 |
| V.acerifolia | V.cinerea | 0.05297 | 0.054351 | 0.02679 | 0.11435 | 0.26315 | 0.17428 | 0.00413 | 0.25069 | 0.33625 | 0.23276 | 0.05019 | 0.0711 | 0.23364 | 0.29247 | 0.2158 | 0.14181 | 0.05814 | 0.14682 | 0.06204 | 0.03018 | 0.02704 | 0.06912 | 0.26702 | 0.15156 |
| V.acerifolia | V.coignetia | 0.25482 | 0.050582 | 0.23522 | 0.08568 | 0.09764 | 0.12515 | 0.5581 | 0.09174 | 0.21347 | 0.16881 | 0.21249 | 0.22664 | 0.31061 | 0.42029 | 0.35836 | 0.08563 | 0.17486 | 0.01566 | 0.01436 | 0.09616 | 0.45858 | 0.60759 | 0.17617 | 0.44157 |
| V.acerifolia | V.labrusca | 0.0853 | 0.41028 | 0.22024 | 0.1259 | 0.12918 | 0.12801 | 0.33013 | 0.11609 | 0.02941 | 0.03896 | 0.23527 | 0.07764 | 0.15615 | 0.07726 | 0.07218 | 0.00227 | 0.17295 | 0.08083 | 0.09656 | 0.10817 | 0.18058 | 0.19726 | 0.05629 | 0.27325 |
| V.acerifolia | V.palmata | 0.13511 | 0.02004 | 0.11933 | 0.04964 | 0.27839 | 0.07589 | 0.05821 | 0.30425 | 0.33829 | 0.13616 | 0.05039 | 0.07774 | 0.23097 | 0.19981 | 0.1711 | 0.09501 | 0.06934 | 0.03546 | 0.03941 | 0.12037 | 0.13196 | 0.09474 | 0.0597 | 0.25964 |
| V.acerifolia | V.riparia | 0.31302 | 0.13733 | 0.1165 | 0.16689 | 0.27543 | 0.05237 | 0.14044 | 0.40935 | 0.5685 | 0.09749 | 0.16772 | 0.23481 | 0.07916 | 0.25731 | 0.17717 | 0.00958 | 0.081 | 0.04546 | 0.14373 | 0.26407 | 0.32368 | 0.17003 | 0.24252 | 0.02271 |
| V.acerifolia | V.rupestris | 0.11911 | 0.12939 | 0.29226 | 0.00231 | 0.07489 | 0.15552 | 0.45459 | 0.18003 | 0.50656 | 0.17092 | 0.35314 | 0.06926 | 0.21703 | 0.02555 | 0.02506 | 0.10713 | 0.32777 | 0.17367 | 0.29568 | 0.4187 | 0.36735 | 0.28908 | 0.0286 | 0.28008 |
| V.acerifolia | V.vulpina | 0.08915 | 0.12977 | 0.12519 | 0.02535 | 0.18403 | 0.02466 | 0.23792 | 0.13581 | 0.30882 | 0.01536 | 0.01507 | 0.34812 | 0.3451 | 0.30966 | 0.28177 | 0.05551 | 0.10353 | 0.01787 | 0.05535 | 0.11489 | 0.25268 | 0.58798 | 0.08121 | 0.30263 |
| V.aestivalis | V.amurensi | 0.00044 | 0.063502 | 0.1871 | 0.03612 | 0.08104 | 0.00398 | 0.07187 | 0.13116 | 0.04271 | 0.1653 | 0.26704 | 0.08056 | 0.04767 | 0.16193 | 0.10391 | 0.1164 | 0.07944 | 0.07787 | 0.09876 | 0.12438 | 0.11803 | 0.02954 | 0.14373 | 0.1504 |
| V.aestivalis | V.cinerea | 0.33089 | 0.32921 | 0.08303 | 0.10992 | 0.29608 | 0.3072 | 0.03764 | 0.56982 | 0.58329 | 0.34173 | 0.91324 | 0.07155 | 0.20802 | 0.17546 | 0.18245 | 0.24618 | 0.2047 | 0.27974 | 0.04938 | 0.20353 | 0.20096 | 0.17094 | 0.00027 | 0.24096 |
| V.aestivalis | V.coignetia | 0.06427 | 0.17786 | 0.03418 | 0.22143 | 0.23173 | 0.08043 | 0.28807 | 0.07985 | 0.15424 | 0.46657 | 0.59946 | 0.08473 | 0.10783 | 0.27644 | 0.27374 | 0.00474 | 0.05767 | 0.01075 | 0.03287 | 0.06943 | 0.25228 | 0.28792 | 0.01423 | 0.22063 |
| V.aestivalis | V.labrusca | 0.01407 | 0.23961 | 0.19424 | 0.02844 | 0.10714 | 0.03649 | 0.16119 | 0.05834 | 0.01901 | 0.15431 | 0.1021 | 0.13723 | 0.11291 | 0.12784 | 0.05806 | 0.12582 | 0.09643 | 0.09343 | 0.14333 | 0.18518 | 0.09109 | 0.02184 | 0.15653 | 0.09483 |
| V.aestivalis | V.palmata | 0.21564 | 0.24374 | 0.33476 | 0.31671 | 0.71373 | 0.45201 | 0.04871 | 0.85263 | 0.75494 | 0.76273 | 0.74339 | 0.02773 | 0.37467 | 0.13041 | 0.33242 | 0.12948 | 0.0606 | 0.16076 | 0.0261 | 0.11657 | 0.00161 | 0.17955 | 0.58634 | 0.26905 |
| V.aestivalis | V.riparia | 0.05748 | 0.28503 | 0.21048 | 0.14212 | 0.16252 | 0.12347 | 0.11569 | 0.38368 | 0.34463 | 0.56841 | 0.58721 | 0.15508 | 0.13892 | 0.04425 | 0.03643 | 0.14619 | 0.04662 | 0.00746 | 0.07005 | 0.16477 | 0.12206 | 0.05723 | 0.06929 | 0.25064 |
| V.aestivalis | V.rupestris | 0.10742 | 0.10721 | 0.17585 | 0.06876 | 0.07516 | 0.04269 | 0.28668 | 0.23694 | 0.2782 | 0.39065 | 0.52127 | 0.13827 | 0.14007 | 0.1072 | 0.02673 | 0.17219 | 0.12013 | 0.09282 | 0.15967 | 0.22213 | 0.02004 | 0.29317 | 0.1626 | 0.24035 |
| V.aestivalis | V.vulpina | 0.11805 | 0.15365 | 0.25839 | 0.05842 | 0.25007 | 0.22161 | 0.00841 | 0.40584 | 0.43873 | 0.24816 | 0.61704 | 0.34513 | 0.31732 | 0.18404 | 0.20692 | 0.03988 | 0.04085 | 0.05176 | 0.04351 | 0.17565 | 0.12997 | 0.25878 | 0.11164 | 0.00954 |
| V.amurensi | V.cinerea | 0.13596 | 0.20918 | 0.13927 | 0.22277 | 0.34971 | 0.42099 | 0.27543 | 0.40192 | 0.36105 | 0.46622 | 0.07072 | 0.07532 | 0.26622 | 0.25202 | 0.23595 | 0.04764 | 0.08558 | 0.12771 | 0.08248 | 0.03546 | 0.33585 | 0.1334 | 0.08031 | 0.02273 |
| V.amurensi | V.coignetia | 0.1791 | 0.32673 | 0.18989 | 0.33469 | 0.22974 | 0.21371 | 0.15336 | 0.16331 | 0.01855 | 0.28564 | 0.17188 | 0.16869 | 0.08863 | 0.44337 | 0.40149 | 0.13838 | 0.06596 | 0.10449 | 0.15966 | 0.21123 | 0.04901 | 0.07286 | 0.01646 | 0.06486 |
| V.amurensi | V.labrusca | 0.1369 | 0.28839 | 0.14916 | 0.02969 | 0.00721 | 0.05267 | 0.14628 | 0.06962 | 0.1455 | 0.02567 | 0.18836 | 0.12558 | 0.09521 | 0.03557 | 0.04877 | 0.0399 | 0.01444 | 0.01301 | 0.04639 | 0.08009 | 0.23976 | 0.06823 | 0.03619 | 0.13191 |
| V.amurensi | V.palmata | 0.08174 | 0.25649 | 0.05189 | 0.23575 | 0.10261 | 0.16172 | 0.26438 | 0.50047 | 0.08777 | 0.10937 | 0.06286 | 0.24029 | 0.1447 | 0.28484 | 0.27032 | 0.21398 | 0.1405 | 0.1887 | 0.14846 | 0.14701 | 0.26194 | 0.55828 | 0.05552 | 0.04131 |
| V.amurensi | V.riparia | 0.2202 | 0.44434 | 0.37738 | 0.33841 | 0.13753 | 0.25656 | 0.21728 | 0.54982 | 0.34715 | 0.05432 | 0.18105 | 0.57602 | 0.36445 | 0.14192 | 0.13213 | 0.02089 | 0.19588 | 0.05364 | 0.04046 | 0.00061 | 0.05173 | 0.24897 | 0.04291 | 0.12036 |
| V.amurensi | V.rupestris | 0.10781 | 0.48988 | 0.26971 | 0.18539 | 0.00273 | 0.12502 | 0.13791 | 0.53087 | 0.60927 | 0.08017 | 0.27763 | 0.34811 | 0.03108 | 0.04182 | 0.01941 | 0.01213 | 0.00228 | 0.00441 | 0.03501 | 0.08794 | 0.31824 | 0.36206 | 0.04909 | 0.44305 |
| V.amurensi | V.vulpina | 0.02932 | 0.026001 | 0.13524 | 0.13766 | 0.22821 | 0.21099 | 0.07686 | 0.25417 | 0.21719 | 0.31646 | 0.00546 | 0.18921 | 0.3203 | 0.25136 | 0.22035 | 0.03234 | 0.03027 | 0.07227 | 0.04195 | 0.02539 | 0.01007 | 0.03284 | 0.11389 | 0.00557 |
| V.cinerea | V.coignetia | 0.17073 | 0.01636 | 0.01207 | 0.01121 | 0.22342 | 0.3288 | 0.40538 | 0.64113 | 0.73161 | 0.14065 | 0.10167 | 0.22292 | 0.20965 | 0.02855 | 0.0113 | 0.15276 | 0.18756 | 0.2278 | 0.03663 | 0.27817 | 0.49113 | 0.14507 | 0.14209 | 0.03074 |
| V.cinerea | V.labrusca | 0.15526 | 0.28754 | 0.07261 | 0.05206 | 0.07621 | 0.16075 | 0.13808 | 0.25991 | 0.30897 | 0.06143 | 0.25571 | 0.06769 | 0.00699 | 0.06432 | 0.0294 | 0.1002 | 0.22324 | 0.19443 | 0.1087 | 0.02053 | 0.12756 | 0.13651 | 0.1057 | 0.07899 |
| V.cinerea | V.palmata | 0.10027 | 0.07045 | 0.08459 | 0.04938 | 0.0206 | 0.08881 | 0.00362 | 0.00119 | 0.0432 | 0.14634 | 0.01458 | 0.01202 | 0.08498 | 0.20923 | 0.08449 | 0.11123 | 0.22793 | 0.16512 | 0.13999 | 0.1035 | 0.04687 | 0.39757 | 0.36824 | 0.24044 |
| V.cinerea | V.riparia | 0.43059 | 0.006343 | 0.32505 | 0.1093 | 0.09875 | 0.08945 | 0.03358 | 0.19472 | 0.2927 | 0.05022 | 0.02566 | 0.26918 | 0.33445 | 0.05389 | 0.01053 | 0.19583 | 0.12716 | 0.2394 | 0.13871 | 0.06685 | 0.18818 | 0.15633 | 0.20229 | 0.10408 |
| V.cinerea | V.rupestris | 0.14708 | 0.019655 | 0.07603 | 0.08033 | 0.13884 | 0.17106 | 0.10288 | 0.10099 | 0.01174 | 0.29081 | 0.06536 | 0.04524 | 0.06753 | 0.16626 | 0.08729 | 0.04994 | 0.18621 | 0.17605 | 0.18061 | 0.18519 | 0.19348 | 0.02835 | 0.32544 | 0.0413 |
| V.cinerea | V.vulpina | 0.3133 | 0.12389 | 0.02275 | 0.0245 | 0.03178 | 0.04173 | 0.0427 | 0.00825 | 0.0259 | 0.26062 | 0.00641 | 0.19276 | 0.1996 | 0.22516 | 0.20125 | 0.08984 | 0.2985 | 0.19757 | 0.1958 | 0.20383 | 0.19842 | 0.15184 | 0.03106 | 0.0358 |
| V.coignetia | V.labrusca | 0.05848 | 0.31473 | 0.16651 | 0.08058 | 0.00836 | 0.09328 | 0.18239 | 0.03663 | 0.03693 | 0.06173 | 0.17309 | 0.14622 | 0.10382 | 0.05433 | 0.0396 | 0.01901 | 0.03717 | 0.01379 | 0.00316 | 0.02117 | 0.17713 | 0.04456 | 0.03938 | 0.1305 |
| V.coignetia | V.palmata | 0.43635 | 0.12463 | 0.30289 | 0.45106 | 0.47894 | 0.44552 | 0.88612 | 1.2529 | 0.96105 | 0.95338 | 0.29554 | 0.49649 | 0.20195 | 0.2605 | 0.01749 | 0.10325 | 0.28637 | 0.3279 | 0.08832 | 0.11016 | 1.0248 | 1.0372 | 0.738 | 0.30641 |
| V.coignetia | V.riparia | 0.16657 | 0.25662 | 0.22139 | 0.08972 | 0.07294 | 0.21123 | 0.29896 | 0.46417 | 0.50617 | 0.04353 | 0.08147 | 0.3719 | 0.33525 | 0.10902 | 0.10854 | 0.05861 | 0.1862 | 0.03605 | 0.03513 | 0.13913 | 0.19464 | 0.30412 | 0.00445 | 0.41761 |
| V.coignetia | V.rupestris | 0.08513 | 0.033953 | 0.10598 | 0.12872 | 0.05337 | 0.01044 | 0.10236 | 0.39562 | 0.50108 | 0.03395 | 0.008 | 0.13024 | 0.0348 | 0.44436 | 0.32135 | 0.16014 | 0.02985 | 0.07192 | 0.17193 | 0.27266 | 0.08857 | 0.07585 | 0.36089 | 0.30896 |
| V.coignetia | V.vulpina | 0.17488 | 0.14622 | 0.23648 | 0.10444 | 0.14015 | 0.15416 | 0.23626 | 0.38175 | 0.51672 | 0.0383 | 0.20561 | 0.35732 | 0.20343 | 0.01669 | 0.006 | 0.03881 | 0.0739 | 0.08785 | 0.02444 | 0.14005 | 0.08522 | 0.12854 | 0.02176 | 0.11857 |
| V.labrusca | V.palmata | 0.04169 | 0.18722 | 0.19137 | 0.03321 | 0.14141 | 0.12675 | 0.10539 | 0.20363 | 0.27857 | 0.04994 | 0.14112 | 0.06067 | 0.01107 | 0.01914 | 0.03487 | 0.00116 | 0.03341 | 0.02499 | 0.01171 | 0.05012 | 0.007 | 0.15254 | 0.04978 | 0.10998 |
| V.labrusca | V.riparia | 0.01891 | 0.40859 | 0.20657 | 0.10837 | 0.08877 | 0.18905 | 0.29087 | 0.30851 | 0.28359 | 0.02049 | 0.22903 | 0.16131 | 0.13213 | 0.03384 | 0.01179 | 0.02843 | 0.09153 | 0.03253 | 0.00928 | 0.05673 | 0.06765 | 0.04395 | 0.06417 | 0.24374 |
| V.labrusca | V.rupestris | 0.05413 | 0.39159 | 0.15395 | 0.00702 | 0.00704 | 0.09042 | 0.12181 | 0.22387 | 0.28533 | 0.00279 | 0.24299 | 0.10087 | 0.05013 | 0.00661 | 0.01545 | 0.01755 | 0.02642 | 0.00854 | 0.02293 | 0.05848 | 0.09217 | 0.1174 | 0.01083 | 0.25375 |
| V.labrusca | V.vulpina | 0.08117 | 0.16923 | 0.13853 | 0.06808 | 0.19042 | 0.14588 | 0.03078 | 0.2877 | 0.37505 | 0.07047 | 0.18784 | 0.00798 | 0.17006 | 0.11219 | 0.12007 | 0.06595 | 0.03273 | 0.06442 | 0.00692 | 0.08277 | 0.01449 | 0.11186 | 0.06413 | 0.04246 |
| V.palmata | V.riparia | 0.11381 | 0.097598 | 0.44321 | 0.08208 | 0.38419 | 0.05722 | 0.17249 | 0.24099 | 0.30871 | 0.15634 | 0.02529 | 0.08061 | 0.22273 | 0.10572 | 0.14637 | 0.05253 | 0.04654 | 0.0292 | 0.02325 | 0.05477 | 0.14785 | 0.32504 | 0.13003 | 0.23351 |
| V.palmata | V.rupestris | 0.12028 | 0.017206 | 0.35102 | 0.1455 | 0.19988 | 0.08519 | 0.23768 | 0.07885 | 0.04597 | 0.01388 | 0.03339 | 0.24523 | 0.0673 | 0.01497 | 0.03583 | 0.07131 | 0.0711 | 0.05842 | 0.06507 | 0.07207 | 0.05812 | 0.43611 | 0.0869 | 0.3054 |
| V.palmata | V.vulpina | 0.08748 | 0.16207 | 0.18923 | 0.08516 | 0.04479 | 0.02346 | 0.25904 | 0.17535 | 0.10111 | 0.07358 | 0.06822 | 0.53106 | 0.24262 | 0.29148 | 0.20568 | 0.05447 | 0.04008 | 0.01659 | 0.01101 | 0.04172 | 0.17905 | 0.61174 | 0.19301 | 0.1891 |
| V.riparia | V.rupestris | 0.66867 | 0.076619 | 0.1966 | 0.10559 | 0.25588 | 0.28959 | 0.81452 | 0.22394 | 0.02503 | 0.17242 | 0.104 | 0.26605 | 0.38337 | 0.05761 | 0.04647 | 0.03047 | 0.25758 | 0.06817 | 0.08036 | 0.0781 | 0.07595 | 0.06791 | 0.01923 | 0.37408 |
| V.riparia | V.vulpina | 0.02575 | 0.19691 | 0.05532 | 0.08451 | 0.14643 | 0.00243 | 0.18636 | 0.0889 | 0.22242 | 0.16629 | 0.02225 | 0.49187 | 0.58744 | 0.2491 | 0.21729 | 0.08041 | 0.18482 | 0.01742 | 0.03687 | 0.06781 | 0.16462 | 0.357 | 0.07885 | 0.18407 |
| V.rupestris | V.vulpina | 0.06668 | 0.1027 | 0.16917 | 0.03908 | 0.14312 | 0.0794 | 0.05435 | 0.04662 | 0.12979 | 0.1728 | 0.14683 | 0.18456 | 0.15948 | 0.24451 | 0.18509 | 0.14387 | 0.06347 | 0.11228 | 0.08756 | 0.05253 | 0.08329 | 0.18666 | 0.13051 | 0.29354 |

**Table S4** Trait Pagel's lambda for phylogenetic analysis

| | lambda | pvalue |
|---|---|---|
| Volume | **0.8693595** | 6.77E-16 |
| ConvexHullVolume | **0.9958489** | 2.20E-22 |
| Solidity | 0.6673019 | 5.05E-10 |
| SurfaceArea | **0.9241048** | 1.26E-19 |
| Sphericity | **0.8197016** | 2.37E-20 |
| TotalBranchLength | **0.9721602** | 6.37E-24 |
| RachisLength | **0.99398** | 1.32E-18 |
| AvgBranchLength | 0.7653311 | 4.37E-12 |
| 2nd/LongestBranchLength | 0.0606601 | 0.341537 |
| BranchDiameter | 0.7420774 | 1.72E-06 |
| NumberOfPedicel | **0.9289866** | 2.79E-20 |
| PedicelLength | 0.3980271 | 0.000233 |
| PedicleDiameter | **0.8019726** | 7.28E-11 |
| PedicelAngle | 0.3421429 | 0.000391 |
| PedicelLength/RachisLength | 0.4242821 | 4.53E-07 |
| TotalBerryPotentialVolume | **0.9693417** | 8.91E-18 |
| AvgBerryPotentialDiameter | 0.6729516 | 4.20E-21 |
| BerryPotentialTouchingDensity | 0.2483078 | 4.96E-04 |
| PH_PC1 | **0.9999339** | 1.12E-21 |
| PH_PC2 | **0.9829545** | 1.08E-06 |
| PH_PC3 | 0.4414896 | 0.000314 |
| PHn_PC1 | **0.8707632** | 3.76E-22 |
| PHn_PC2 | 0.4427778 | 3.44E-06 |
| PHn_PC3 | 0.5169356 | 2.46E-08 |

**Table S5** Trait variation for each clade

| | Asian Clade | NA Clade I | NA Clade II |
|---|---|---|---|
| all traits | 0.38761 | 0.13645 | 0.64194 |
| PHn_PC3 | 0.044256 | 0.12313 | 0.5196 |
| AvgBerryPotentialDiameter | 0.66801 | 0.013841 | 0.79987 |
| PedicelBranchAngle | 0.040962 | 0.0073817 | 0.29219 |
| PedicelLength/RachisLength | 0.69688 | 0.045289 | 0.539 |
| Solidity | 0.30902 | 0.17599 | 0.59485 |
| Sphericity | 0.41354 | 0.17918 | 0.78967 |
| PHn_PC1 | 0.020012 | 0.52043 | 0.67961 |
| PedicleDiameter | 0.0073523 | 0.15227 | 1.186 |
| BranchDiameter | 0.14483 | 0.37703 | 0.75482 |
| PH_PC2 | 0.57456 | 0.015576 | 0.66602 |
| TotalBerryPotentialVolume | 0.18507 | 0.16057 | 2.0931 |
| PedicelLength | 0.076242 | 0.040299 | 0.28491 |
| AvgBranchLength | 0.19157 | 0.020708 | 0.31189 |
| PH_PC1 | 1.1903 | 0.021492 | 0.49693 |
| RachisLength | 1.1004 | 0.028016 | 0.54476 |
| ConvexHullVolume | 0.12859 | 0.021048 | 0.66791 |
| NumberOfPedicel | 0.030807 | 0.20208 | 0.99798 |
| TotalBranchLength | 0.084014 | 0.077846 | 0.91994 |
| SurfaceArea | 0.21317 | 0.16573 | 0.76814 |
| Volume | 0.38337 | 0.26725 | 0.63877 |
| PHn_PC2 | 3.36E-05 | 0.28041 | 0.19312 |
| 2nd/LongestBranchLength | 0.0026573 | 0.12072 | 0.35642 |
| PH_PC3 | 0.0067554 | 0.067675 | 0.19692 |
| BerryPotentialTouchingDensity | 0.023076 | 0.042071 | 0.091786 |

**Video S1** Illustration of quantifying branching topology using persistent homology.

**Video S2** Berry potential simulation.
